# Endothelial Cell Expression of STING^V154M^ Gain-of-Function Mutation Delays the Resolution of UVB-induced Skin Injury

**DOI:** 10.1101/2025.07.14.664621

**Authors:** Jane Chuprin, Carolina Salomāo Lopes, Kevin MingJie Gao, Kristy Chiang, Khashayar Afshari, Nazgol Haddadi, Manuel Garber, Milka Koupenova, Katherine A. Fitzgerald, Ann Marshak-Rothstein, Mehdi Rashighi

**Affiliations:** UMass Chan Medical School, Department of Dermatology, Worcester, MA 01605; UMass Chan Medical School, Department of Genomics and Computational Biology, Worcester, MA 01605; UMass Chan Medical School, Department of Medicine, Worcester, MA 01605

**Keywords:** SAVI, STING, endothelial cells, UVB injury, macrophage

## Abstract

Gain-of-function mutations in STimulator of INterferon Genes (STING) cause STING-Associated Vasculopathy with Onset in Infancy (SAVI), a rare autoinflammatory disease characterized by debilitating inflammatory lung disease and hallmark skin manifestations, such as chilblains and progressive, non-healing ulcers. Mice expressing the most common SAVI-associated variant STING^V154M^ (VM) recapitulate many clinical features of SAVI, including inflammatory lung disease, but do not develop spontaneous skin lesions. In this study, we show that a single low dose of ultraviolet B (UVB) irradiation, which induces only transient skin inflammation in wild-type (WT) mice, causes severe and progressive skin injury in VM mice. Notably, this phenotype persisted in VM mice depleted of hematopoietic cells and reconstituted with WT bone marrow, demonstrating that STING^V154M^ expression in non-hematopoietic cells is sufficient to drive persistent skin inflammation. Further analysis identified endothelial cells expressing STING^V154M^ as the primary driver of the cutaneous phenotype. Flow cytometry and bulk RNA sequencing showed that VM mice exhibited reduced early skin infiltration of macrophages and dendritic cells after UVB exposure. These findings establish a critical link between endothelial STING activation, impaired recruitment of skin myeloid cells, and defective resolution of acute inflammation, offering new insights into the pathogenesis of SAVI-associated skin disease.

## INTRODUCTION

The cyclic GMP-AMP synthase (**cGAS**)/ Stimulator of Interferon Genes (**STING**) pathway is a critical component of the innate immune system, primarily involved in sensing cytosolic DNA and subsequent production of type 1 interferons (**T1IFNs**) and pro-inflammatory cytokines such as IL-1f3, IL-6, and TNFa. STING activation helps orchestrate immune responses to viral infections, bacterial DNA, and other forms of cellular stress. However, aberrant activation of STING has been implicated in various immune-mediated disorders, including STING-associated vasculopathy with onset in infancy (**SAVI**) (Liu et al., 2014).

SAVI is a rare, monogenetic, autoinflammatory syndrome caused by a gain-of-function (**GOF**) mutation in the gene TMEM173, encoding STING protein, that leads to the persistent, cGAS-independent activation of STING. In addition to interstitial lung disease, SAVI patients typically develop vasculopathic skin lesions, manifesting as chilblains as well as progressive ulcers that can result in necrosis and autoamputation (Alehashemi et al., 2025, Frémond et al., 2021, Keskitalo et al., 2019, Liu et al., 2014, Melki et al., 2017, Sanchez et al., 2018, Valeri et al., 2024). Histologically, these lesions are characterized by immune infiltration, microthrombosis, and destruction of the skin capillaries and small vessels (Frémond et al., 2021, Liu et al., 2014). There are also case reports of impaired healing at vaccination sites, as well as non-healing wounds that required surgical intervention, immunosuppressive therapy, or the use of a negative pressure wound vac (Isoherranen et al., 2021, Keskitalo et al., 2019, Melki et al., 2017).

Mouse models of SAVI recapitulate the development of interstitial lung disease and have further revealed a key role for STING GOF-expressing endothelial cells in the recruitment of lymphocytes to the lung and the formation of bronchus-associated lymphoid tissue (BALT) (Gao et al., 2024, Gao et al., 2022, Luksch et al., 2019, Warner et al., 2017). However, SAVI mice do not develop spontaneous skin lesions, suggesting that additional environmental or inflammatory triggers may be required to unmask skin pathology (Motwani et al., 2019). This possibility is supported by studies showing that baseline immune activation or pre-existing inflammation can significantly modify STING-mediated tissue responses. For example, while STING activation in mechanical wound healing models can promote tissue repair (Mizutani et al., 2020), it can also exacerbate inflammation and impair wound healing in chronically inflamed skin, such as in diabetic wounds (Feng et al., 2022, Geng et al., 2023). Moreover, pre-existing immune activation has been shown to amplify STING-driven inflammation in settings such as WT mice pre-treated with poly(I:C) and mouse models of psoriasis driven by STING agonists (Pan et al., 2021, Pyclik et al., 2023, Sun et al., 2024, Yu et al., 2022).

To explore this potential synergy in the context of SAVI skin disease, we used acute, low-dose UVB radiation as a trigger, given that UVB is a known activator of the STING pathway (Skopelja-Gardner et al., 2020, Sontheimer et al., 2017). We now show that expression of STING^V154M^ in endothelial cells drives exacerbated and prolonged skin injury following UVB exposure. These studies have important implications for the management of SAVI patients and potentially other conditions that impact the wound healing process.

## RESULTS AND DISCUSSION

### STING^V154M^ causes exacerbated and prolonged skin injury in response to UVB irradiation

To investigate the impact of STING GOF on the skin’s response to injury, we used gene-targeted STING^V154M/WT^ (**VM**) mice, which harbor the most common mutation observed in SAVI patients (Frémond et al., 2021, Gao et al., 2022). Unlike previous UVB studies (Skopelja-Gardner et al., 2020, Sontheimer et al., 2017), our mice were given a single low dose of UVB irradiation to limit both the extent and timing of the injury. Fur was removed from the dorsal skin of VM and WT mice and 24 hours later, the treated areas were exposed to broad-band UVB irradiation (100 mJ/cm²). Mice were monitored for up to 21 days using a clinical scoring system based on the extent and severity of the lesions (see Methods).

At baseline and following fur removal, there were no notable clinical or histological differences between WT and VM mice (**Supplementary** Figure 1). However, by day 3 post-UVB irradiation, VM mice displayed prominent erythematous patches and crusting, while WT mice exhibited only mild erythema and fine scaling. Over the following few days, the condition of VM mice worsened, progressing to skin erosions and, in most cases, full-thickness ulceration that persisted through day 21. In contrast, WT mice never developed ulcerations and were nearly completely healed by day 7 (**Figure 1a-b**). These results demonstrate that STING GOF significantly exacerbates and prolongs UVB-induced skin injury, leading to chronic skin injury. This outcome further suggests that hyperinflammatory responses and/or an inability to appropriately heal relatively mild skin injuries may also account for the skin phenotype of SAVI patients.

**Figure 1.**
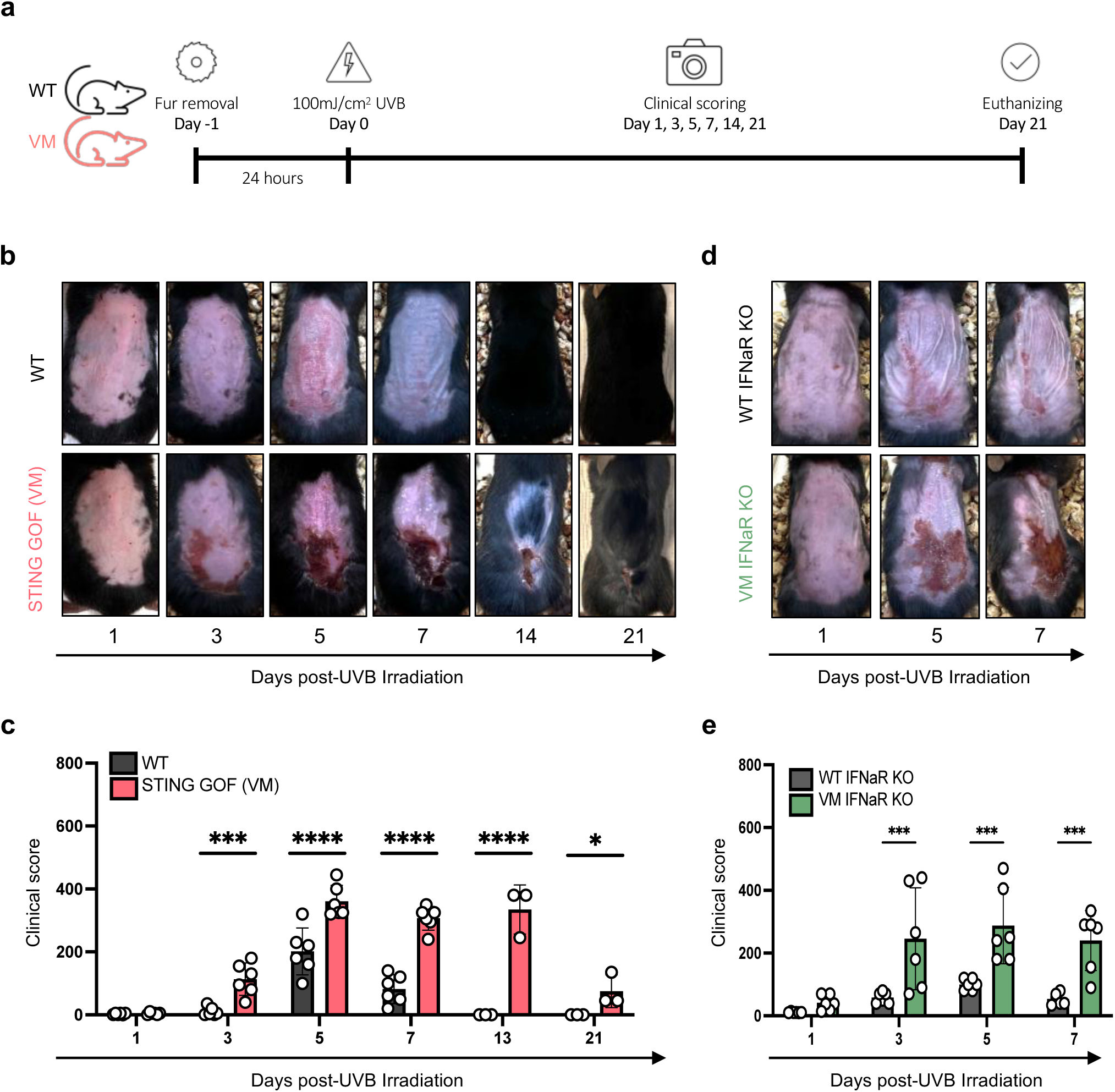
STING Gain of function leads to exacerbated inflammatory response following UVB irradiation. (a) Schematic of experimental setup; (b) Representative images of UVB-irradiated 6-10 wk old mice at indicated time points, WT (n=6), VM (n=6) and up to 21 days WT (n=3), VM (n=3); (c) Clinical scores of WT vs VM mice; (d) Representative images of UVB treated 6-10-week-old WT IFNaR KO (n=6), VM x IFNaR KO (n=6); (e) Clinical scores for IFNαR mice.

### Type 1 Interferons are not required for STING-driven exacerbated skin responses

SAVI is often classified as a “type I interferonopathy,” where inflammation is attributed to IRF3-mediated induction of type I interferons (T1IFNs) and subsequent activation of IFN-stimulated genes. However, previous studies have shown that lung pathology in SAVI mouse models does not require T1IFN signaling [7, 9]. To determine whether the exacerbated skin response to UVB irradiation in VM mice is dependent on T1IFNs, we intercrossed VM mice with type I interferon receptor knockout (IFNαR^-/-^) mice to generate STING^V154M/WT^ x IFNαR^-/-^ mice. These mice exhibited comparably severe and persistent UVB-induced skin lesions as the IFNαR-sufficient VM mice (**Figure 1c-d**). Thus, similar to their lung pathology, the exacerbated UVB-induced skin injury in VM mice occurs independently of type I IFN signaling (Gao et al., 2022, Luksch et al., 2019).

### Non-hematopoietic cells expressing STING GOF drive exacerbated skin disease

To better understand how the VM mutant drives an exacerbated response to UVB, it was necessary to identify the cell type(s) directly responsible for the hyperinflammatory response. This question was first addressed by using chimeric mice in which the hematopoietic compartment of VM mice was replaced by stem cells from WT mice. To avoid any confounding effects of gamma irradiation damage on the skin, we used busulfan-based myeloablation, a chemotherapy approach previously described for chimera production (Gao et al., 2022, Montecino-Rodriguez and Dorshkind, 2020). CD45.2 recipient mice (WT or VM) were pretreated with busulfan for two days and then reconstituted with CD45.1 WT donor bone marrow cells **(Figure 2a)**. Two months later, flow cytometric analysis of bone marrow, spleen, and skin confirmed that ∼95% of the hematopoietic cells in WT → VM chimeras were donor-derived, indicating successful engraftment (**Figure 2b-c**). By contrast, WT → WT chimeras showed only ∼50% donor reconstitution, likely due to the lower efficacy of busulfan in depletion of WT hematopoietic cells. The enhanced busulfan sensitivity of VM stem cells or hematopoietic cells is likely due to an amplified ER stress resulting from chronic STING activation (Wu et al., 2019). Despite these differences, we successfully generated two groups of mice with near-complete WT hematopoietic systems. Following UVB exposure, WT → VM chimeras exhibited significantly more severe skin injury than the control WT → WT chimeras (**Figure 2d**), mirroring the skin pathology observed in VM mice (**Figure 1a-b**). These findings indicate that STING GOF in non-hematopoietic cells is sufficient to drive the exacerbated UVB-induced skin pathology observed in VM mice.

**Figure 2.**
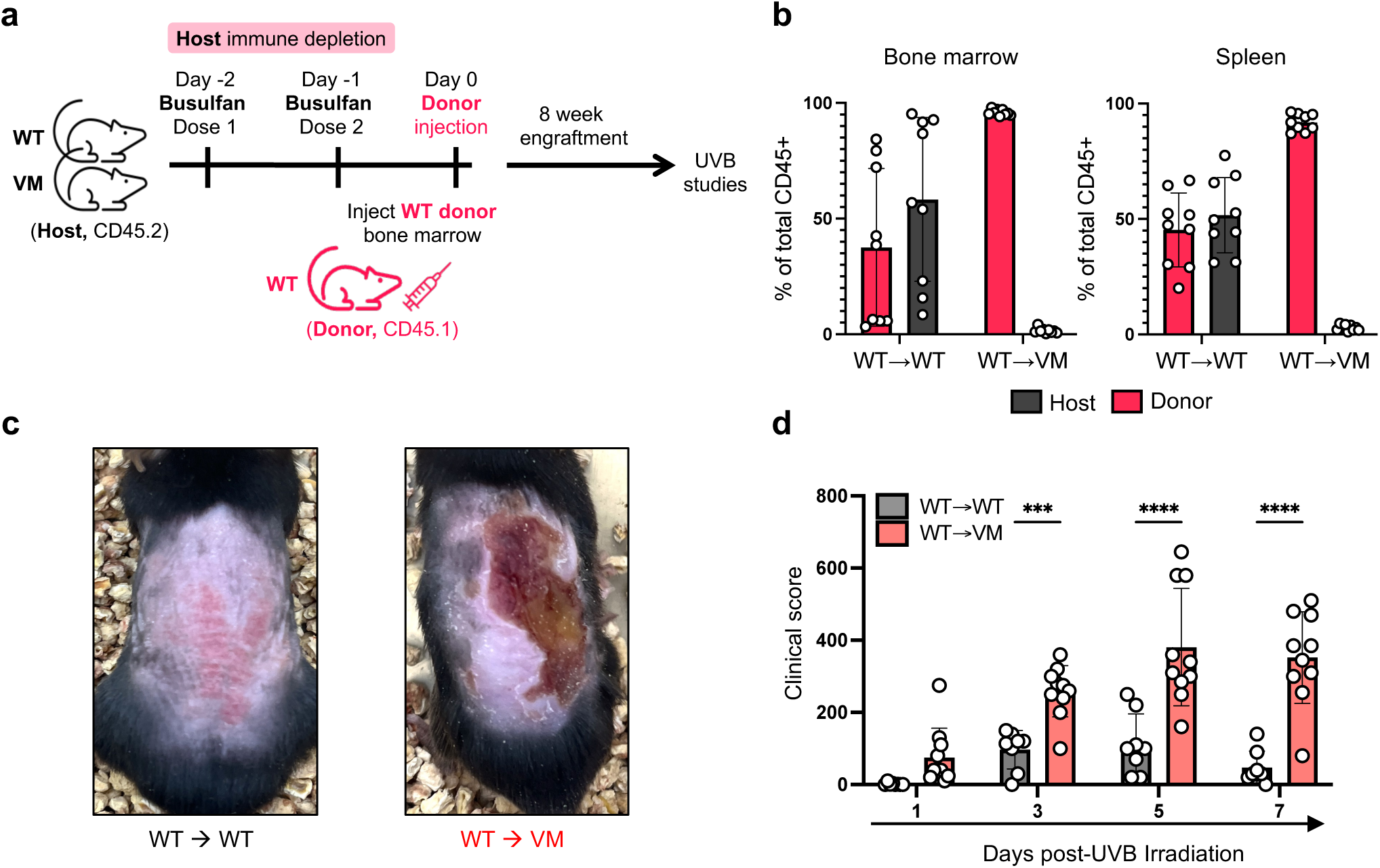
Exacerbated UVB-induced injury depends on VM-expressing radioresistant cells. (a) Schematic of experimental setup; (b) Extent of donor and host stem cell engraftment in the bone marrow and spleen of WT➔WT (n=9) and WT➔VM (n=9) chimeric mice; (c) Representative image of skin damage 5 days post UVB; (d) Clinical scores for skin damage on days 1-7 post-UVB in WT→WT (n=8) and WT→VM (n=10) chimeric mice.

Radiation chimeras have been used previously to assess the importance of hematopoietic versus radioresistant cells on the development of autoinflammation in SAVI mouse models (Gao et al., 2024). Mice that express the N153S mutation spontaneously develop an inflammatory bowel disease (**IBD**), and NS IBD depends on NS-expressing myeloid cells, as shown by the development of IBD in lethally irradiated WT mice reconstituted with Rag1^-/-^ NS bone marrow stem cells (Rag1-/-NS➔WT chimeras) (Shmuel-Galia et al., 2021). By contrast, VM mice spontaneously develop an inflammatory lung disease (**ILD**) associated with BALT formation, which, similar to the skin phenotype, depends on VM expression in radioresistant cells (Gao et al., 2022).

### Endothelial cells expressing STING GOF are the main drivers of exacerbated skin injury

To identify the specific non-hematopoietic cell type responsible for the exacerbated UVB-induced skin injury, we used a previously described VM-conditional knock-in (**cKI**) mouse line (Gao et al., 2024). Gene targeting of the endogenous STING locus inserted a floxed STOP codon into the intron preceding the exon encoding the VM mutation, and thus, Cre-mediated deletion of the STOP codon resulted in tissue-specific expression of the mutant allele under the control of the normal STING promoter.

From publicly available protein and single-cell RNAseq expression data of human skin we identified endothelial cells, fibroblasts, and keratinocytes as the top three STING expressing non-hematopoietic cell types (Uhlen M, 2023). The role of these cell types in the UVB response of VM mice was evaluated by using Cre-recombinase-expressing mouse strains specific for each lineage. Expression of the VM mutation in either keratinocytes [K14-Cre x cKI] or fibroblasts (Fsp1-Cre x cKI), failed to replicate the exacerbated skin injury of VM mice (**Figure 3a-b**). However, expression of the VM mutation in endothelial cells [Tie2-Cre x cKI] resulted in exacerbated UVB-induced skin injury (**Figure 3a-b**). Although the Tie2 promoter is widely used to target endothelial cells, it is also active in hematopoietic progenitor cells, which can lead to expression in CD45+ hematopoietic lineages (Tang et al., 2010).

**Figure 3:**
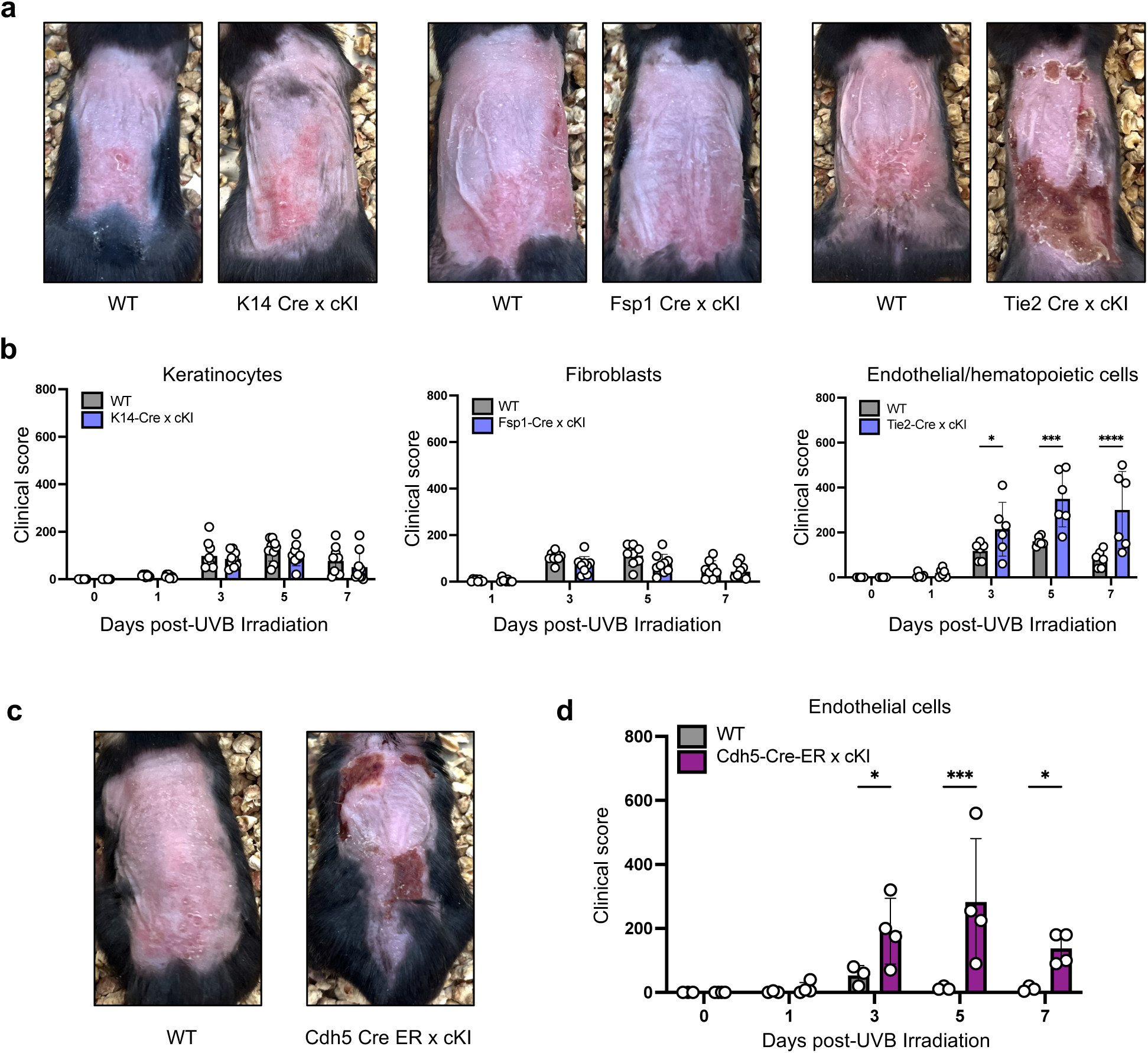
VM-expressing endothelial cells are sufficient for UVB-induced skin injury. (a) 11-15 week old male and females K14-Cre x cKI (n=9) or littermate WT controls (n=9), 10-16 week old male and female Fsp1-Cre x cKI (n=9) or littermate WT (n=8), and 7-8 week old male and females Tie2-Cre x cKI (n=6) or littermate WT (n=6) were followed for 7 days post-UVB; (b) Clinical scores of images in a; (c) 13 week old female CAGG-Cre-ER x cKI (n=2) or littermate WT (n=5), and 11-18 week old male and female Cdh5-Cre-ER x cKI (n=4) or littermate WT (n=3) were followed for 7 days (d) Clinical scores of images in c.

To achieve more specific targeting of endothelial cells, we utilized a tamoxifen-inducible Cdh5-Cre-ER x cKI system where postnatal administration of tamoxifen induced VM expression in endothelial cells and not hematopoietic cells (Gao et al., 2024). These mice were compared to CAGG-Cre-ER x cKI mice, where tamoxifen induced ubiquitous VM expression. Importantly, Cdh5-Cre-ER x cKI mice exhibited exacerbated UVB-induced skin injury (**Figure 3c-d**), confirming that endothelial cells are key mediators of the observed skin phenotype. The severity of skin injury in the Cdh5-Cre-ER x cKI model was somewhat milder than CAGG-Cre-ER x cKI VM mice, indicating that additional VM-expressing cell types synergize with endothelial cells to promote VM skin disease. This finding was consistent with prior studies in which overexpression of STING in endothelial cells, due to either the VM mutation or excessive cGAS induction, led to the development of interstitial lung disease associated with the formation of tertiary lymphoid structures. In both cases, lung disease was exacerbated by synergistic interactions with other cell types (Gao et al., 2024).

The robust skin injury phenotype observed following a single low-dose UVB exposure in VM mice contrasts with the lack of clear UVB photosensitivity in patients with SAVI. This discrepancy may be attributed to inherent structural differences between mouse and human skin, particularly in terms of epidermal thickness (Zomer and Trentin, 2018). The depth of UV penetration into the skin depends on its wavelength, with UVB being largely absorbed by the thicker human epidermis, thereby limiting its direct impact on the underlying dermal endothelial cells (Meinhardt et al., 2008). Conversely, the thinner mouse epidermis allows greater UVB penetration, potentially leading to direct activation of dermal endothelial cells. In SAVI patients, the primary environmental trigger for skin manifestations appears to be cold exposure, leading to characteristic chilblains on acral areas. Mice, however, possess different thermoregulatory mechanisms, making cold exposure an ineffective trigger in VM mice (Gordon, 2017). Whether endothelial cells in VM mice are directly damaged by UVB exposure or respond to an inflamed environment will be an important question for future studies.

### Reduced numbers of skin resident macrophages and dendritic cells correlate with dysregulated immune responses and exacerbated skin injury in VM mice

To investigate the immune mechanisms underlying the exacerbated UVB-induced skin inflammation in VM mice, immune cell subsets present in the skin at baseline and following UVB irradiation were analyzed by flow cytometry at various times post UVB exposure. For baseline immune status of skin, mice were euthanized, their backs were shaved, and then the skin was digested and analyzed. VM mice had significantly fewer macrophages and dendritic cells at baseline than WT mice, with reductions of 50% and 85%, respectively, suggesting a deficiency in tissue-resident immune cells (**Figure 4a**).

**Figure 4:**
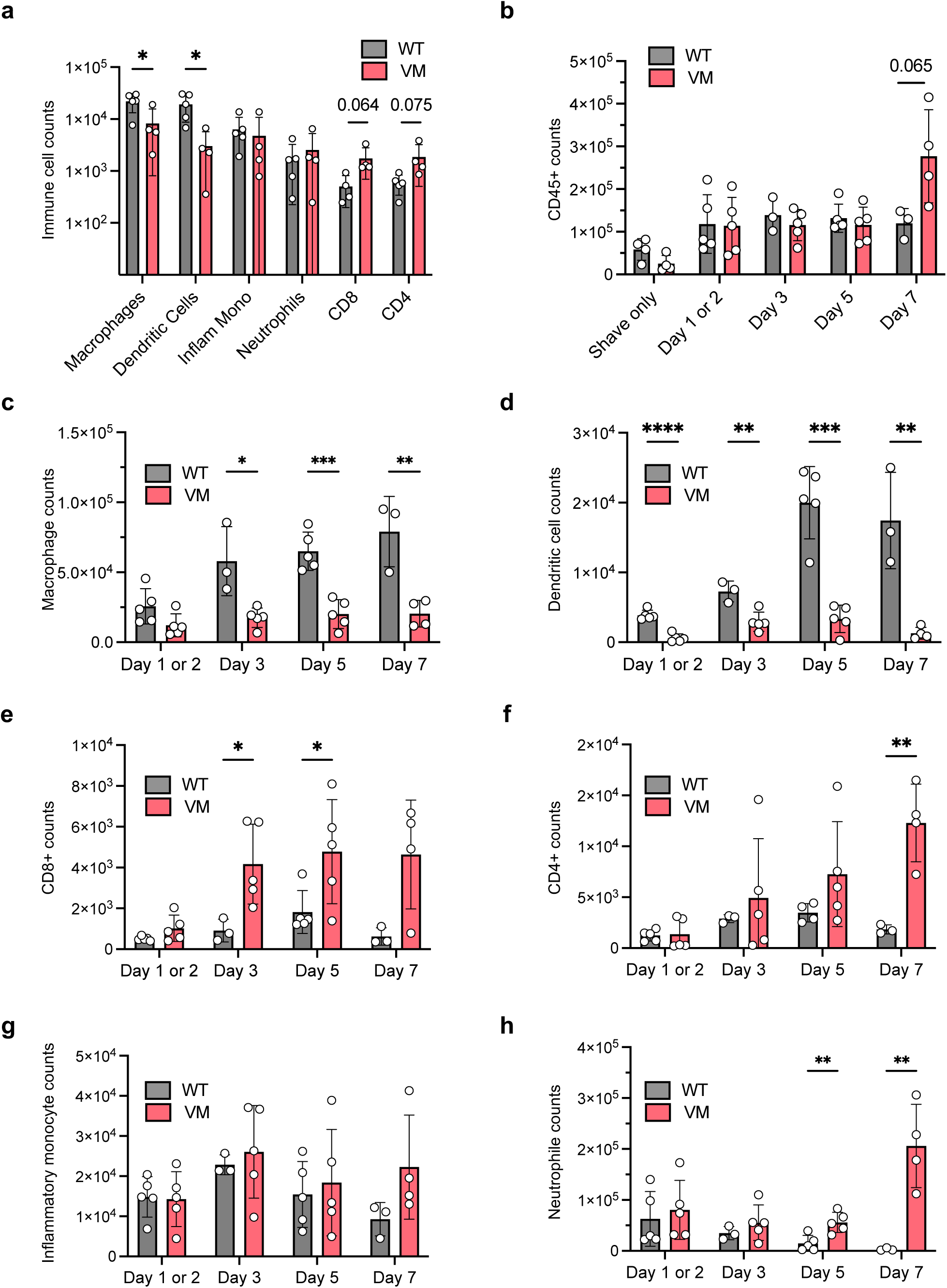
Flow cytometry reveals reduced macrophage and dendritic cell populations in VM skin. (a) Cell suspensions prepared from the skin of untreated euthanized WT (n=5) and VM (n=4) mice were analyzed by flow cytometry and the total number of CD45+ cells was determined; (b-h) Cell suspensions prepared from the skin at the indicated times post UVB-irradiation were analyzed by flow cytometry. Cell types were identified using the markers specified in Supplementary Figure 3. Each dot represents an individual mouse.

Despite similar total numbers of CD45+ cells in WT and VM mice during the first 5 days after UVB irradiation, their cellular composition differed between the two groups (**Figure 4b)**. Strikingly, unlike WT mice, macrophage accumulation was impaired in VM skin, particularly from 3 days post-UVB irradiation onward (**Figure 4c**), coinciding with the first signs of exacerbated skin inflammation (**Figure 1a**). Instead, VM skin contained more T cells than WT skin (**Figure 4e-f**). However, since VM T cell-deficient mice still developed severe skin disease following UVB irradiation (**Supplementary** Figure 2), a direct pathogenic role for T cells in the development of skin lesions appears unlikely.

By day 7, VM mice exhibited a dramatic surge in neutrophil numbers, reflecting a failure to resolve the injury (**Figure 4g**). In contrast, neutrophil levels in WT mice declined to baseline, consistent with successful inflammation resolution. Unlike F480+ macrophages, Ly6C+ inflammatory monocyte numbers remained comparable between VM and WT mice at all analyzed time points (**Figure 4h**), suggesting that the lack of macrophages in VM skin was due to a reduction in tissue resident cells and not reduced recruitment of monocyte precursors. Additionally, dendritic cell numbers remained consistently lower in VM mice relative to WT throughout this timeframe (**Figure 4d**), indicating a persistent deficit in antigen-presenting cell populations. Together, these findings suggest that the exacerbated skin injury in VM mice is not due to massive immune cells infiltration but rather a failure to recruit or expand the macrophages required for effective tissue repair.

### RNA sequencing reveals extensive disruption in the expression of genes associated with immune activation, coagulation, and tissue repair in VM mice

To assess the transcriptional impact of UVB exposure in the SAVI model, we performed bulk RNA sequencing of dorsal skin samples from WT and VM mice collected at 6 hours and 3 days after a single UVB dose, as well as from unirradiated controls (noUVB_6h and noUVB_3d) (**Figure 5a**). Hierarchical clustering of differentially expressed genes revealed four major gene clusters (Clusters 1–4) with distinct temporal and genotypic expression patterns (**Figure 5b**). We had assumed that the 24-hour interval between fur removal and UVB exposure would allow the mice to recover from any minor irritation caused by shaving and depilation. However, the 6-hr no-UVB data revealed increased expression of numerous immune genes in both WT and VM skin, which eventually reduced to baseline in the 3d no-UVB groups (**Figure 5b, Clusters 1 and 2**). In contrast, VM skin exhibited prolonged and exaggerated transcriptional changes in response to UVB, with sustained upregulation of genes, particularly in Cluster 1 and to a lesser degree in Cluster 2 (**Figure 5b**).

**Figure 5:**
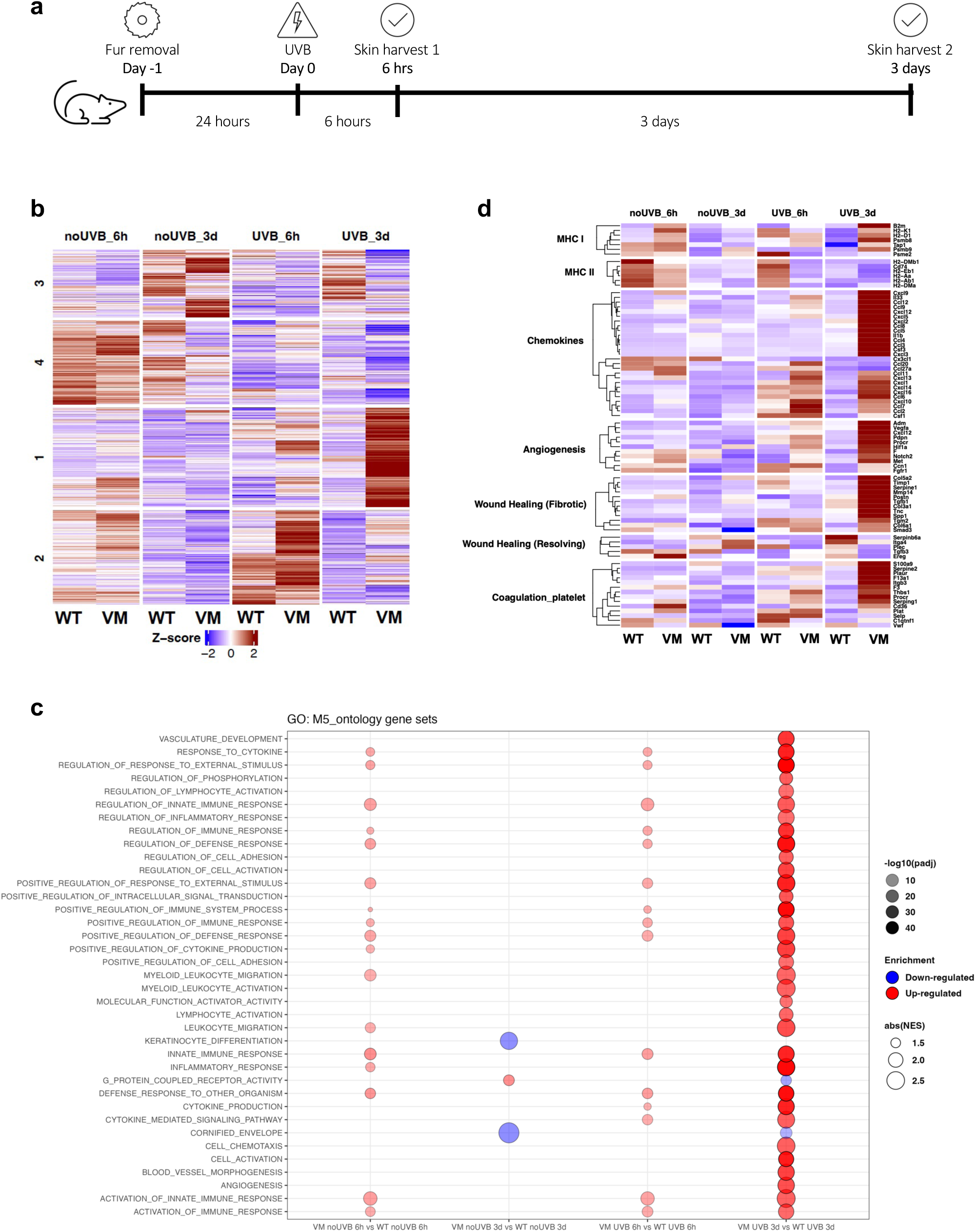
Bulk RNA-seq reveals differentially expressed genes between VM and WT skin at 6 hours and 3 days post-UVB. (a) schematic of experimental design; (b) Condensed heatmap of 1369 DE genes based on the following number of mice: no UVB 6hr WT (n=7), VM (n=7); no UVB 3dy WT (n=4), VM (n=3); UVB 6hrs WT (n=5), VM (n=7); UVB 3 days WT (n=5,) VM (n=6); (c) Gene ontology (GO) analysis; (d) Selected heatmap pathways.

Overall, the data identified dramatic differences between WT and VM mice both in response to the modest effects of fur removal and the more extensive injury induced by low-dose UVB irradiation. To define the biological processes underlying the transcriptional dysregulation observed in SAVI skin, we performed Gene Ontology (GO) enrichment analysis on differentially expressed genes across genotypes and time points (**Figure 5c**). Compared to WT, VM skin at baseline (noUVB_6h) already exhibited modest upregulation of pathways related to interferon signaling, inflammatory response, and cytokine production. These differences became more pronounced upon UVB exposure, particularly at 3 days, where VM skin showed marked enrichment of GO categories, including activation of immune response, leukocyte activation, keratinocyte differentiation, and cytokine-mediated signaling (**Figure 5c**). Notably, the VM vs WT UVB_3d comparison yielded the most extensive pathway enrichment, dominated by pro-inflammatory and tissue remodeling signatures, consistent with a non-resolving immune response.

To contextualize these global enrichment results, we next examined a panel of manually curated genes involved in MHC signaling, chemokine production, angiogenesis, and wound repair. These genes displayed a genotype- and UVB-dependent pattern of dysregulation, highlighting key drivers of the SAVI inflammatory phenotype (**Figure 5d**). The early WT response to both fur removal alone and UVB exposure included upregulation of MHC class II genes (eg. *H2-Aa, Cd74, H2-Eb1, H2-Ab1, and H2-DM*a) (**Figure 5d)**. This likely reflects the greater presence of classical antigen-presenting cells, including macrophages and DCs, in WT skin at baseline, which was previously observed by flow cytometry (**Figure 4a**). By contrast, VM mice showed more pronounced upregulation of MHC class I-associated genes (eg. *B2m, H2-K1, and Psmb9*). UVB-irradiated VM mice also exhibited distinct temporal waves of chemokine gene expression, including an early phase indicative of inflammatory monocyte and lymphocyte recruitment (eg*. Ccl2, Ccl20, Ccl7, Cxcl10)* followed by a massive wave of chemokines associated with macrophage and neutrophil recruitment and activation (eg. Ccl9, Cxcl2, Cxcl5) (**Figure 5d**). However, this dramatic increase in chemokine expression did not correlate with the relatively low numbers of monocytes and macrophages detected in VM skin by flow cytometry, suggesting a defect in leukocyte extravasation, potentially due to endothelial cell damage and dysregulated intravascular thrombosis.

This hypothesis is supported by upregulation of genes involved in platelet aggregation, clot formation and coagulation (e.g., *F3, F13a1, Cd36, Itgb3, Plaur, S100a9, Serpine2, Procr*), along with dysregulation of genes essential for effective clot resolution (*Serpine 1*). Furthermore, maintenance of an intact endothelial barrier relies on coordinated interaction between platelets and endothelial cells, including adhesion molecules such as *vWF*, which mediate platelet adhesion and aggregation at sites of injury. Notably, *vWF* and related genes are significantly downregulated in VM skin, further supporting the notion that impaired endothelial function contributes to defective leukocyte extravasation.

Endothelial cells, platelets, and coagulation are critical regulators of wound healing and angiogenesis. Consistently, VM mice displayed altered expression patterns of wound healing-related genes (eg. *Tgfb1, Pdgfb, Vegfa, Fn1*) at both 6 hours and 3 days post-UVB exposure compared to WT mice. VM mice also exhibited markedly elevated levels of pro-fibrotic genes, reflecting an exaggerated wound healing response. Markers of angiogenesis were significantly upregulated in VM mice, with early induction of *Hif1a*, *Adm, Ccn1, Notch2,* and *Pdpn*, and later expanding to include *Cxcl12, Vegfa, Fgfr1,* and *Procr.* Fibrotic remodeling was particularly pronounced by day 3, with increased expression of collagen genes (eg. *Col5a2, Col3a1, Col6a1*), matrix components (*Postn, Tnc, Spp1*), key wound healing regulators (*Tgfb1, Smad3*), matrix turnover genes (*Mmp14, Serpine1, Timp1*), and crosslinking enzymes (*Tgm2*). In contrast, WT mice upregulated genes associated with epithelial stabilization (*Plec*), regulation of immune cell migration (*Itga4*), tissue remodeling (*Capn3*), protection from protease-driven damage (*Serpinb6a*), and tissue regeneration signaling (*Ereg*), reflecting a shift towards tissue repair and barrier restoration. Together, these data highlight significant differences in the inflammatory regulation and wound healing responses to UVB between WT and VM mice. Many of the upregulated genes in WT mice were associated with tissue repair, while the marked gene upregulation in VM mice reflected a sustained inflammatory response.

Many of the genes upregulated in VM skin 3 days after UVB exposure in VM, but not in WT, belong to the category of IFN-stimulated genes (ISGs). This IFN signature included type I ISGs such as *Ifit3*, *Isg15*, *Oasl1*, *Rsad2*, *Stat1*, and *Irf7*, as well as type II ISGs including *Stat1, Irf1*, *Cxcl9/10, Psmb8/9, Tap1,* and *Ifng1*. Although type I IFNs are unlikely to be the primary drivers of skin pathology in this model, since IFNaR-/- VM mice still developed severe skin disease, the robust ISG expression may in part reflect increased production of IFN-γ by T cells or NK cells.

In summary, our findings demonstrate that the STING gain-of-function (GOF) VM mutation markedly impairs the cutaneous response to UVB-induced injury, resulting in persistent, non-healing ulcerative lesions. Although SAVI is classified as a type I interferonopathy, the exaggerated inflammatory response observed in VM mice occurs independently of type I IFN signaling. We further show that STING GOF drives this aberrant inflammatory response by disrupting normal myeloid cell trafficking and expansion. Notably, the VM mutation induces heightened expression of MHC class I genes, robust chemokine production, and excessive platelet activation, clot formation, and coagulation. In contrast, WT mice exposed to the same UVB insult exhibit a transcriptional program consistent with wound repair. Collectively, these results highlight endothelial STING as a key driver of skin pathology in SAVI and suggest that targeting STING-expressing endothelial cells may represent a novel therapeutic strategy.

## MATERIALS AND METHODS

### Mice/genotyping

STING^V154M^ (VM) mice and VM conditional knock-in (CKI) mice have been previously described (Gao et al., 2024, Motwani et al., 2019). IFNaR KO mice and were originally provided by Dr. J. Sprent (Scripps, Ma Jolla, CA)) or purchased from Jackson Labs (Kolumam et al., 2005). TCR-deficient TCRb^-/-^ TCRd^-/-^ B6 female mice were purchased from Jackson Labs and then backcrossed to VM. STING V154M conditional knock-in mice (cKI) were made by Drs. M. Shlomchik and S. Gingras at the University of Pittsburg, PA (Villar-Pazos et al., 2023). Mouse genotyping was done by Transnetyx, using real-time PCR and validated probes for each strain as described previously (Gao et al., 2024). Both male and female mice were used in experiments as there were no sex-specific phenotypic differences. All mouse procedures were approved by the Institutional Animal Care and Use Committee and conducted in accordance with relevant ethical guidelines.

### UVB Irradiation

Mice were anesthetized with 20 *µ*g/mL ketamine and 2 *µ*g/*µ*L xylazine, and fur was removed with electric pet clippers and 2 min treatment with Nair Sensitive Face Cream (hair depilatory cream). Anesthetized mice received one dose of 100mJ/cm^2^ 24 hours after fur removal using the Daavlin Irradiator with broad band UVB (290-320 nm) set to emit ∼2.3mJ/cm2/s.

### Clinical Score

Clinical scores were calculated as: (affected surface area, 0–100%) × (Severity score, 0=No lesion, 1=Accidental shaving nicks, 2=Mild erythema/scaling, 4=Severe erythema/scaling, 7=Erosion, and 9=Ulceration). Lesions of differing severity within the same mouse were scored separately and combined. Scores were determined blindly by a board-certified dermatologist.

### Flow Cytometry

Telogen-phase skin was collected and weighed. The skin was finely chopped and digested 30mins at 37C in digestion cocktail (0.1mg/mL DNAse I, 0.5mg/mL hyaluronidase, 2mg/mL collagenase IV in RPMI media). Gating strategy and antibodies found in supplementary materials.

### Chimera

Mice were lymphodepleted with 40 mg/kg busulfan (Caymen #14843) per protocol as previously described (Montecino-Rodriguez and Dorshkind, 2020).

### Bulk RNA Sequencing and analysis

Total RNA was extracted from frozen samples preserved in RLT buffer using the RNeasy Mini Plus Kit (QIAGEN). To remove residual genomic DNA, samples were treated with RNase-free DNase I for 15 minutes at room temperature. RNA-seq libraries were generated using an optimized CellSeq2 protocol for bulk RNA-seq per previous protocol (Gellatly et al., 2021). Libraries were prepared in pools of 24 samples, with 50 ng of input RNA per sample. Sequencing was performed on the NovaSeq 6000 platform. Raw FASTQ files were processed using a publicly available pipeline (https://github.com/Carol-Salomao/skin_cellseq2_bulk_pipeline), aligning to the mm10 genome to produce count matrices. Differential expression analysis was conducted using DESeq2, and gene set enrichment analysis (GSEA) was applied to identify significantly enriched pathways. All analyses were performed in R (version 4.3.3).

### Statistical Analysis

Statistics used for Clinical Scoring and Flow Cytometry analysis were analyzed using Prism9 by un-paired parametric t-tests between control and test groups, indicating p<0.05 * <0.05, **<0.01, ***<0.001, ****<0.0001.

## DATA AVAILABILITY STATEMENT

The RNA-sequencing data generated in this study have been deposited in the NCBI Gene Expression Omnibus (GEO) under the accession number PRJNA1273459. The dataset can be accessed directly at the following link: https://dataview.ncbi.nlm.nih.gov/object/PRJNA1273459?reviewer=o2nu0mb0gq217f8ul5e9l1p0lv

## ACKNOWLEDGEMENTS

Grant support: 5T32AI132152 (Chuprin, Gao, Autoimmunity and Autoinflammation Training Grant); T32GM107000 (Chuprin, Gao, MSTP at UMass Chan); 1R01HL165787-01 (Fitzgerald, Marshak-Rothstein MPI); 1R01HL153235 (Koupenova); 1R21AG093333 (Koupenova).

HL165787-02 and AI178978 Fitzgerald. The UVB irradiator was purchased with grant funds from the Lupus Research Alliance and kindly made available by Dr. J. Richmond UMass).

## SUPPLEMENTARY MATERIALS

**Supplementary Figure 1. VM skin exhibits impaired wound healing compared to WT following UVB injury.** Representative H&E-stained sections of dorsal skin from WT and VM mice at baseline and following UVB exposure. At baseline and early time points (day 2 after fur removal and day 3 post-UVB), no major histological differences were observed between WT and VM mice. By day 7 post-UVB, WT skin showed near-complete resolution of inflammation, with a normal epidermal architecture, preserved adnexal structures (including hair follicles), and minimal dermal inflammatory infiltrate. In contrast, VM skin demonstrated signs of impaired resolution, including epidermal hyperplasia (acanthosis), loss of adnexal structures, and prominent dermal immune cell infiltration.

**Supplementary Figure 2. T cells are not required for exacerbated UVB-induced injury in VM mice.** (a) Representative images of UVB-irradiated TCRb^-/-^ TCRd^-/-^ (TCR double-knockout, DKO) mice at indicated time points. WT TCR DKO mice (top row, n=2) and VM TCR DKO (bottom row, n=2) were monitored for UVB-induced skin injury over 7 days; (b) Clinical scores for skin damage from days 1 to day 7 post-UVB.

**Supplementary Figure 3. Flow cytometry gating strategy for identification of immune cell subsets infiltrating mouse skin.** Single, live cells were gated based on forward and side scatter (FSC-A/SSC-A) and a live/dead viability dye, followed by selection of CD45 hematopoietic cells. TCRβ T cells, CD11b myeloid cells, NK1.1 NK cells, and TCRγδ cells were gated in subsequent steps. Among TCRβ cells, CD4 and CD8 T cell subsets were identified. Within CD11b cells, neutrophils (Ly6G), inflammatory monocytes (Ly6C^hi^Ly6G), and Ly6C myeloid cells were further distinguished. Dendritic cells (CD11c MHC II) and macrophages (F4/80 MHC II) were resolved as separate populations within the CD11b compartment.

**Supplementary Figure 4. A comprehensive list of differentially expressed genes identified by bulk RNAseq at 6 hours and 3 days post UVB irradiation**. A total of 1,369 differentially expressed genes (DEGs) were identified across the following experimental groups (Fig. 5a): no UVB 6hr WT (n=7), VM (n=7); no UVB 3dy WT (n=4), VM (n=3); UVB 6hrs WT (n=5), VM (n=7); UVB 3 days WT (n=5,) VM (n=6).

**Supplementary Table 1. Antibodies used for flow cytometry analysis.** List of fluorochrome-conjugated antibodies used for flow cytometric analysis of mouse skin infiltrating immune cells. The table includes fluorophore, target antigen, dilution, clone, and vendor.

## AUTHOR CONTRIBUTION STATEMENT

Conceptualization: JC, KG, AMR, MR; Investigation: JC, CL, KA, NH; Formal analysis: JC, CL, KA Methodology; JC, KG, CL, AMR, MR; Data curation: JC, CL; Resources: KG, KC, MG, KF, AMR, MR; Supervision: AMR, MR; Visualization: JC, CL; Funding: AMR, MR Writing: JC, AMR, MR

## Abbreviations

STING: Gain-of-function mutations in STimulator of INterferon Genes
SAVI: STING-Associated Vasculopathy with Onset in Infancy
cKI: conditional knock-in
GOF: gain-of-function
WT: wild type

