## Supplemental Figure 1 for "Endothelial Cell Expression of STING^V154M^ Gain-of-Function Mutation Delays the Resolution of UVB-induced Skin Injury"

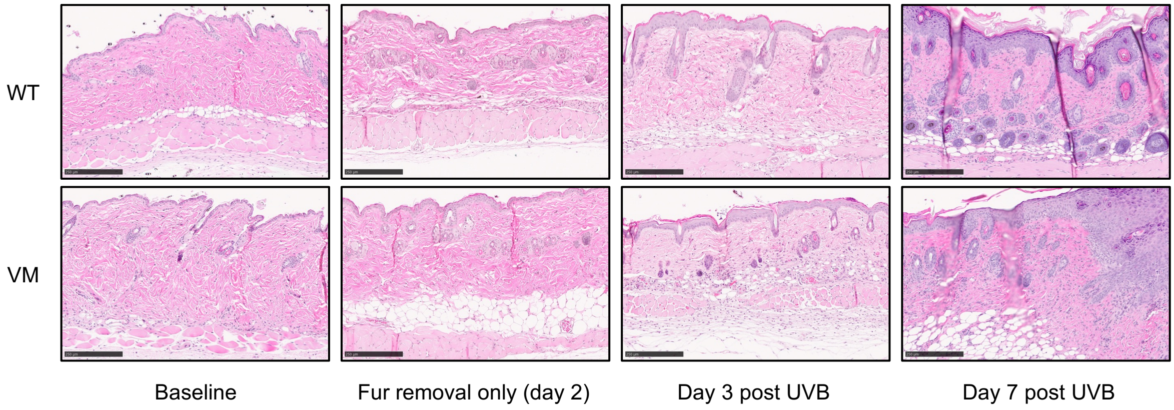


**Supplementary Figure 1. VM skin exhibits impaired wound healing compared to WT following UVB injury.** Representative H&E-stained sections of dorsal skin from WT and VM mice at baseline and following UVB exposure. At baseline and early time points (day 2 after fur removal and day 3 post-UVB), no major histological differences were observed between WT and VM mice. By day 7 post-UVB, WT skin showed near-complete resolution of inflammation, with a normal epidermal architecture, preserved adnexal structures (including hair follicles), and minimal dermal inflammatory infiltrate. In contrast, VM skin demonstrated signs of impaired resolution, including epidermal hyperplasia (acanthosis), loss of adnexal structures, and prominent dermal immune cell infiltration.
