## Supplemental Figure 2 for "Endothelial Cell Expression of STING^V154M^ Gain-of-Function Mutation Delays the Resolution of UVB-induced Skin Injury"

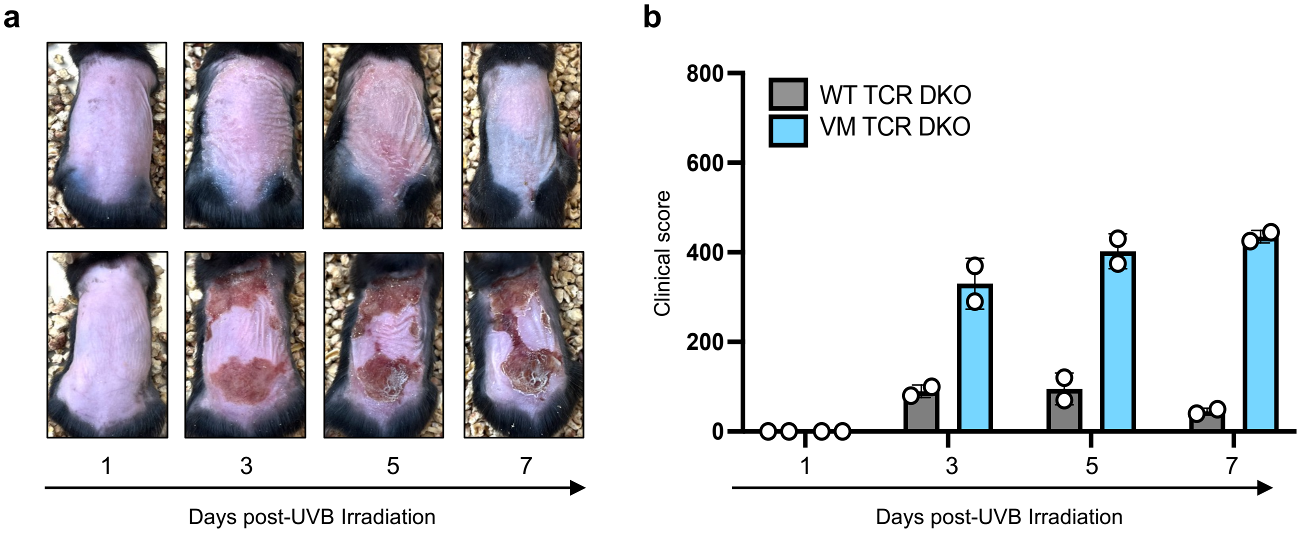


**Supplementary Figure 2. T cells are not required for exacerbated UVB-induced injury in VM mice. (**a) Representative images of UVB-irradiated TCRb-/- TCRd-/- (TCR double-knockout, DKO) mice at indicated time points. WT TCR DKO mice (top row, n=2) and VM TCR DKO (bottom row, n=2) were monitored for UVB-induced skin injury over 7 days; (b) Clinical scores for skin damage from days 1 to day 7 post-UVB.
