## Supplemental Figure 3 for "Endothelial Cell Expression of STING^V154M^ Gain-of-Function Mutation Delays the Resolution of UVB-induced Skin Injury"

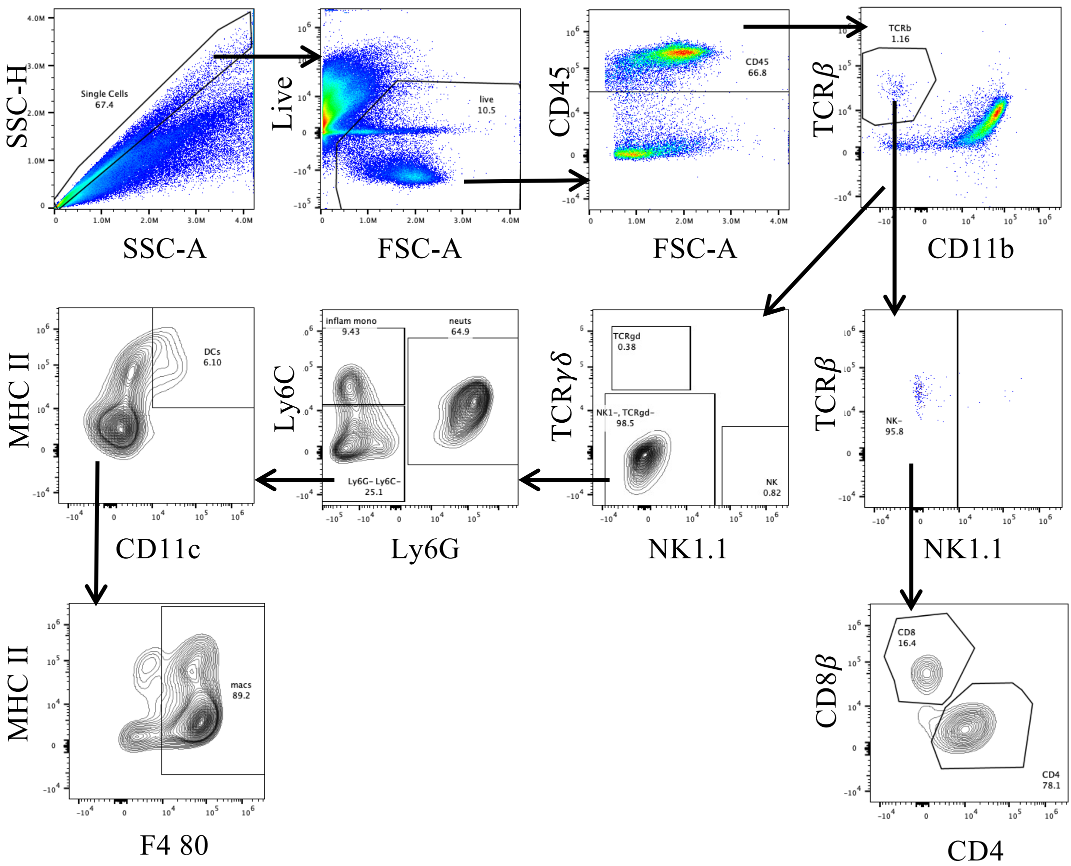


**Supplementary Figure 3. Flow cytometry gating strategy for identification of immune cell subsets infiltrating mouse skin.** Single, live cells were gated based on forward and side scatter (FSC-A/SSC-A) and a live/dead viability dye, followed by selection of CD45$^+$ hematopoietic cells. TCRβ$^+$ T cells, CD11b$^+$ myeloid cells, NK1.1$^+$ NK cells, and TCRγδ$^+$ cells were gated in subsequent steps. Among TCRβ$^+$ cells, CD4$^+$ and CD8$^+$ T cell subsets were identified. Within CD11b$^+$ cells, neutrophils (Ly6G$^+$), inflammatory monocytes (Ly6C^hi^Ly6G$^-$), and Ly6C$^-$ myeloid cells were further distinguished. Dendritic cells (CD11c$^+$MHC II$^+$) and macrophages (F4/80$^+$MHC II$^+$) were resolved as separate populations within the CD11b$^+$ compartment.
