## Supplemental Figure 4 for "Endothelial Cell Expression of STING^V154M^ Gain-of-Function Mutation Delays the Resolution of UVB-induced Skin Injury"

Cluster3 ... Genes 1 to 100

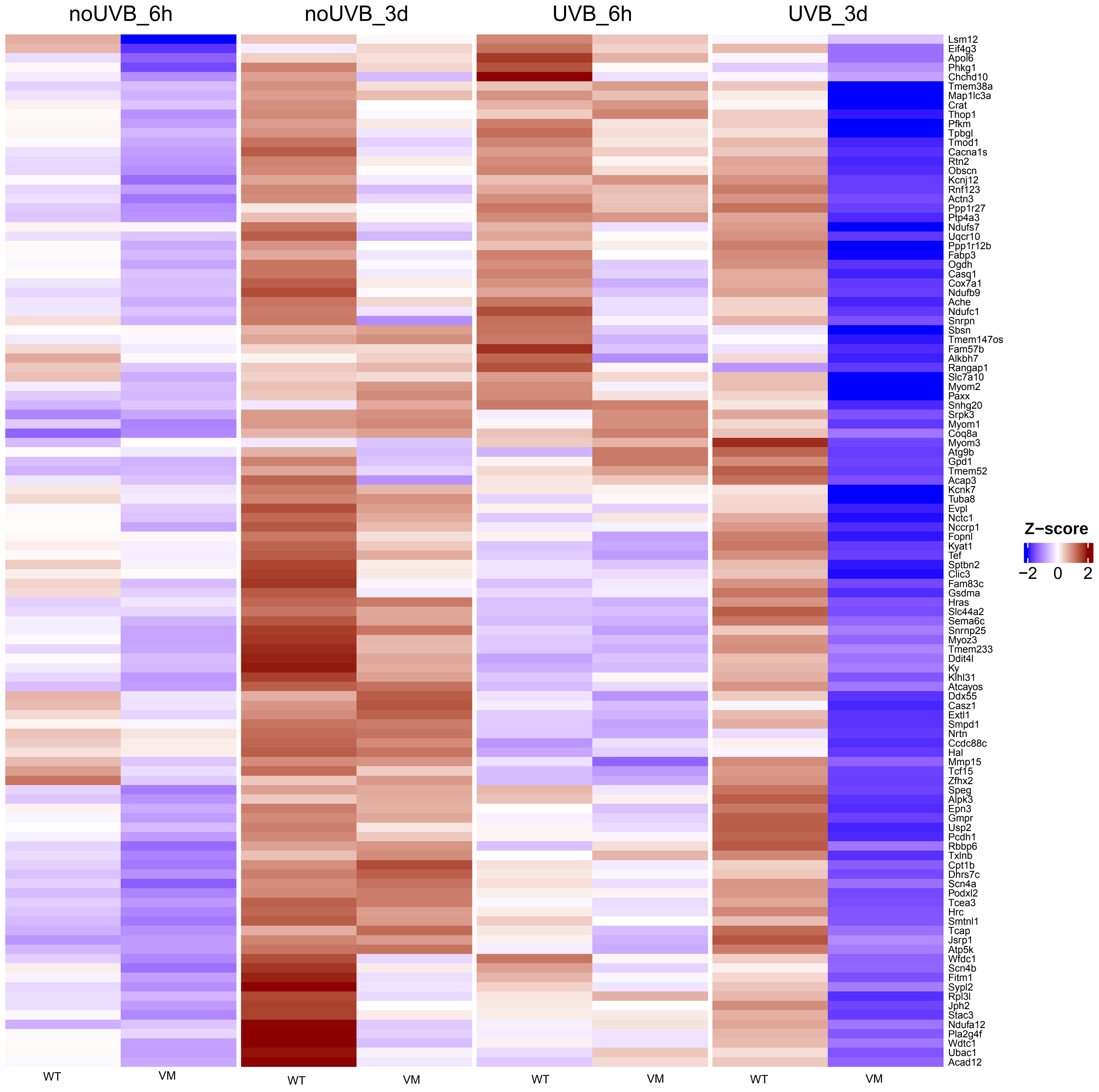

**Cluster3 ... Genes 111 to 210**

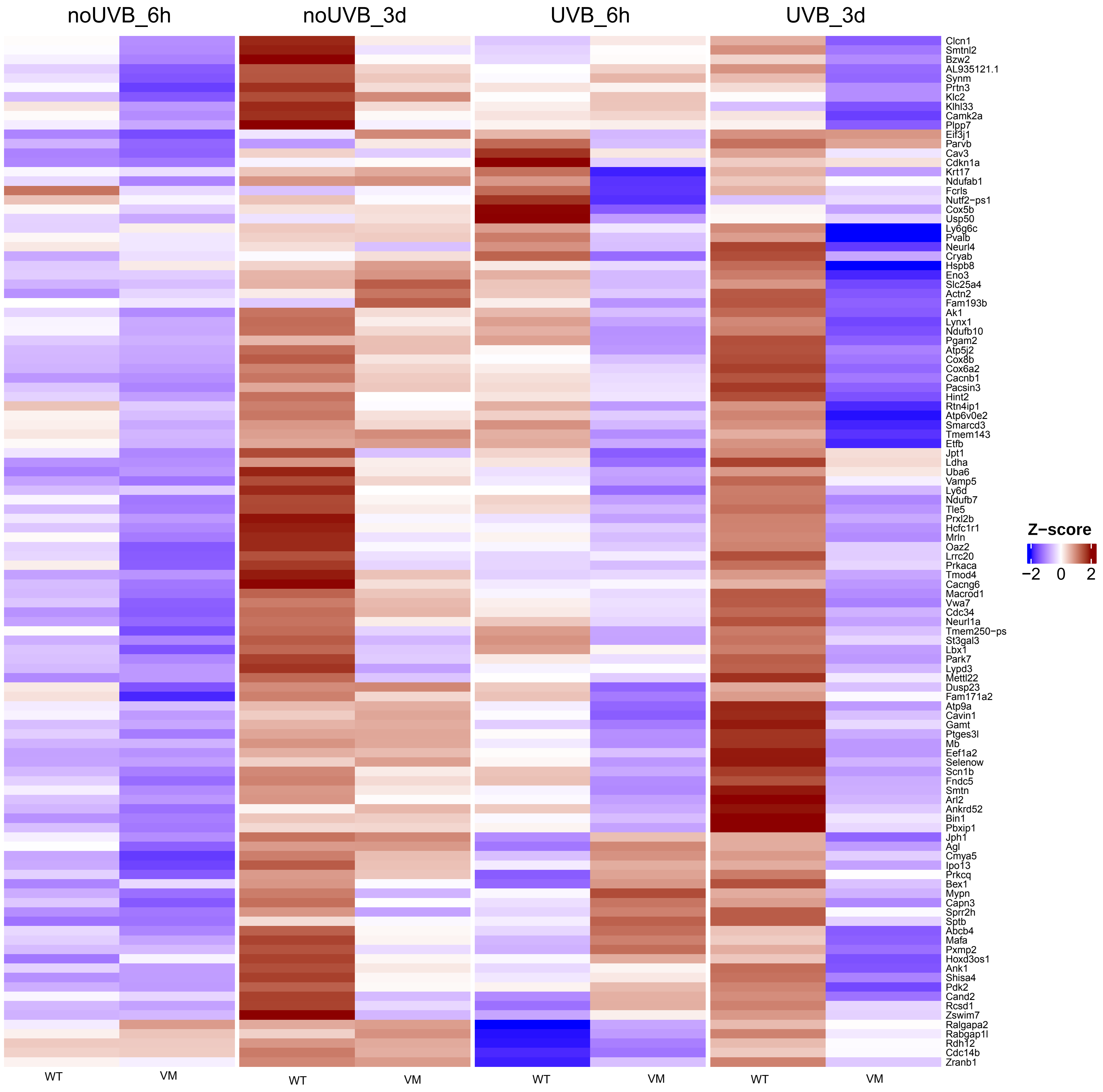

**Cluster3 ... Genes 221 to 320**

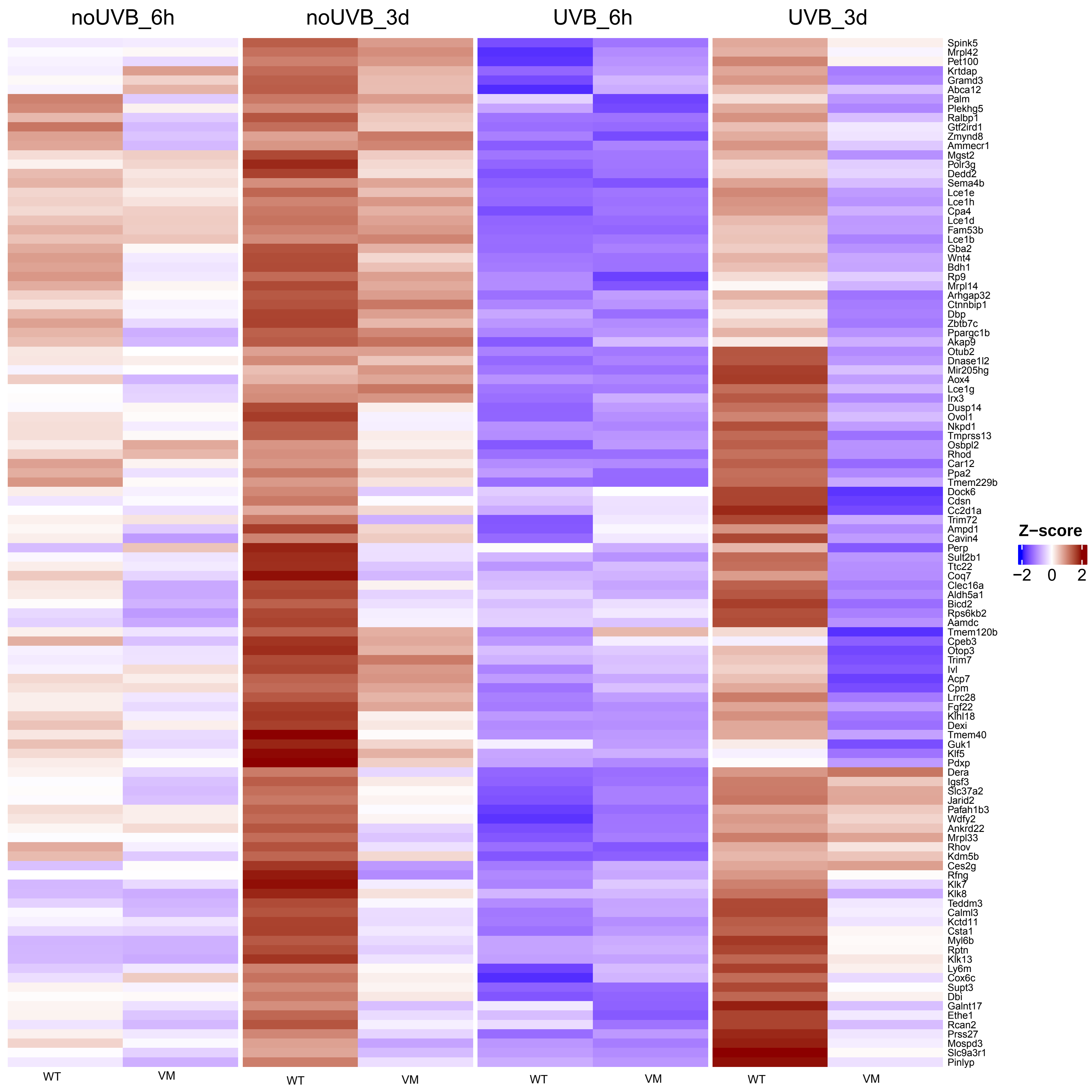

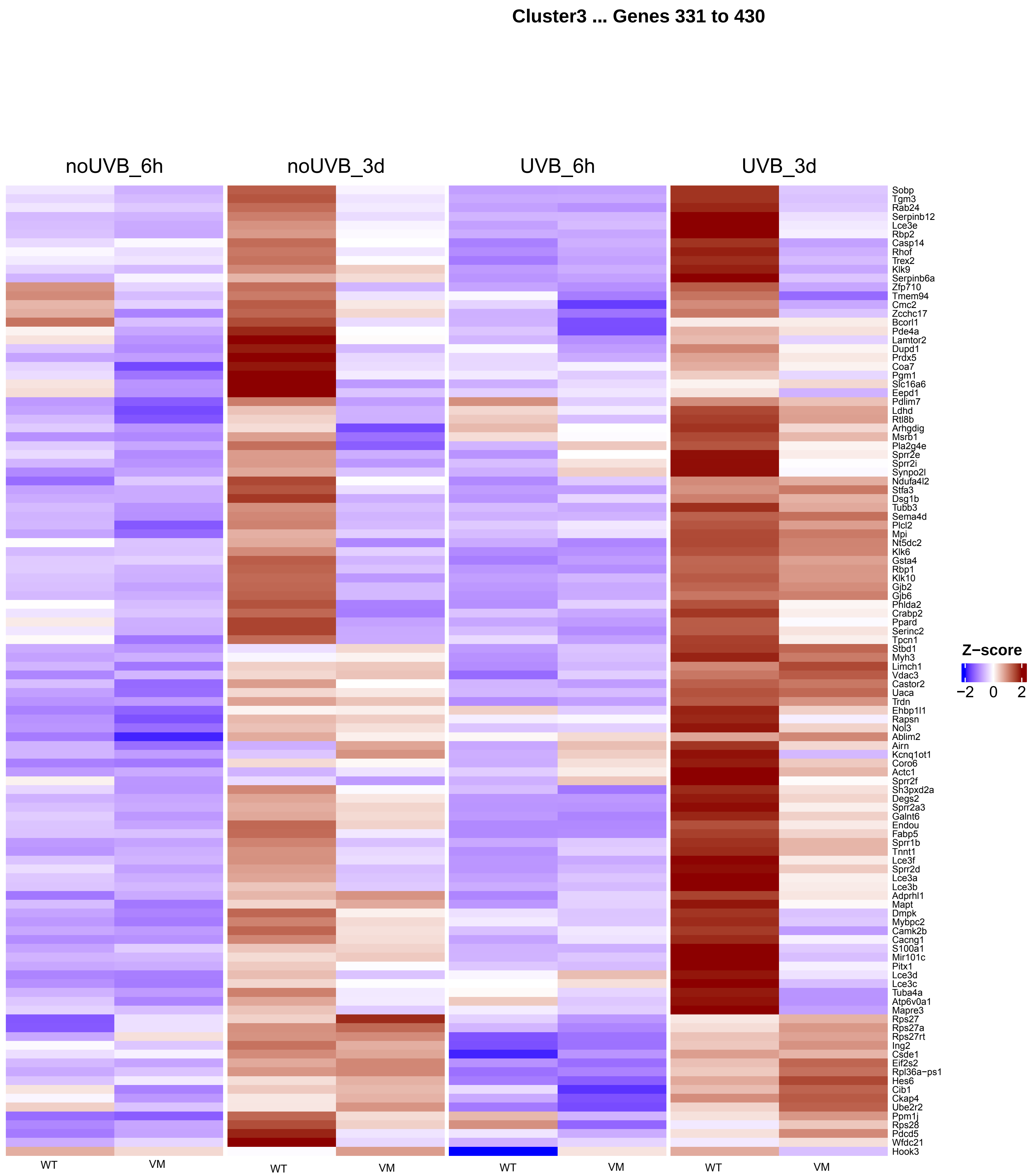
Cluster3 ... Genes 331 to 430
noUVB_6h
noUVB_3d
UVB_6h
UVB_3d
Sobp
Tgm3
Rab24
Serpinb12
Lce3e
Rbp2
Casp14
Rhof
Trex2
Klk9
Serpinb6a
Zfp710
Tmem94
Cmc2
Zcchc17
Bcorl1
Pde4a
Lamtor2
Dupd1
Prdx5
Coa7
Pgm1
Slc16a6
Eepd1
Pdlim7
Ldhd
Rtl8b
Arhgdig
Msrb1
Pla2g4e
Sprr2e
Sprr2i
Synpo2l
Ndufa4l2
Stfa3
Dsg1b
Tubb3
Sema4d
Plcl2
Mpi
Nt5dc2
Klk6
Gsta4
Rbp1
Klk10
Gjb2
Gjb6
Phlda2
Crabp2
Ppard
Serinc2
Tpcn1
Stbd1
Myh3
Limch1
Vdac3
Castor2
Uaca
Trdn
Ehbp1l1
Rapsn
Nol3
Ablim2
Airn
Kcnq1ot1
Coro6
Actc1
Sprr2f
Sh3pxd2a
Degs2
Sprr2a3
Galnt6
Endou
Fabp5
Sprr1b
Tnnt1
Lce3f
Sprr2d
Lce3a
Lce3b
Adprhl1
Mapt
Dmpk
Mybpc2
Camk2b
Cacng1
S100a1
Mir101c
Pitx1
Lce3d
Lce3c
Tuba4a
Atp6v0a1
Mapre3
Rps27
Rps27a
Rps27rt
Ing2
Csde1
Eif2s2
Rpl36a−ps1
Hes6
Cib1
Ckap4
Ube2r2
Ppm1j
Rps28
Pdcd5
Wfdc21
Hook3
Z−score
−2
0
2
WT
VM
WT
VM
WT
VM
WT
VM

#### Cluster3 ... Genes 441 to 540

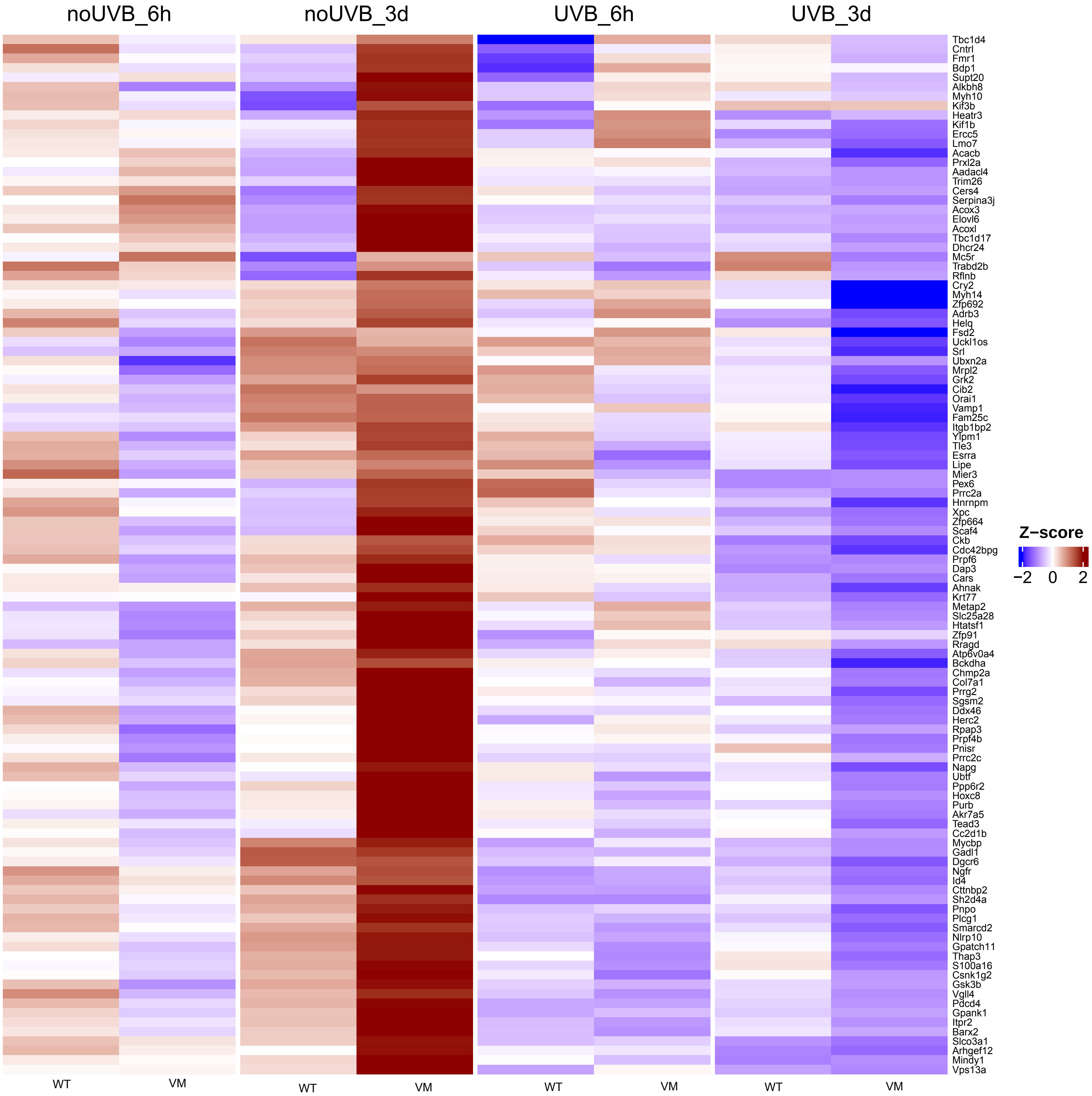

#### Cluster3 ... Genes 551 to 650

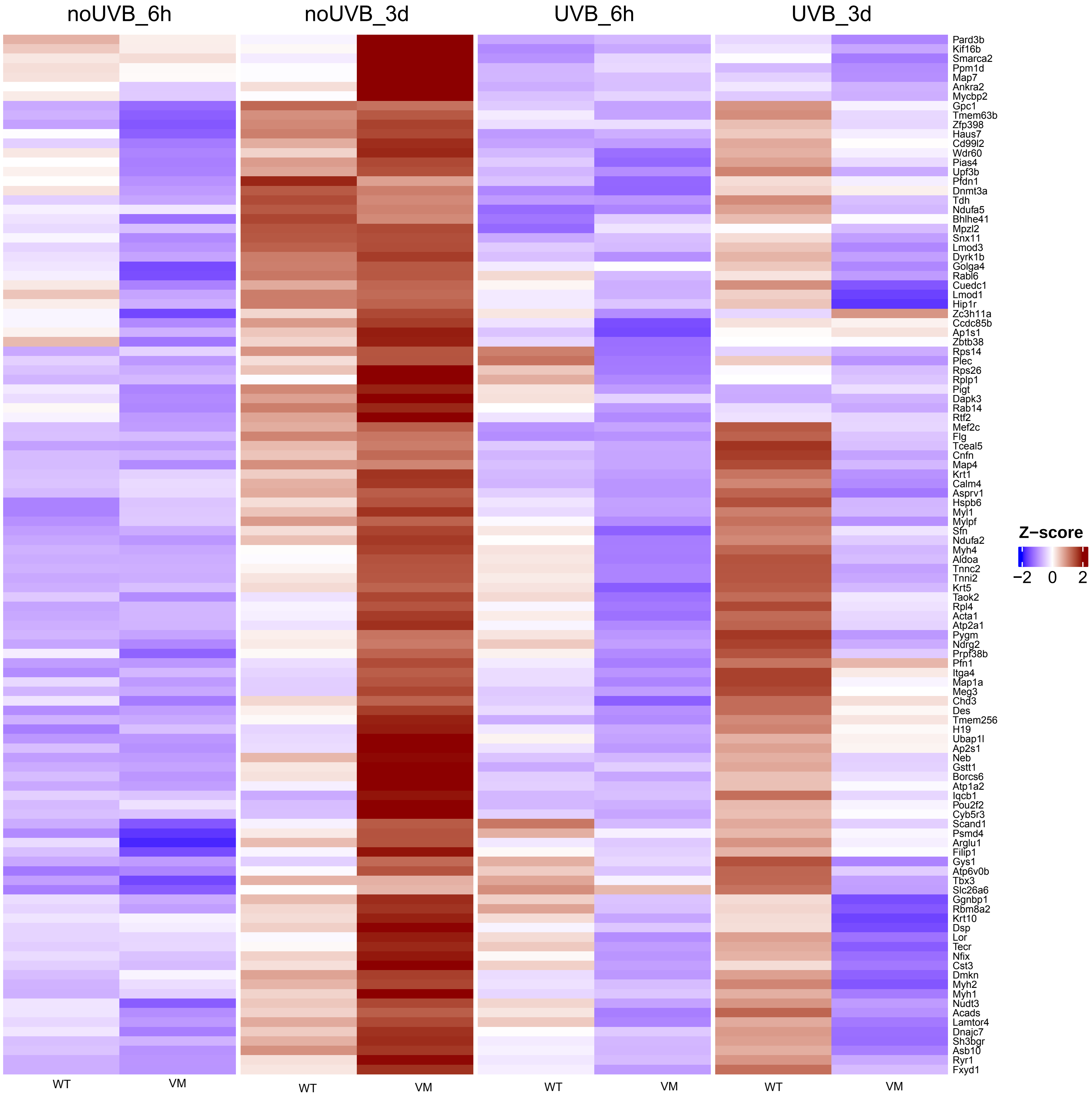

**Cluster3 ... Genes 661 to 760**

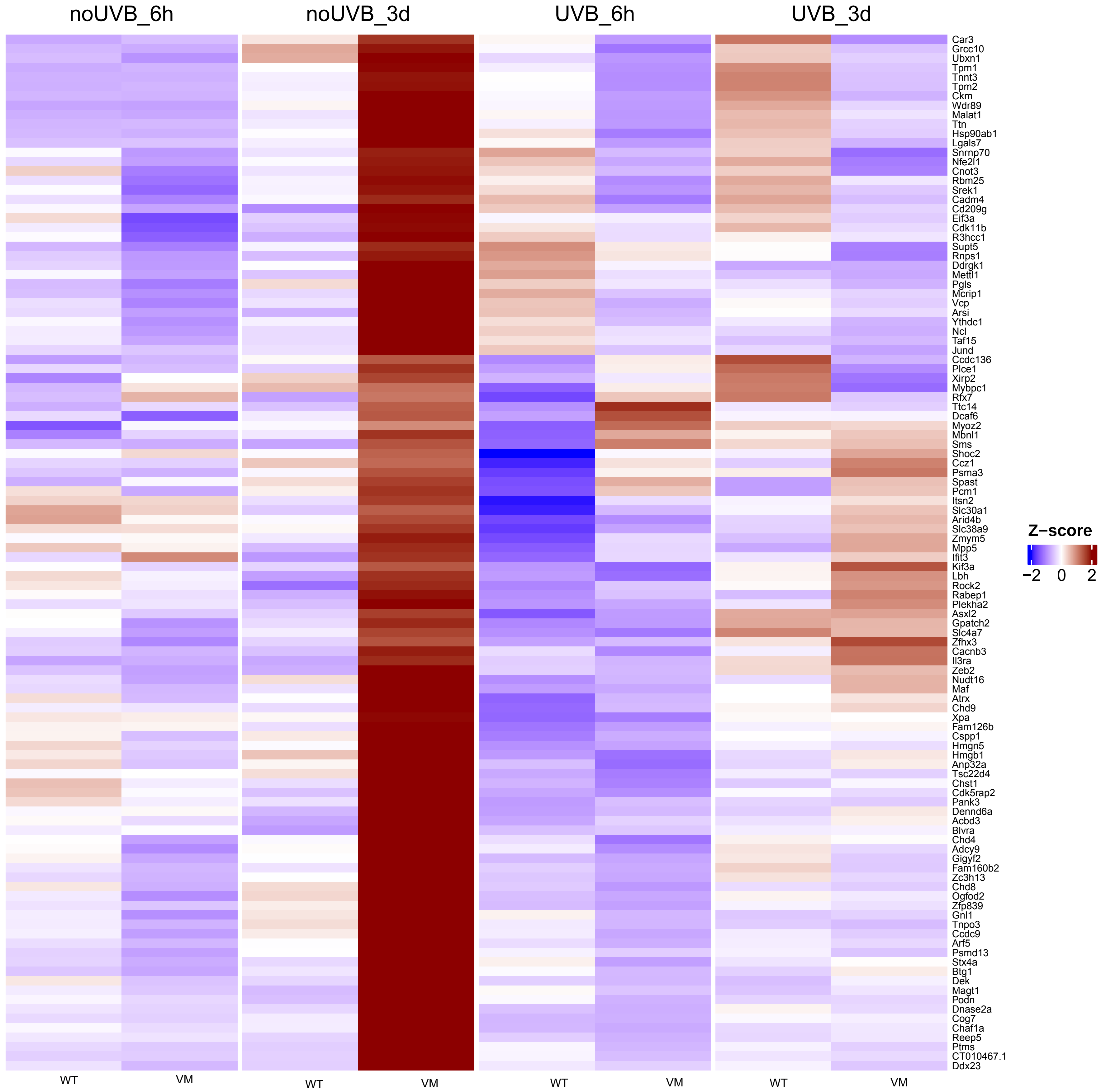

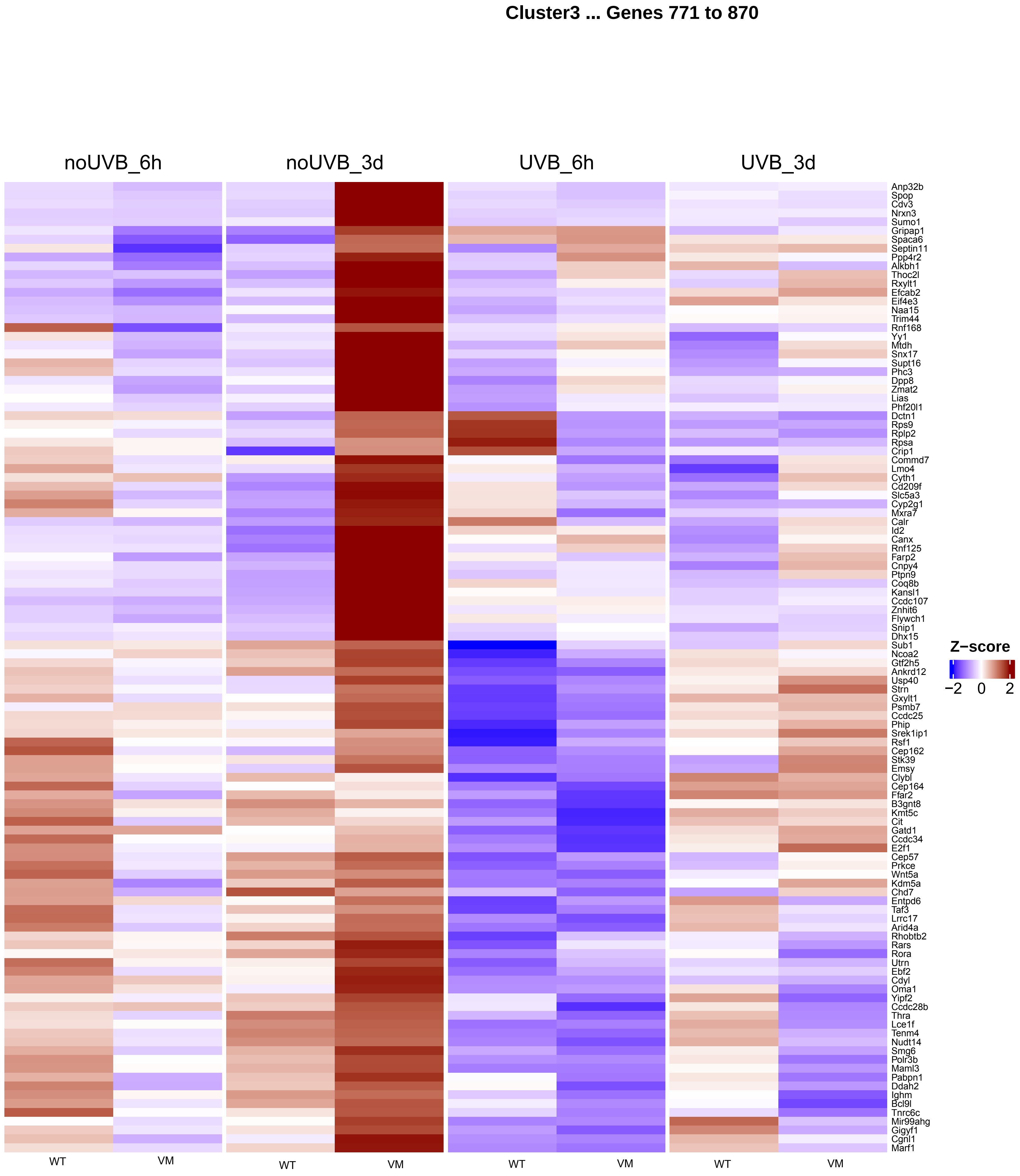
Cluster3 ... Genes 771 to 870
noUVB_6h
noUVB_3d
UVB_6h
UVB_3d
Anp32b
Spop
Cdv3
Nrxn3
Sumo1
Gripap1
Spaca6
Septin11
Ppp4r2
Alkbh1
Thoc2l
Rxylt1
Efcab2
Eif4e3
Naa15
Trim44
Rnf168
Yy1
Mtdh
Snx17
Supt16
Phc3
Dpp8
Zmat2
Lias
Phf20l1
Dctn1
Rps9
Rplp2
Rpsa
Crip1
Commd7
Lmo4
Cyth1
Cd209f
Slc5a3
Cyp2g1
Mxra7
Calr
Id2
Canx
Rnf125
Farp2
Cnpy4
Ptpn9
Coq8b
Kansl1
Ccdc107
Znhit6
Flywch1
Snip1
Dhx15
Sub1
Ncoa2
Gtf2h5
Ankrd12
Usp40
Strn
Gxylt1
Psmb7
Ccdc25
Phip
Srek1ip1
Rsf1
Cep162
Stk39
Emsy
Clybl
Cep164
Ffar2
B3gnt8
Kmt5c
Cit
Gatd1
Ccdc34
E2f1
Cep57
Prkce
Wnt5a
Kdm5a
Chd7
Entpd6
Taf3
Lrrc17
Arid4a
Rhobtb2
Rars
Rora
Utrn
Ebf2
Cdyl
Oma1
Yipf2
Ccdc28b
Thra
Lce1f
Tenm4
Nudt14
Smg6
Polr3b
Maml3
Pabpn1
Ddah2
Ighm
Bcl9l
Tnrc6c
Mir99ahg
Gigyf1
Cgnl1
Marf1
Z−score
−2
0
2
WT
VM
WT
VM
WT
VM
WT
VM

**Cluster3 ... Genes 881 to 961**

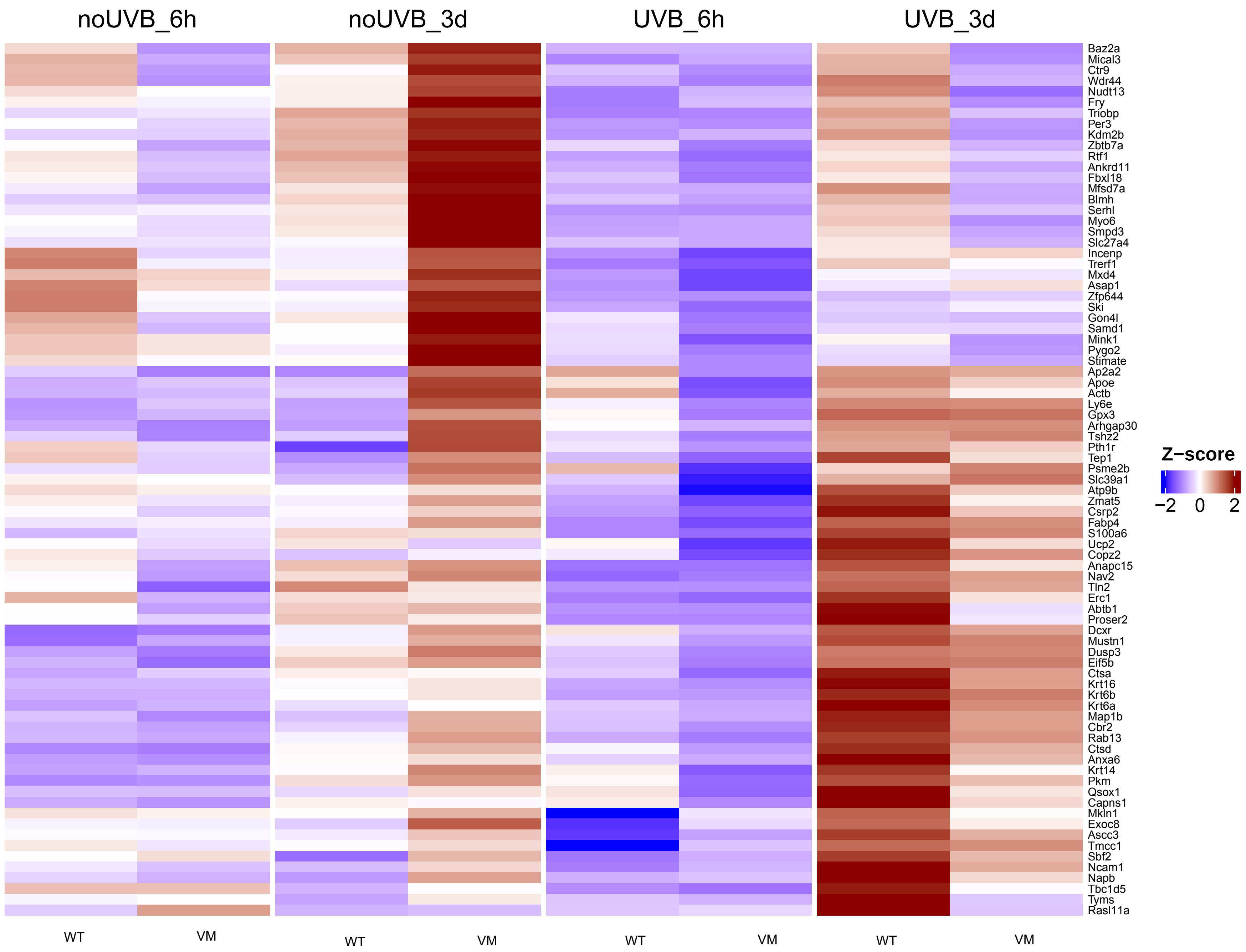

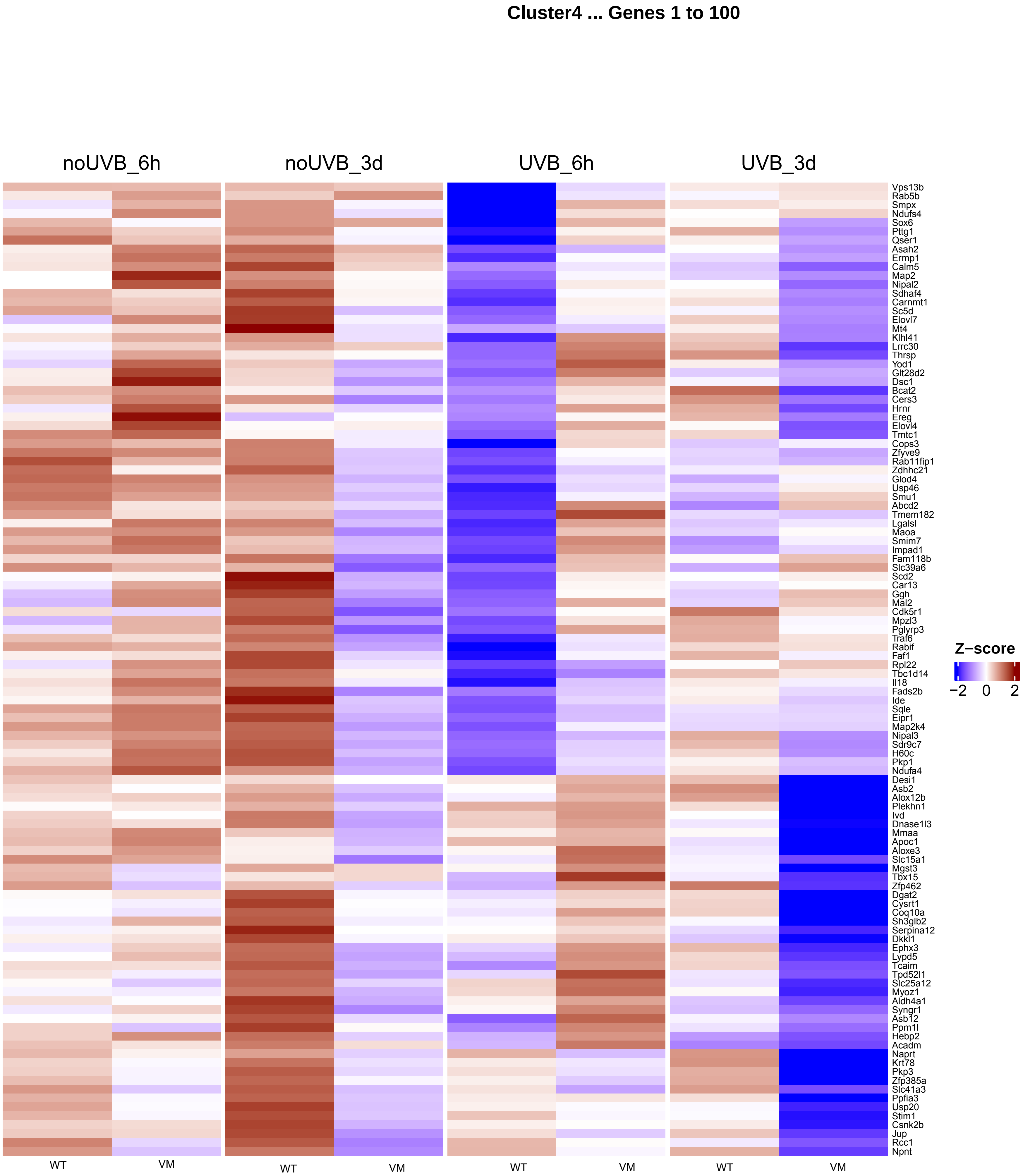
Cluster4 ... Genes 1 to 100
noUVB_6h
noUVB_3d
UVB_6h
UVB_3d
Vps13b
Rab5b
Smpx
Ndufs4
Sox6
Pttg1
Qser1
Asah2
Ermp1
Calm5
Map2
Nipal2
Sdhaf4
Carnmt1
Sc5d
Elovl7
Mt4
Klhl41
Lrrc30
Thrsp
Yod1
Glt28d2
Dsc1
Bcat2
Cers3
Hrnr
Ereg
Elovl4
Tmtc1
Cops3
Zfyve9
Rab11fip1
Zdhhc21
Glod4
Usp46
Smu1
Abcd2
Tmem182
Lgalsl
Maoa
Smim7
Impad1
Fam118b
Slc39a6
Scd2
Car13
Ggh
Mal2
Cdk5r1
Mpzl3
Pglyrp3
Traf6
Rabif
Faf1
Rpl22
Tbc1d14
Il18
Fads2b
Ide
Sqle
Eipr1
Map2k4
Nipal3
Sdr9c7
H60c
Pkp1
Ndufa4
Desi1
Asb2
Alox12b
Plekhn1
Ivd
Dnase1l3
Mmaa
Apoc1
Aloxe3
Slc15a1
Mgst3
Tbx15
Zfp462
Dgat2
Cysrt1
Coq10a
Sh3glb2
Serpina12
Dkkl1
Ephx3
Lypd5
Tcaim
Tpd52l1
Slc25a12
Myoz1
Aldh4a1
Syngr1
Asb12
Ppm1l
Hebp2
Acadm
Naprt
Krt78
Pkp3
Zfp385a
Slc41a3
Ppfia3
Usp20
Stim1
Csnk2b
Jup
Rcc1
Npnt
WT
VM
WT
VM
WT
VM
WT
VM
Z−score
−2
0
2

Cluster4 ... Genes 111 to 210

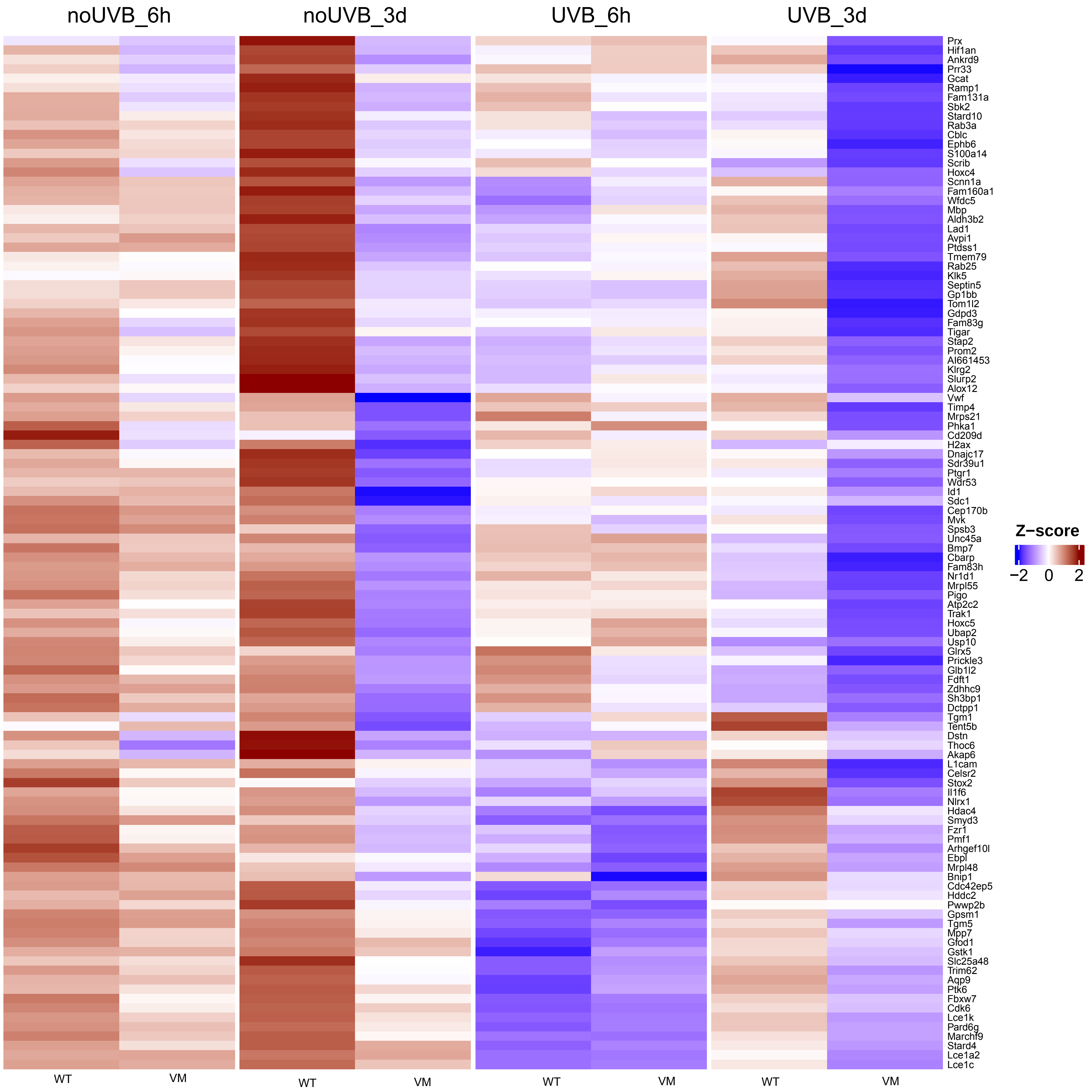

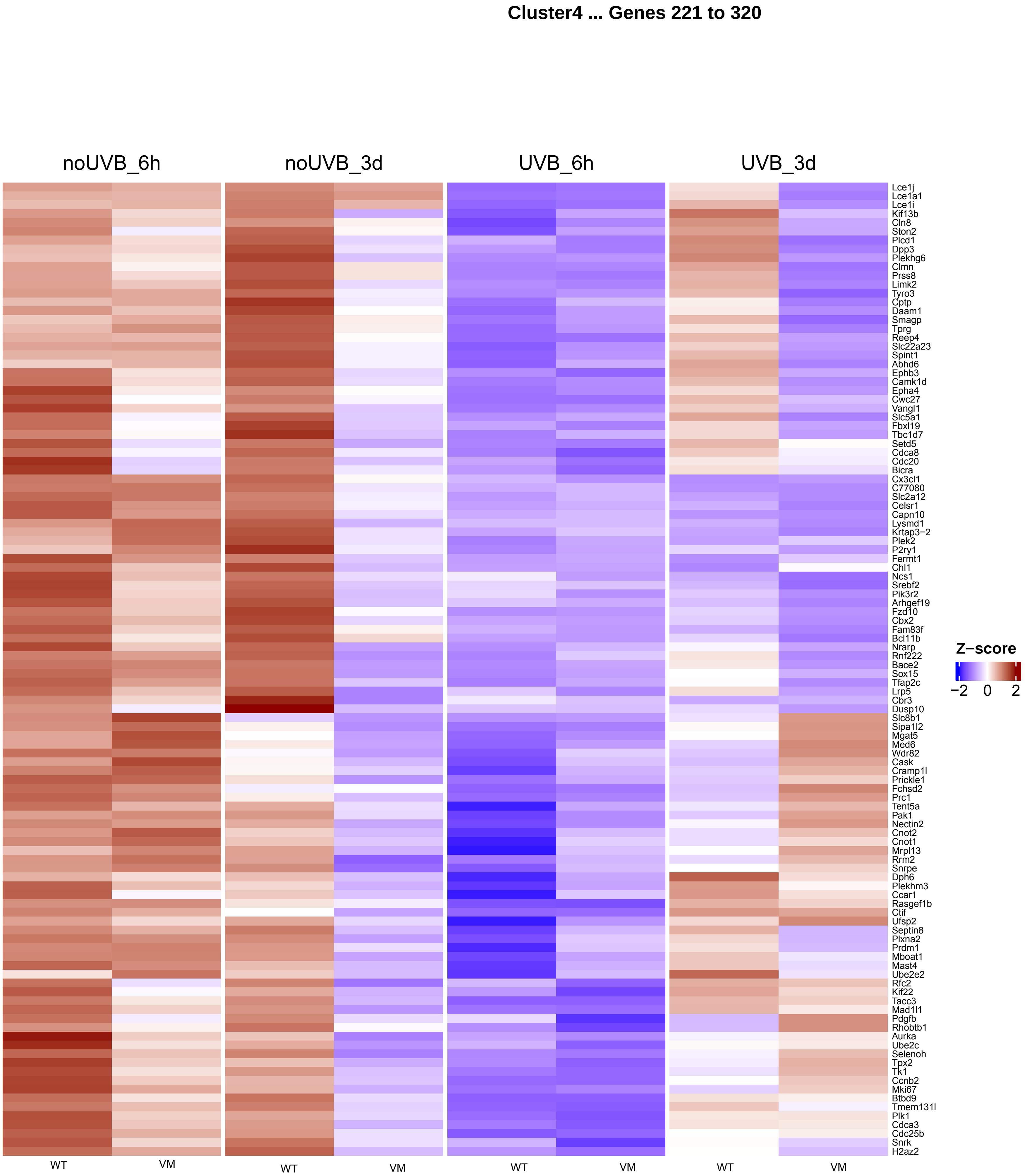
Cluster4 ... Genes 221 to 320
noUVB_6h
noUVB_3d
UVB_6h
UVB_3d
Lce1j
Lce1a1
Lce1i
Kif13b
Cln8
Ston2
Plcd1
Dpp3
Plekhg6
Clmn
Prss8
Limk2
Tyro3
Cptp
Daam1
Smagp
Tprg
Reep4
Slc22a23
Spint1
Abhd6
Ephb3
Camk1d
Epha4
Cwc27
Vangl1
Slc5a1
Fbxl19
Tbc1d7
Setd5
Cdca8
Cdc20
Bicra
Cx3cl1
C77080
Slc2a12
Celsr1
Capn10
Lysmd1
Krtap3−2
Plek2
P2ry1
Fermt1
Chl1
Ncs1
Srebf2
Pik3r2
Arhgef19
Fzd10
Cbx2
Fam83f
Bcl11b
Nrarp
Rnf222
Bace2
Sox15
Tfap2c
Lrp5
Cbr3
Dusp10
Slc8b1
Sipa1l2
Mgat5
Med6
Wdr82
Cask
Cramp1l
Prickle1
Fchsd2
Prc1
Tent5a
Pak1
Nectin2
Cnot2
Cnot1
Mrpl13
Rrm2
Snrpe
Dph6
Plekhm3
Ccar1
Rasgef1b
Ctif
Ufsp2
Septin8
Plxna2
Prdm1
Mboat1
Mast4
Ube2e2
Rfc2
Kif22
Tacc3
Mad1l1
Pdgfb
Rhobtb1
Aurka
Ube2c
Selenoh
Tpx2
Tk1
Ccnb2
Mki67
Btbd9
Tmem131l
Plk1
Cdca3
Cdc25b
Snrk
H2az2
Z−score
−2
0
2
WT
VM
WT
VM
WT
VM
WT
VM

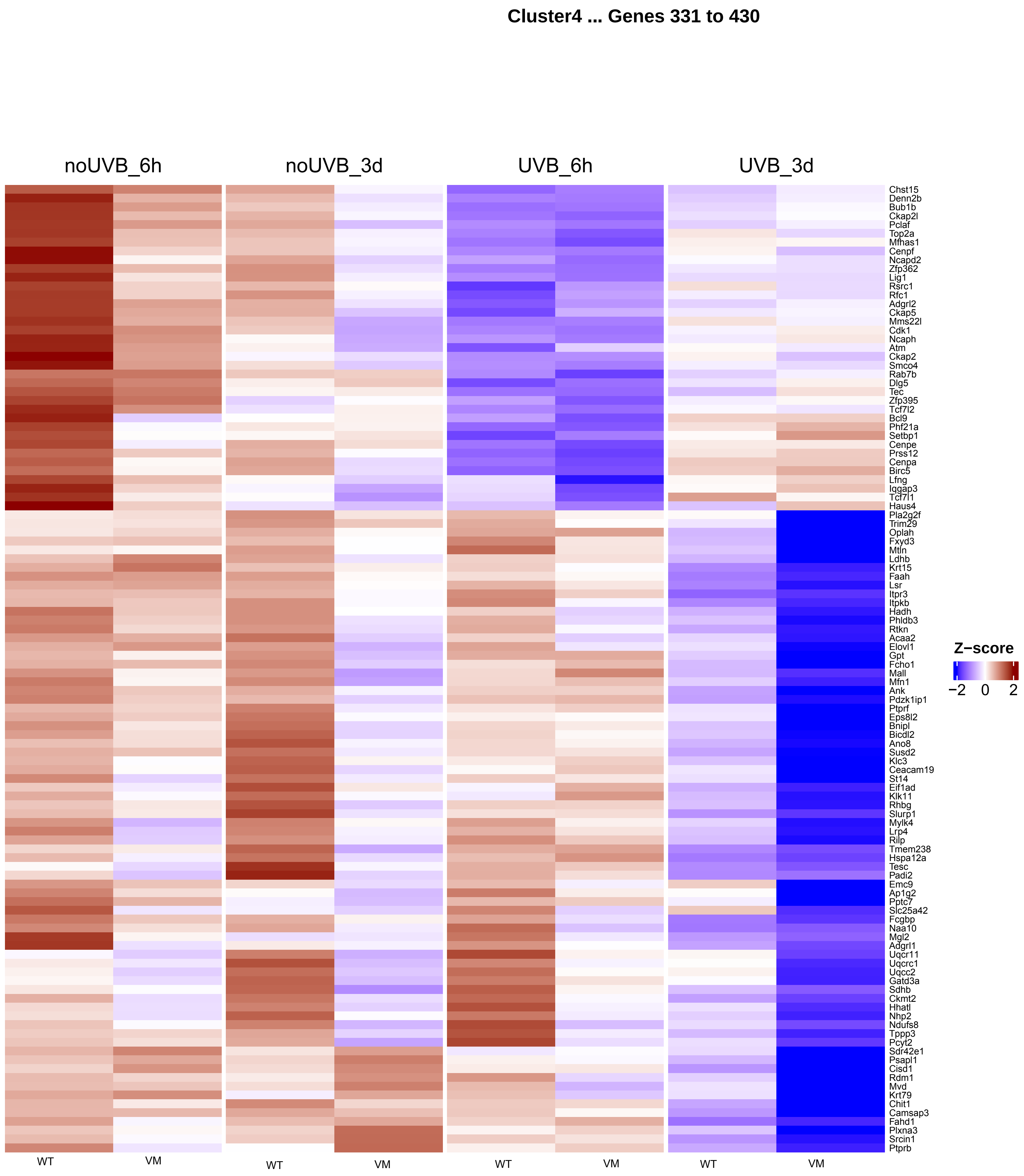
Cluster4 ... Genes 331 to 430
noUVB_6h
noUVB_3d
UVB_6h
UVB_3d
Chst15
Denn2b
Bub1b
Ckap2l
Pclaf
Top2a
Mfhas1
Cenpf
Ncapd2
Zfp362
Lig1
Rsrc1
Rfc1
Adgrl2
Ckap5
Mms22l
Cdk1
Ncaph
Atm
Ckap2
Smco4
Rab7b
Dlg5
Tec
Zfp395
Tcf7l2
Bcl9
Phf21a
Setbp1
Cenpe
Prss12
Cenpa
Birc5
Lfng
Iqgap3
Tcf7l1
Haus4
Pla2g2f
Trim29
Oplah
Fxyd3
Mtln
Ldhb
Krt15
Faah
Lsr
Itpr3
Itpkb
Hadh
Phldb3
Rtkn
Acaa2
Elovl1
Gpt
Fcho1
Mall
Mfn1
Ank
Pdzk1ip1
Ptprf
Eps8l2
Bnipl
Bicdl2
Ano8
Susd2
Klc3
Ceacam19
St14
Eif1ad
Klk11
Rhbg
Slurp1
Mylk4
Lrp4
Rilp
Tmem238
Hspa12a
Tesc
Padi2
Emc9
Ap1g2
Pptc7
Slc25a42
Fcgbp
Naa10
Mgl2
Adgrl1
Uqcr11
Uqcrc1
Uqcc2
Gatd3a
Sdhb
Ckmt2
Hhatl
Nhp2
Ndufs8
Tppp3
Pcyt2
Sdr42e1
Psapl1
Cisd1
Rdm1
Mvd
Krt79
Chit1
Camsap3
Fahd1
Plxna3
Srcin1
Ptprb
Z−score
−2
0
2
WT
VM
WT
VM
WT
VM
WT
VM

**Cluster4 ... Genes 441 to 540**

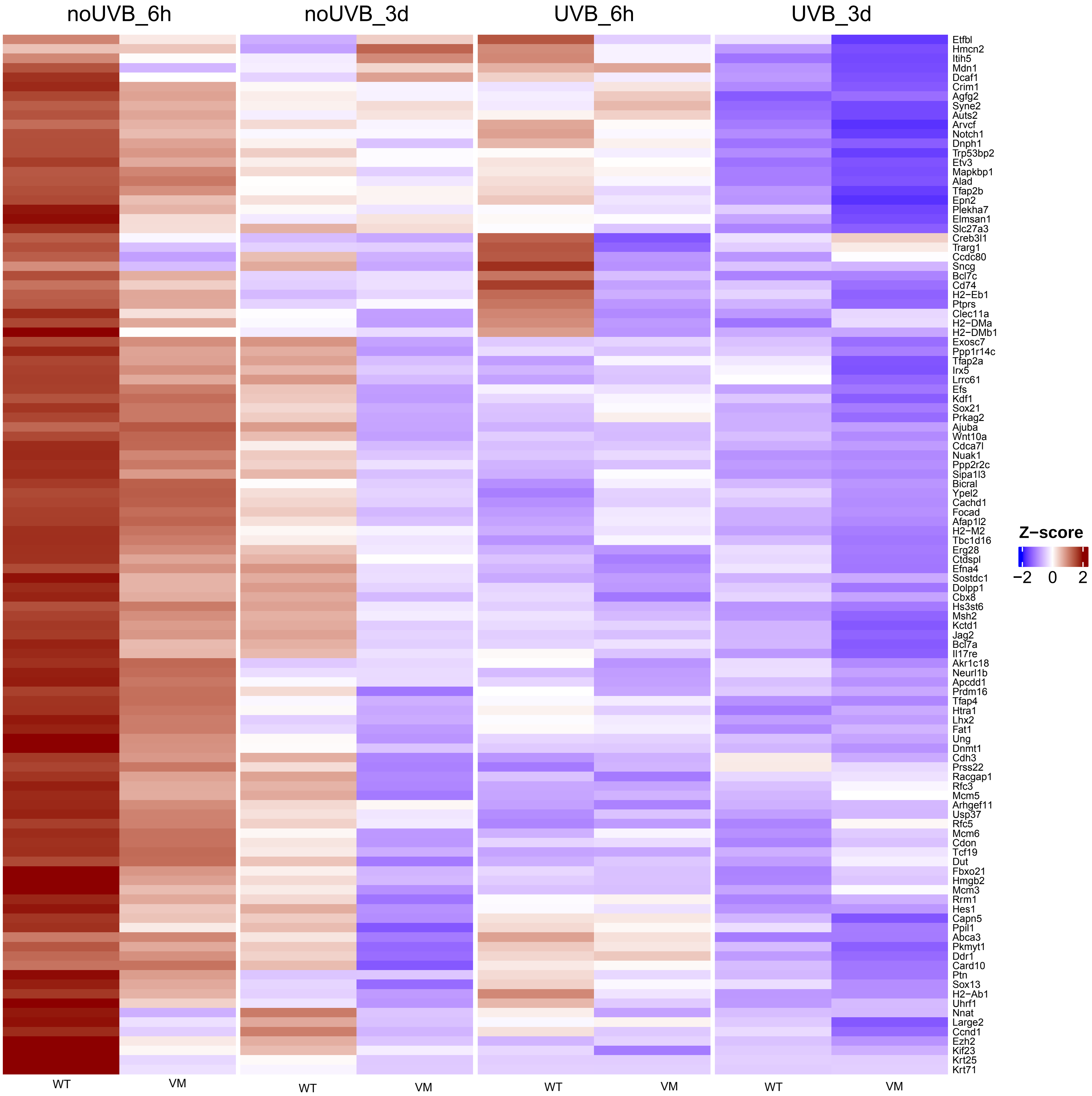

Cluster4 ... Genes 551 to 650

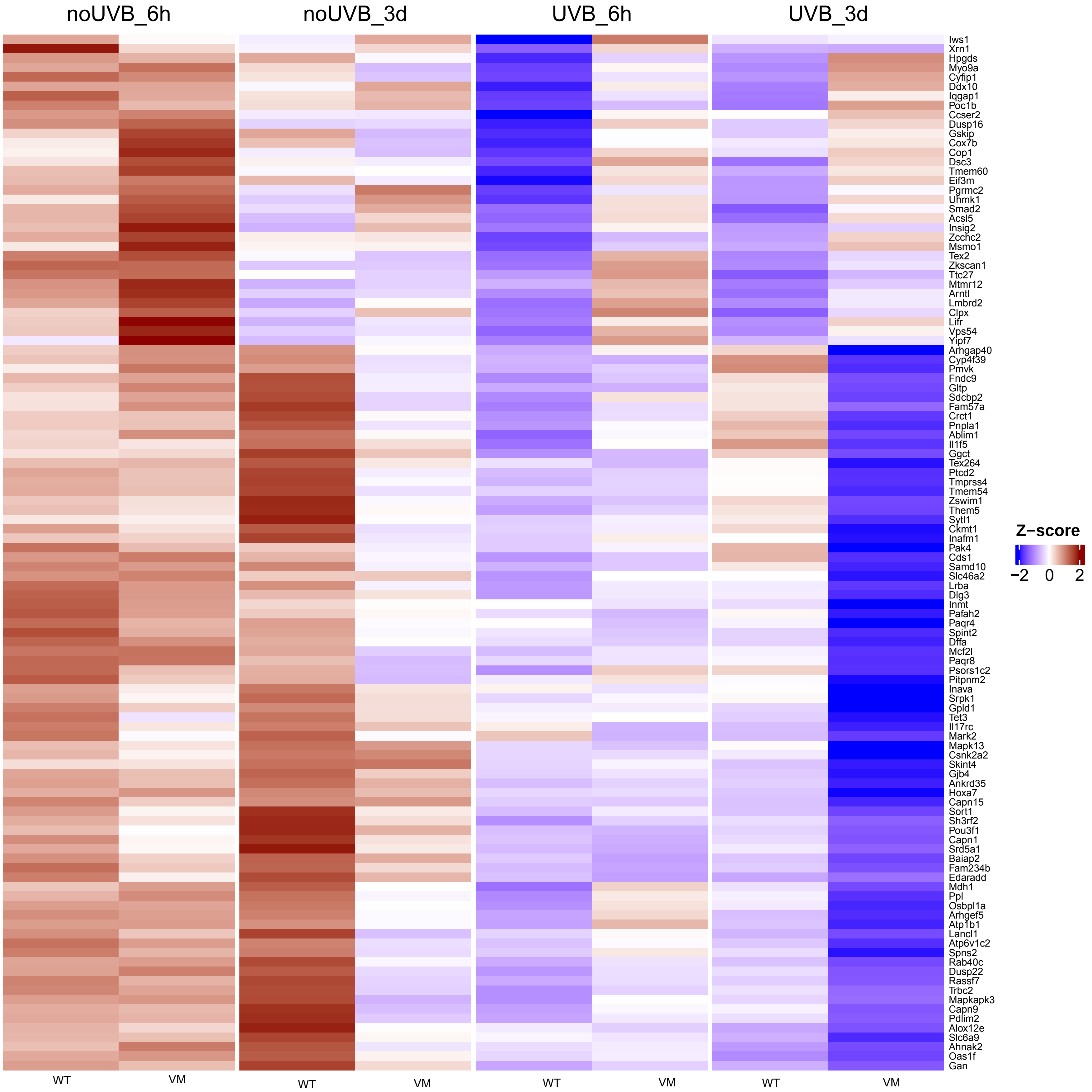

**Cluster4 ... Genes 661 to 760**

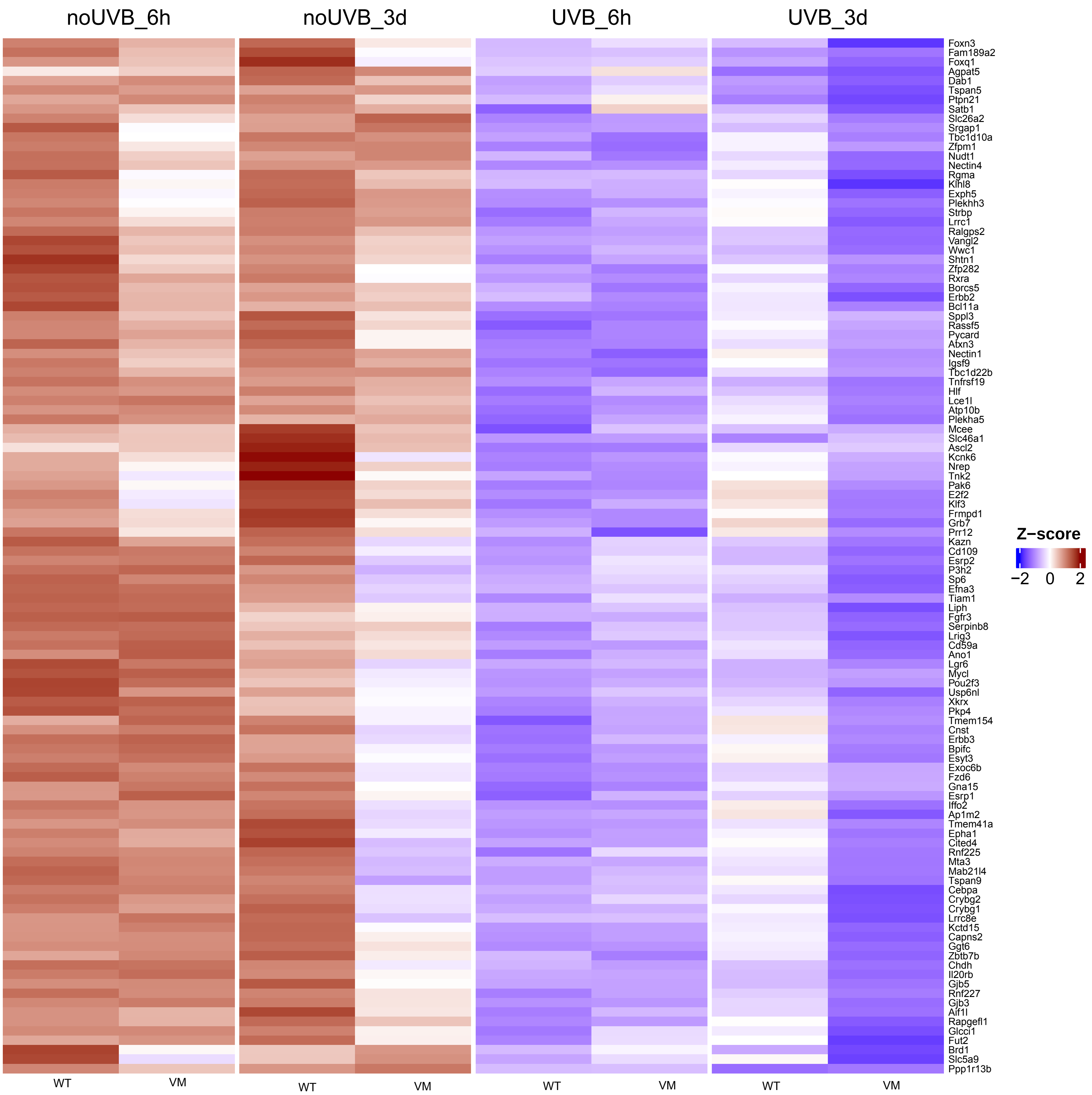

**Cluster4 ... Genes 771 to 870**

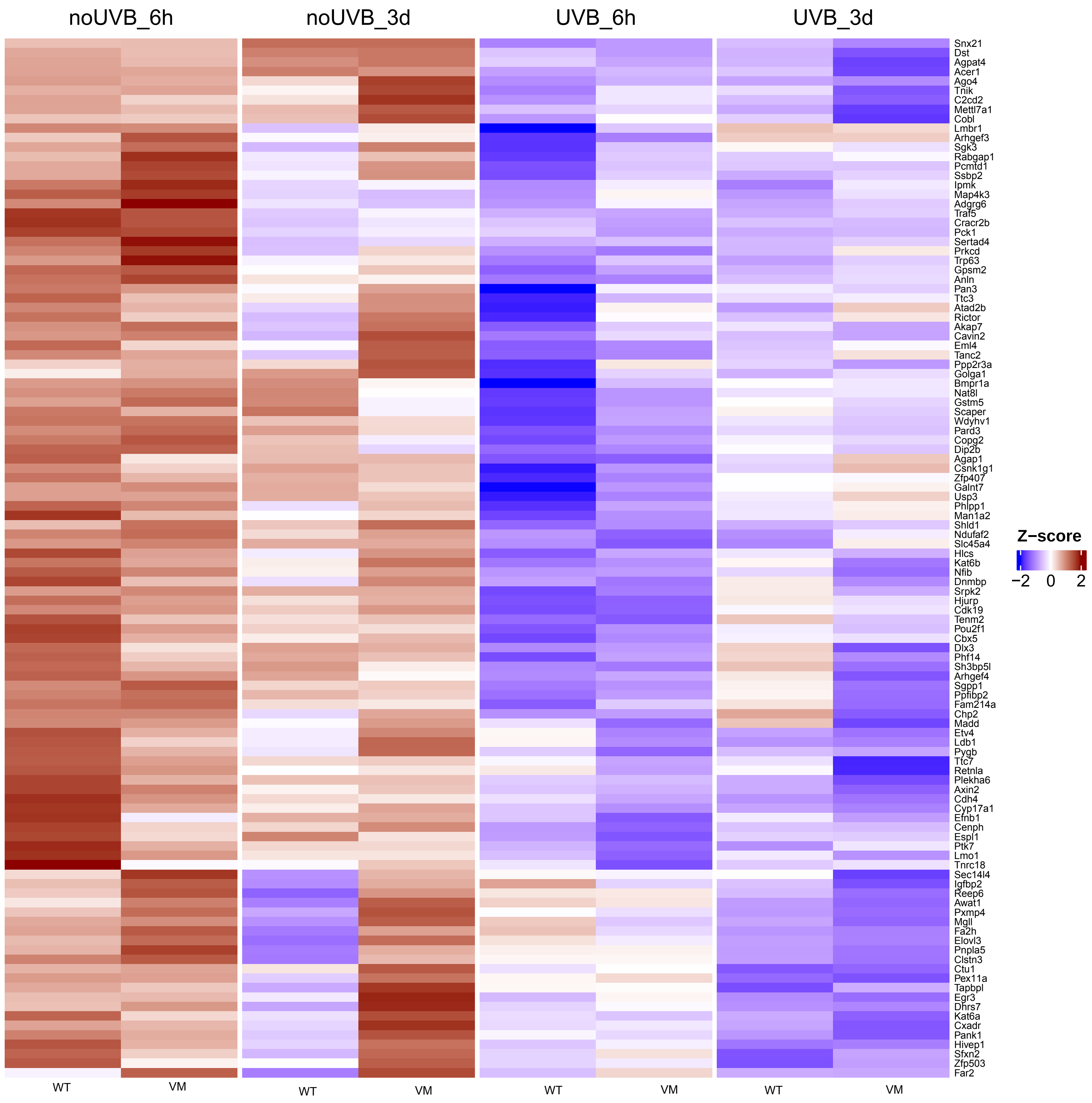

**Cluster4 ... Genes 881 to 980**

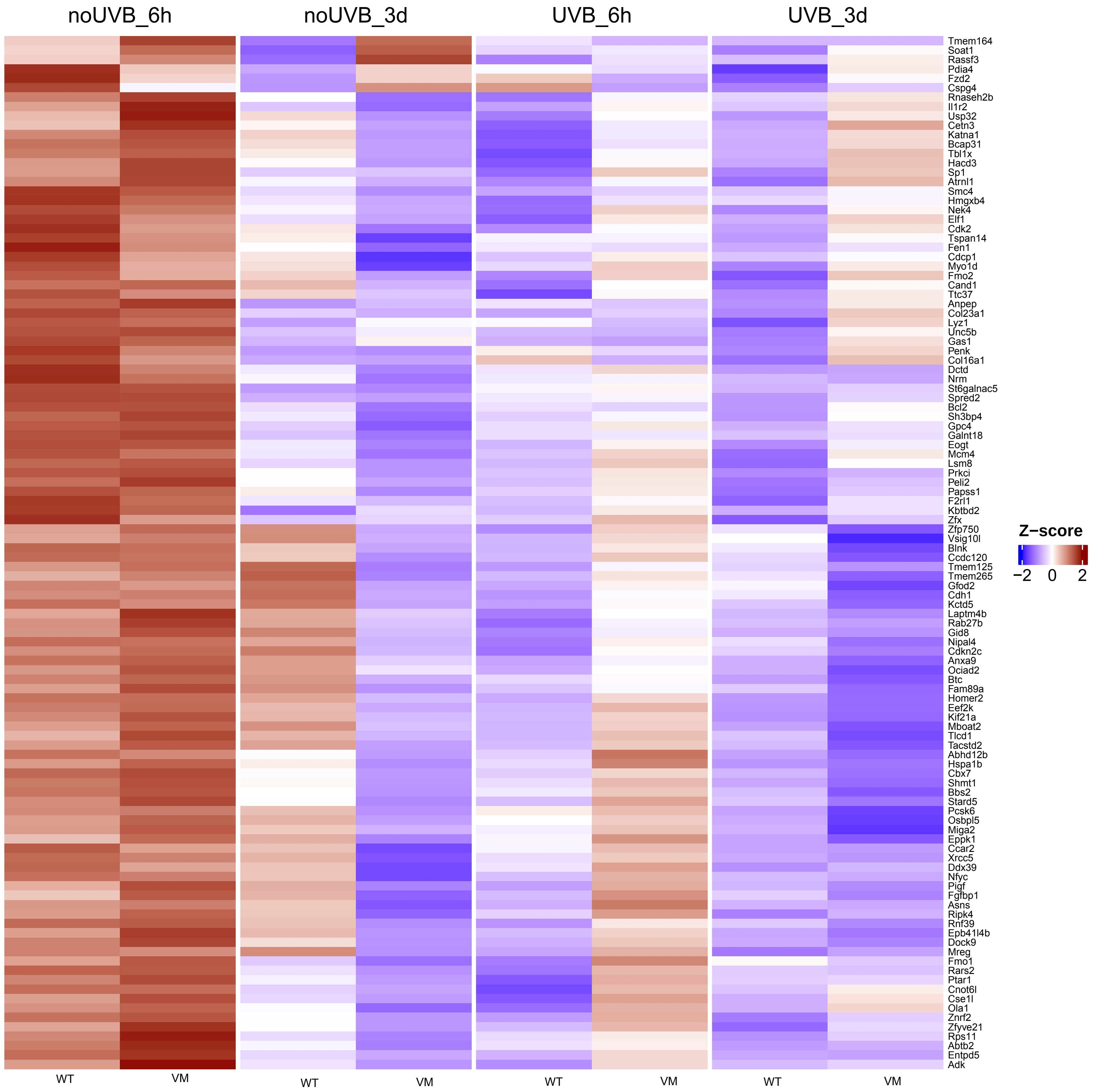

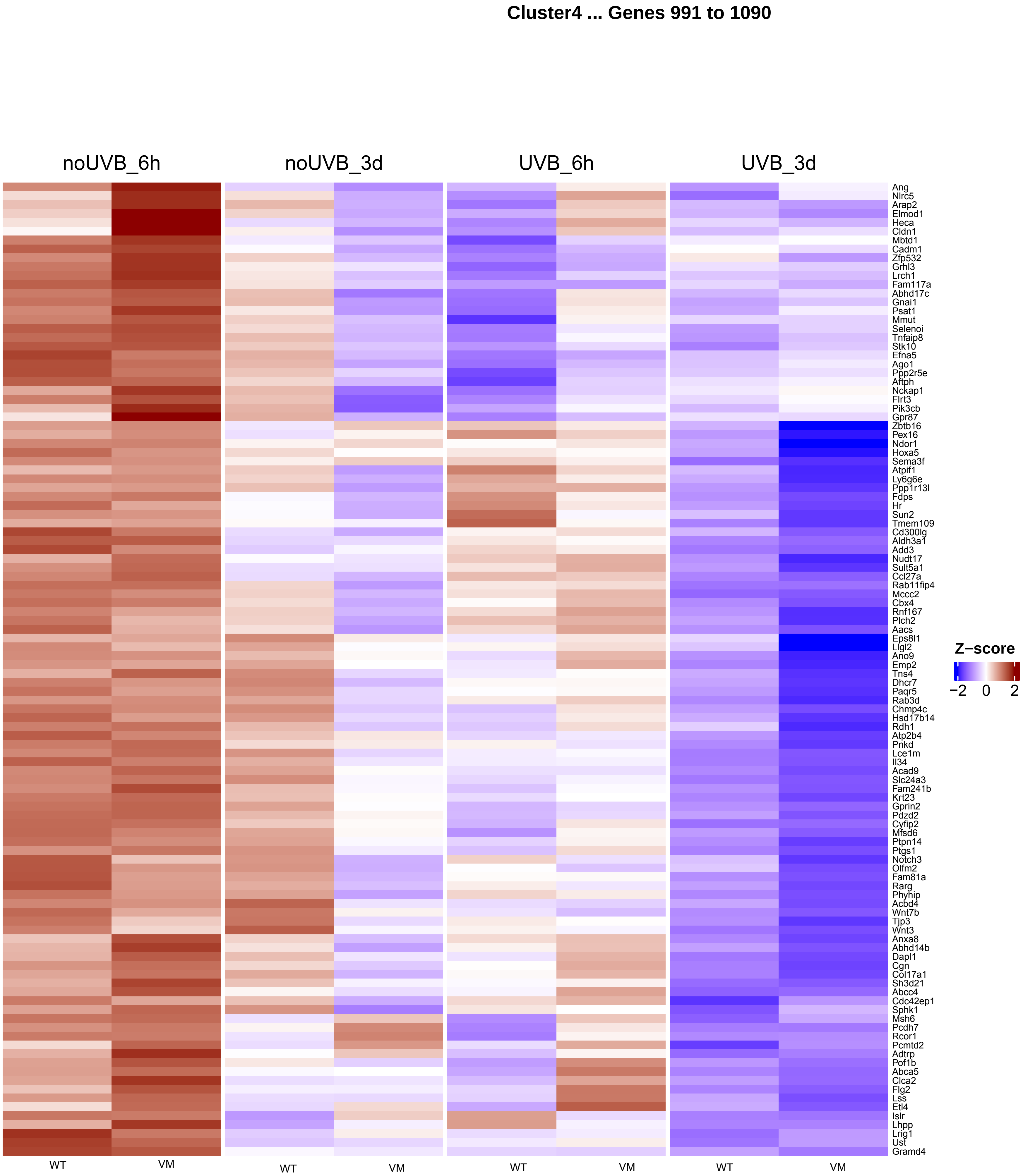
Cluster4 ... Genes 991 to 1090
noUVB_6h
noUVB_3d
UVB_6h
UVB_3d
Ang
Nlrc5
Arap2
Elmod1
Heca
Cldn1
Mbtd1
Cadm1
Zfp532
Grhl3
Lrch1
Fam117a
Abhd17c
Gnai1
Psat1
Mmut
Selenoi
Tnfaip8
Stk10
Efna5
Ago1
Ppp2r5e
Aftph
Nckap1
Flrt3
Pik3cb
Gpr87
Zbtb16
Pex16
Ndor1
Hoxa5
Sema3f
Atpif1
Ly6g6e
Ppp1r13l
Fdps
Hr
Sun2
Tmem109
Cd300lg
Aldh3a1
Add3
Nudt17
Sult5a1
Ccl27a
Rab11fip4
Mccc2
Cbx4
Rnf167
Plch2
Aacs
Eps8l1
Llgl2
Ano9
Emp2
Tns4
Dhcr7
Paqr5
Rab3d
Chmp4c
Hsd17b14
Rdh1
Atp2b4
Pnkd
Lce1m
Il34
Acad9
Slc24a3
Fam241b
Krt23
Gprin2
Pdzd2
Cyfip2
Mfsd6
Ptpn14
Ptgs1
Notch3
Olfm2
Fam81a
Rarg
Phyhip
Acbd4
Wnt7b
Tjp3
Wnt3
Anxa8
Abhd14b
Dapl1
Cgn
Col17a1
Sh3d21
Abcc4
Cdc42ep1
Sphk1
Msh6
Pcdh7
Rcor1
Pcmtd2
Adtrp
Pof1b
Abca5
Clca2
Flg2
Lss
Etl4
Islr
Lhpp
Lrig1
Ust
Gramd4
Z−score
−2
0
2
WT
VM
WT
VM
WT
VM
WT
VM

#### Cluster4 ... Genes 1101 to 1200

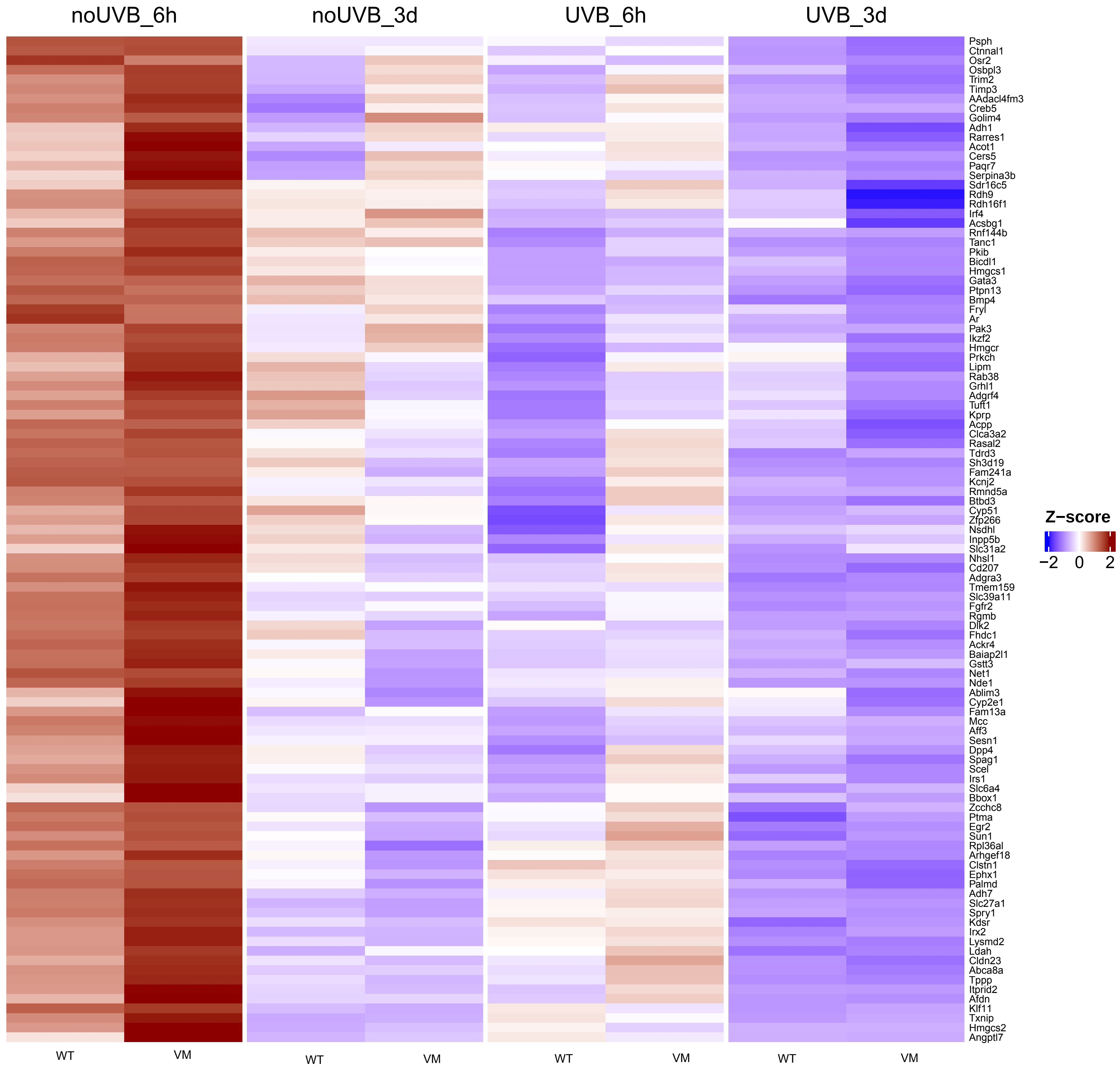

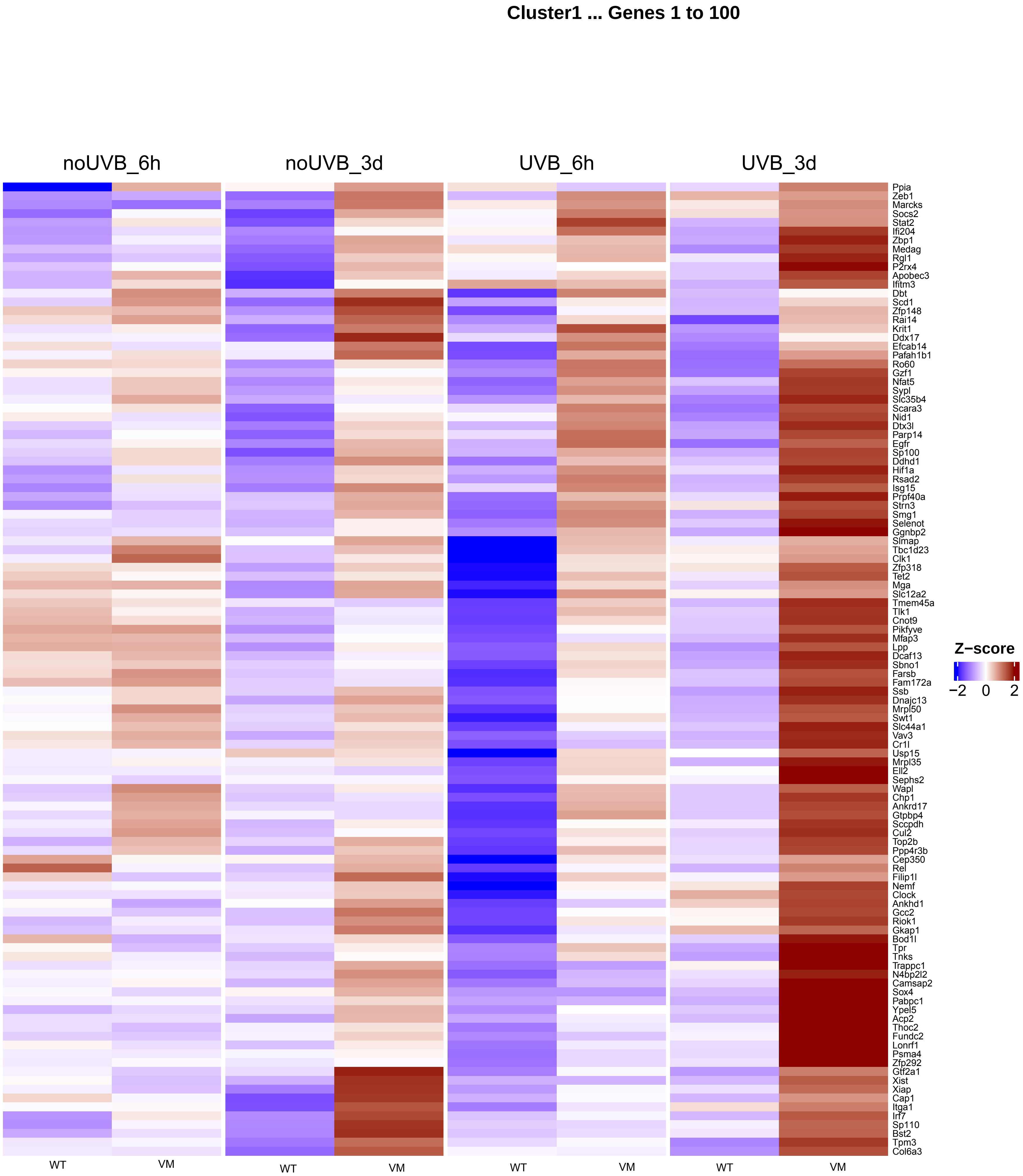

Cluster1 ... Genes 1 to 100
noUVB_6h
noUVB_3d
UVB_6h
UVB_3d
Ppia
Zeb1
Marcks
Socs2
Stat2
Ifi204
Zbp1
Medag
Rgl1
P2rx4
Apobec3
Ifitm3
Dbt
Scd1
Zfp148
Rai14
Krit1
Ddx17
Efcab14
Pafah1b1
Ro60
Gzf1
Nfat5
Sypl
Slc35b4
Scara3
Nid1
Dtx3l
Parp14
Egfr
Sp100
Ddhd1
Hif1a
Rsad2
Isg15
Prpf40a
Strn3
Smg1
Selenot
Ggnbp2
Slmap
Tbc1d23
Clk1
Zfp318
Tet2
Mga
Slc12a2
Tmem45a
Tlk1
Cnot9
Pikfyve
Mfap3
Lpp
Dcaf13
Sbno1
Farsb
Fam172a
Ssb
Dnajc13
Mrpl50
Swt1
Slc44a1
Vav3
Cr1l
Usp15
Mrpl35
Ell2
Sephs2
Wapl
Chp1
Ankrd17
Gtpbp4
Sccpdh
Cul2
Top2b
Ppp4r3b
Cep350
Rel
Filip1l
Nemf
Clock
Ankhd1
Gcc2
Riok1
Gkap1
Bod1l
Tpr
Tnks
Trappc1
N4bp2l2
Camsap2
Sox4
Pabpc1
Ypel5
Acp2
Thoc2
Fundc2
Lonrf1
Psma4
Zfp292
Gtf2a1
Xist
Xiap
Cap1
Itga1
Irf7
Sp110
Bst2
Tpm3
Col6a3
Z−score
−2 0 2
WT
VM
WT
VM
WT
VM
WT
VM

**Cluster1 ... Genes 111 to 210**

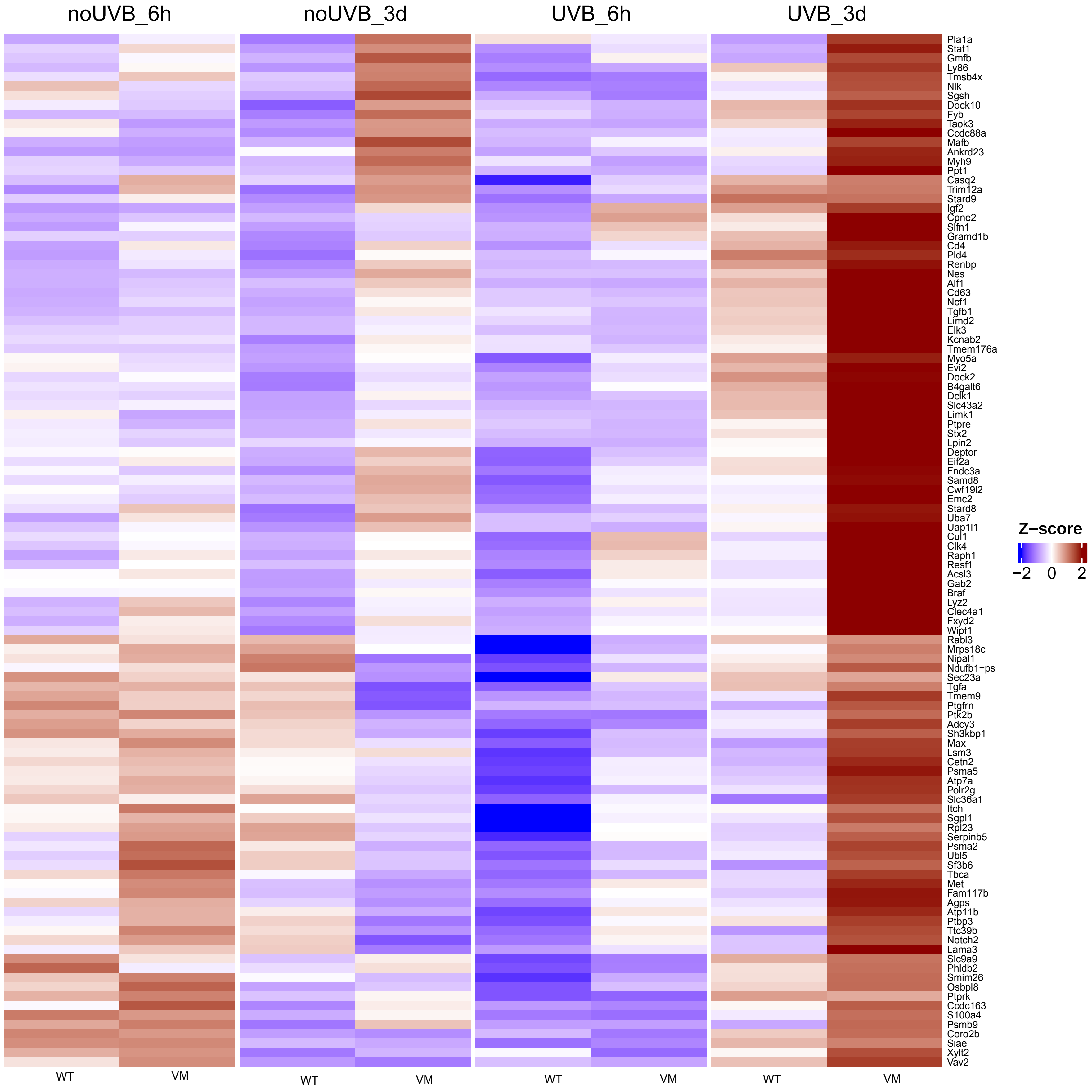

#### Cluster1 ... Genes 221 to 320

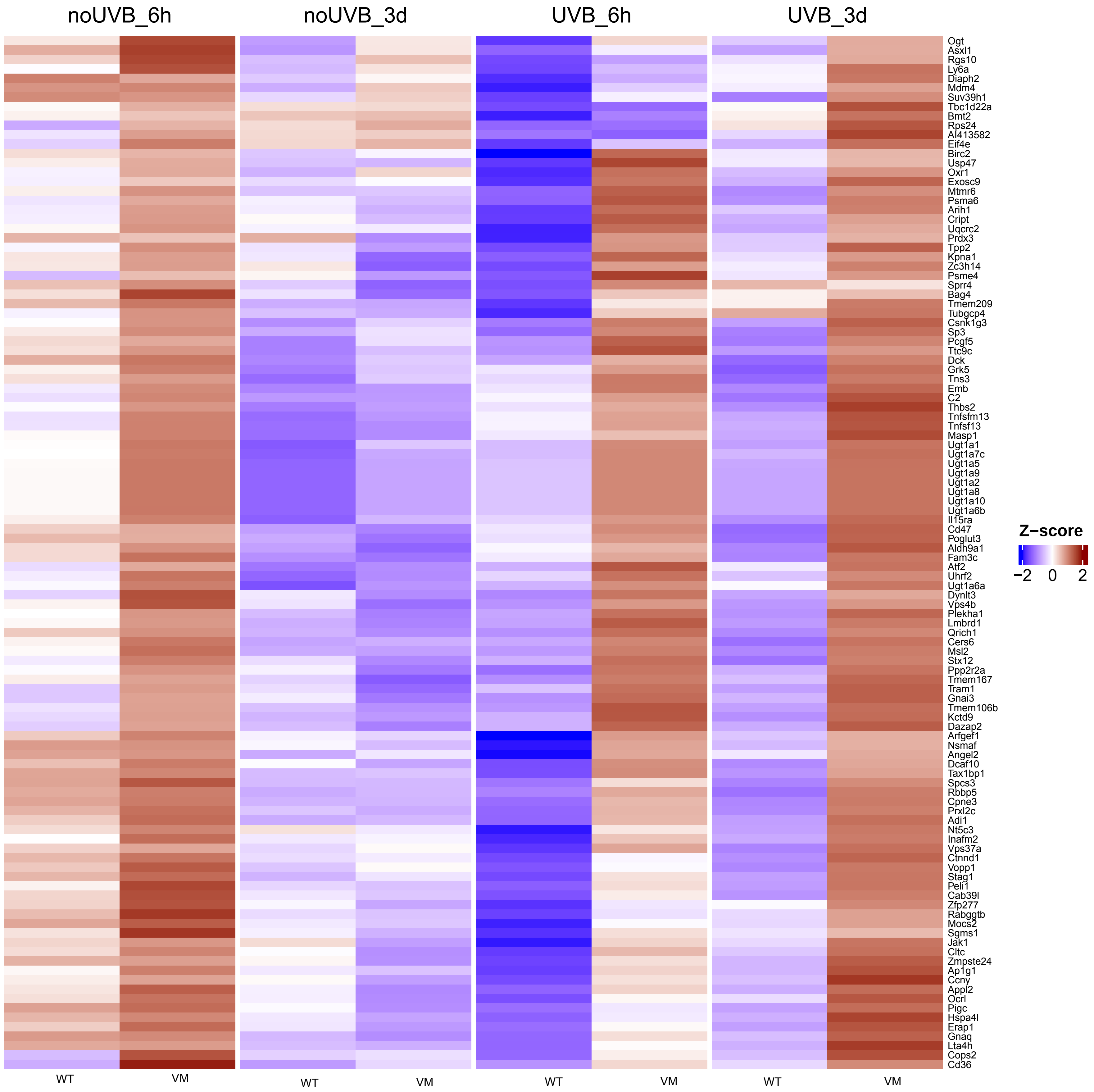

**Cluster1 ... Genes 331 to 430**

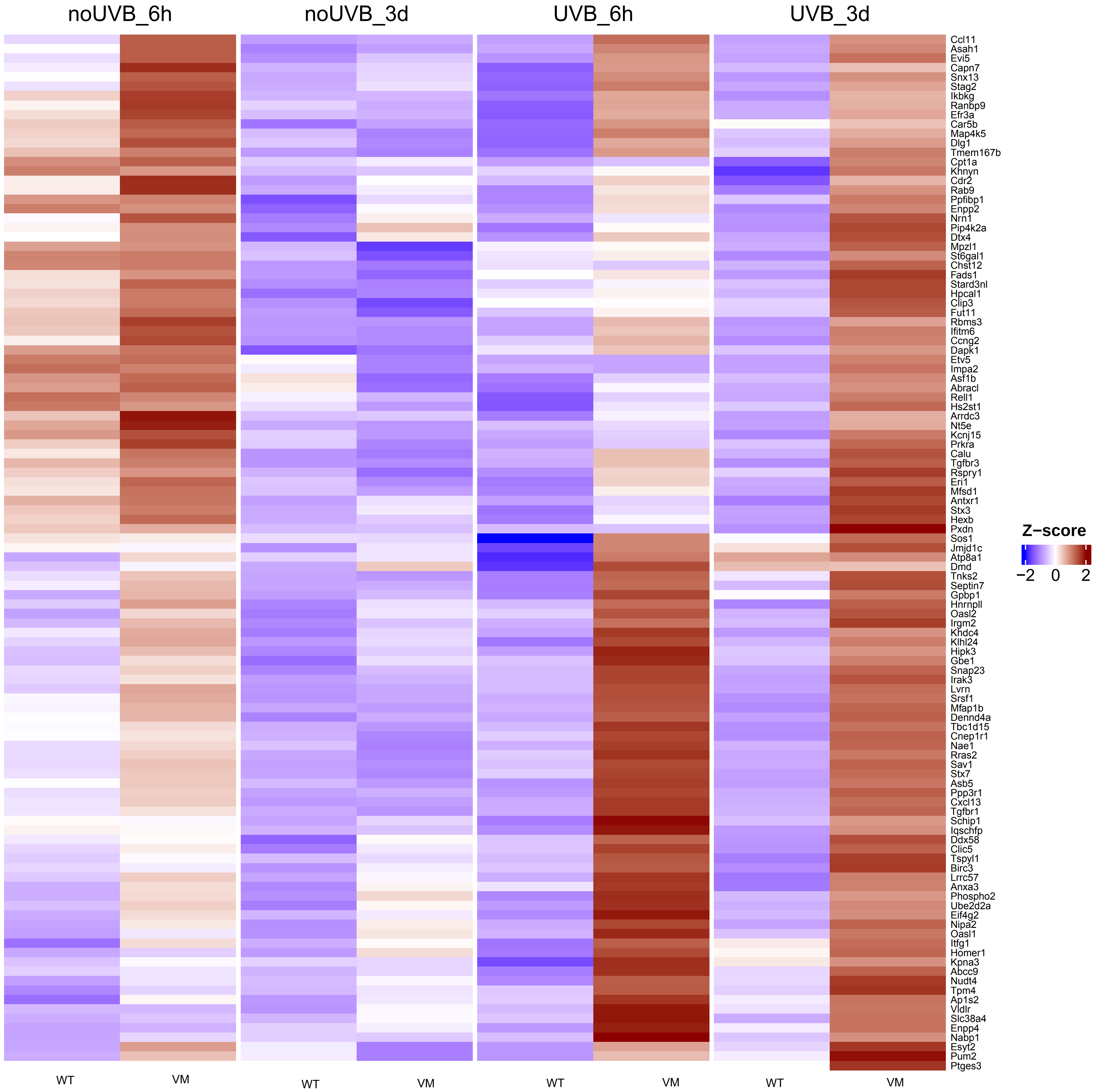

Cluster1 ... Genes 441 to 540

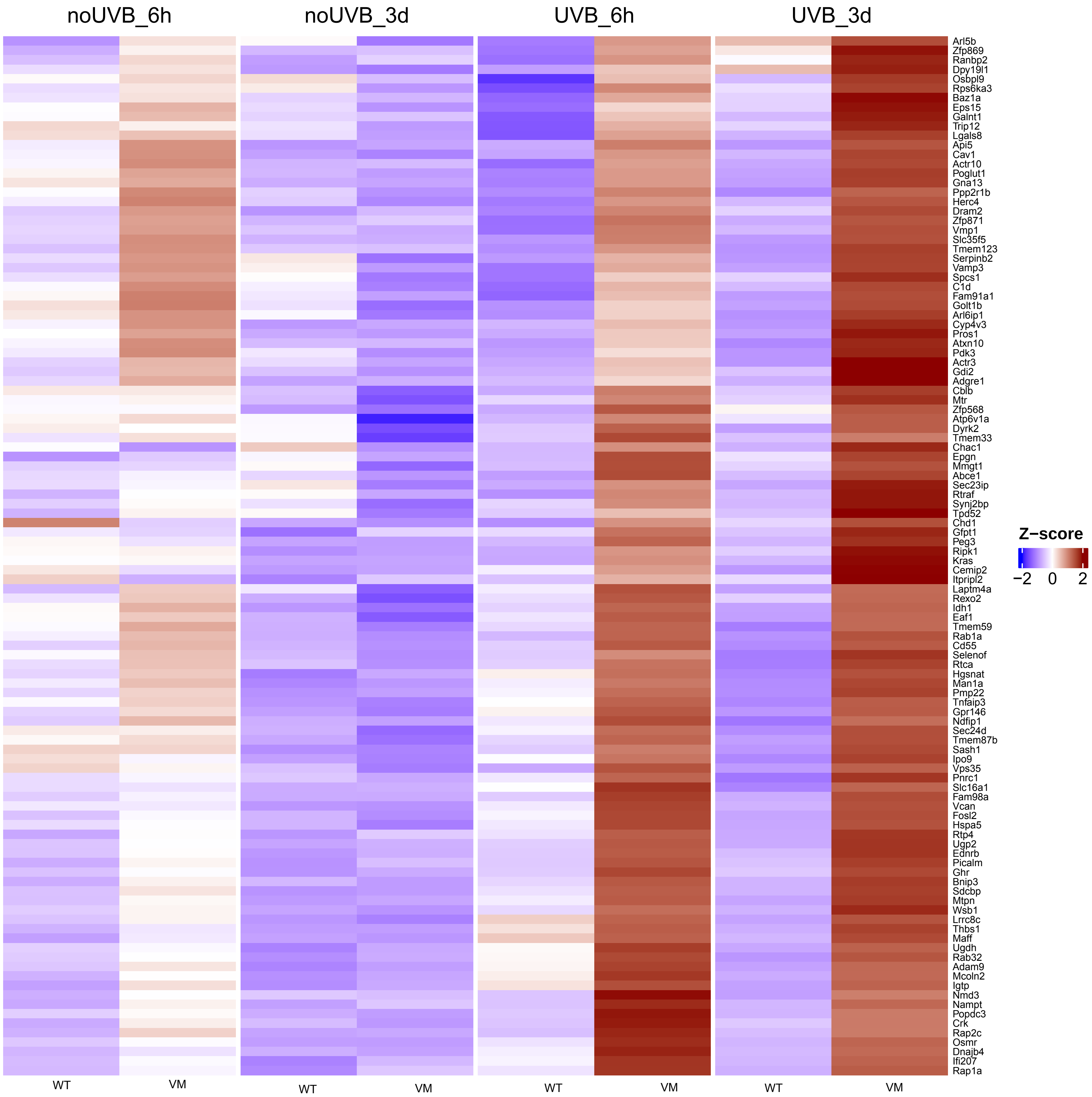

**Cluster1 ... Genes 551 to 650**

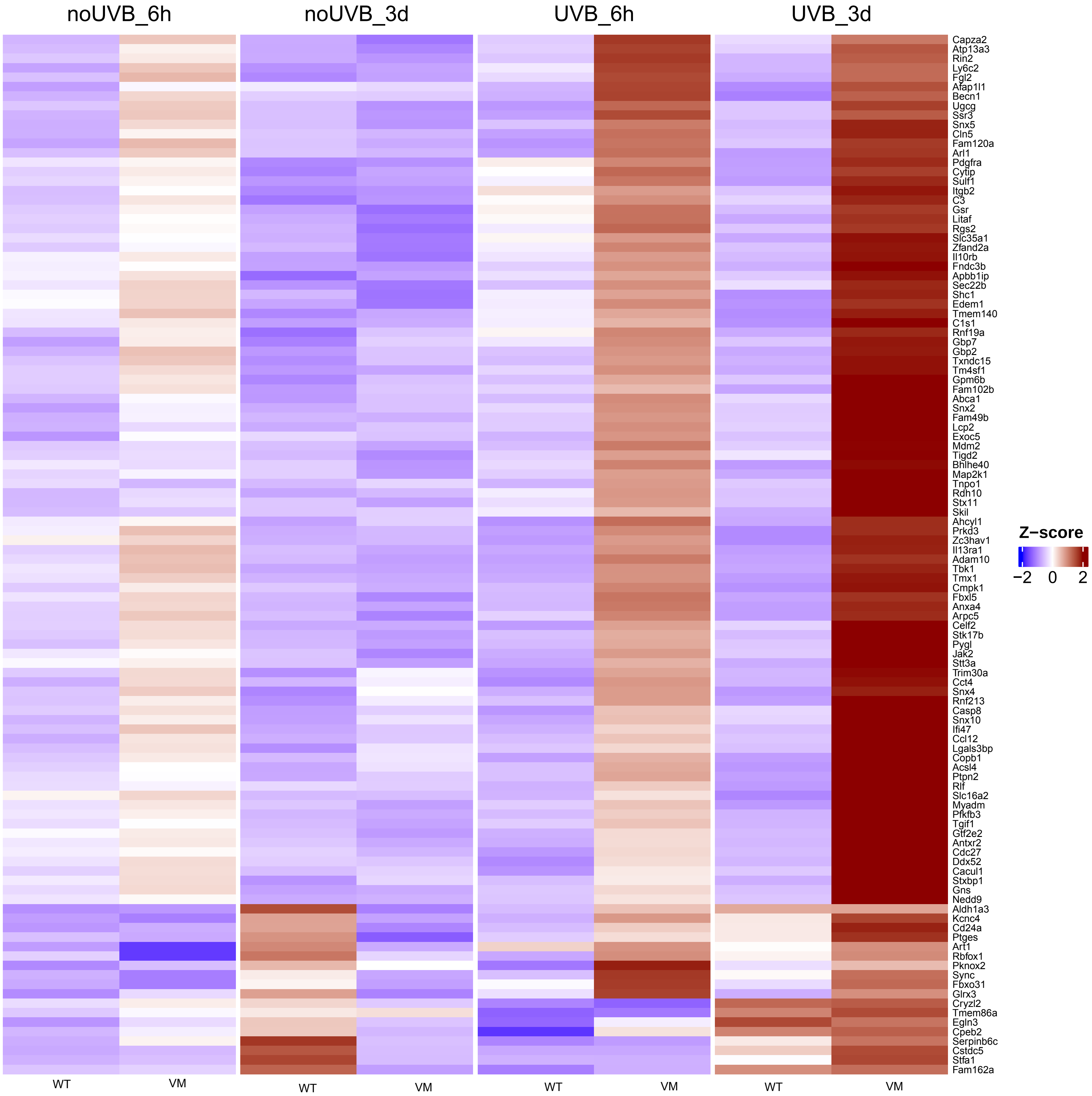

**Cluster1 ... Genes 661 to 760**

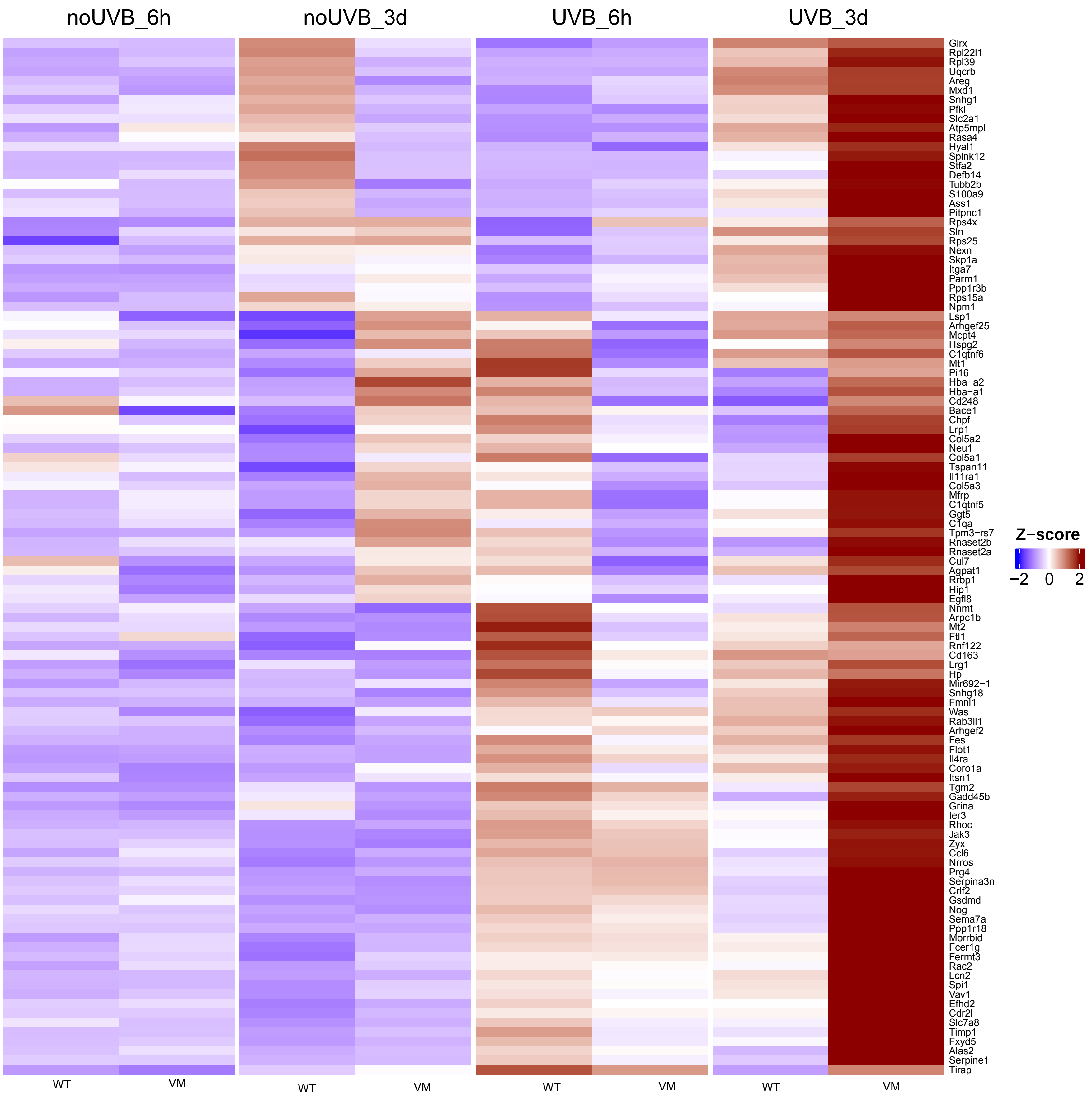

**Cluster1 ... Genes 771 to 870**

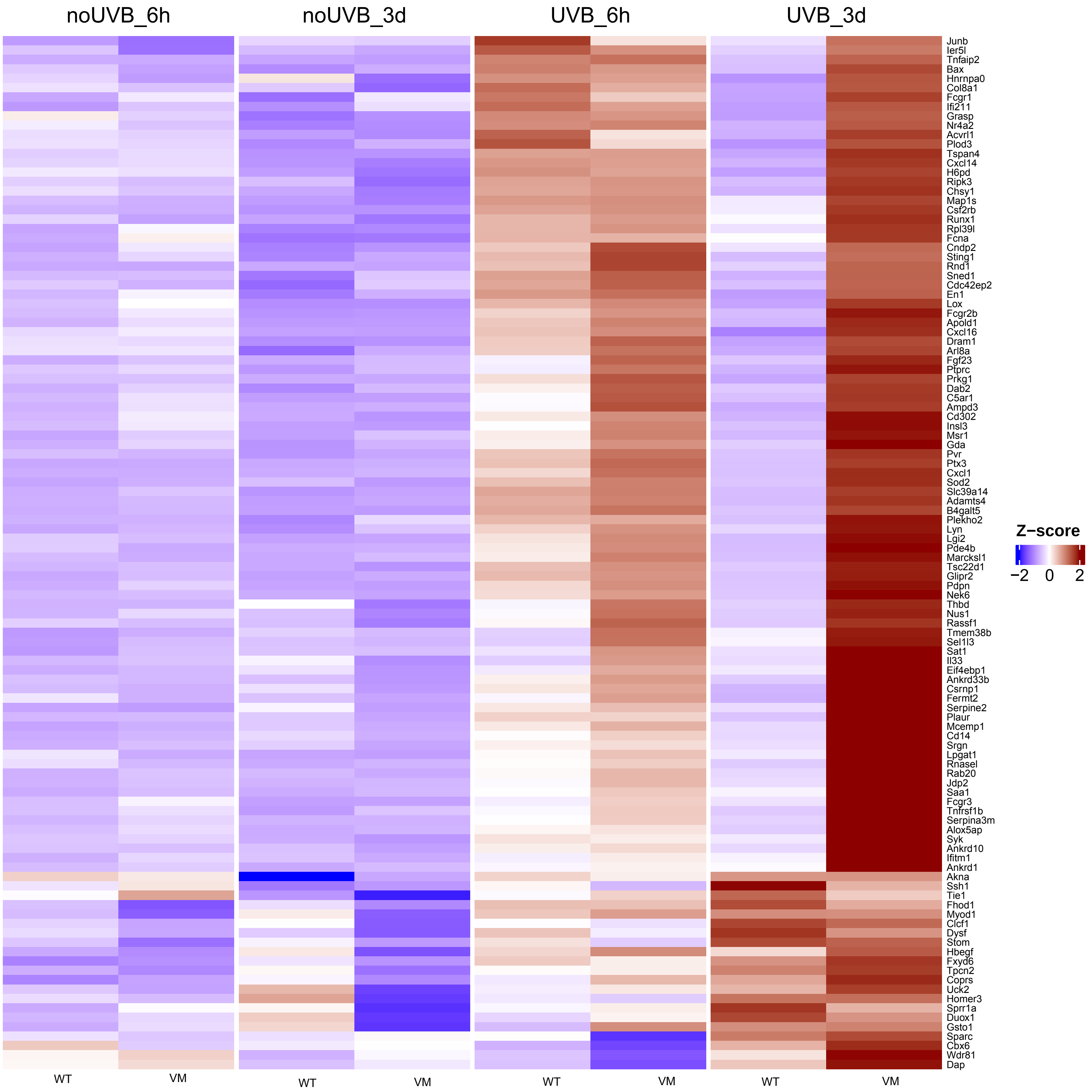

Cluster1 ... Genes 881 to 980

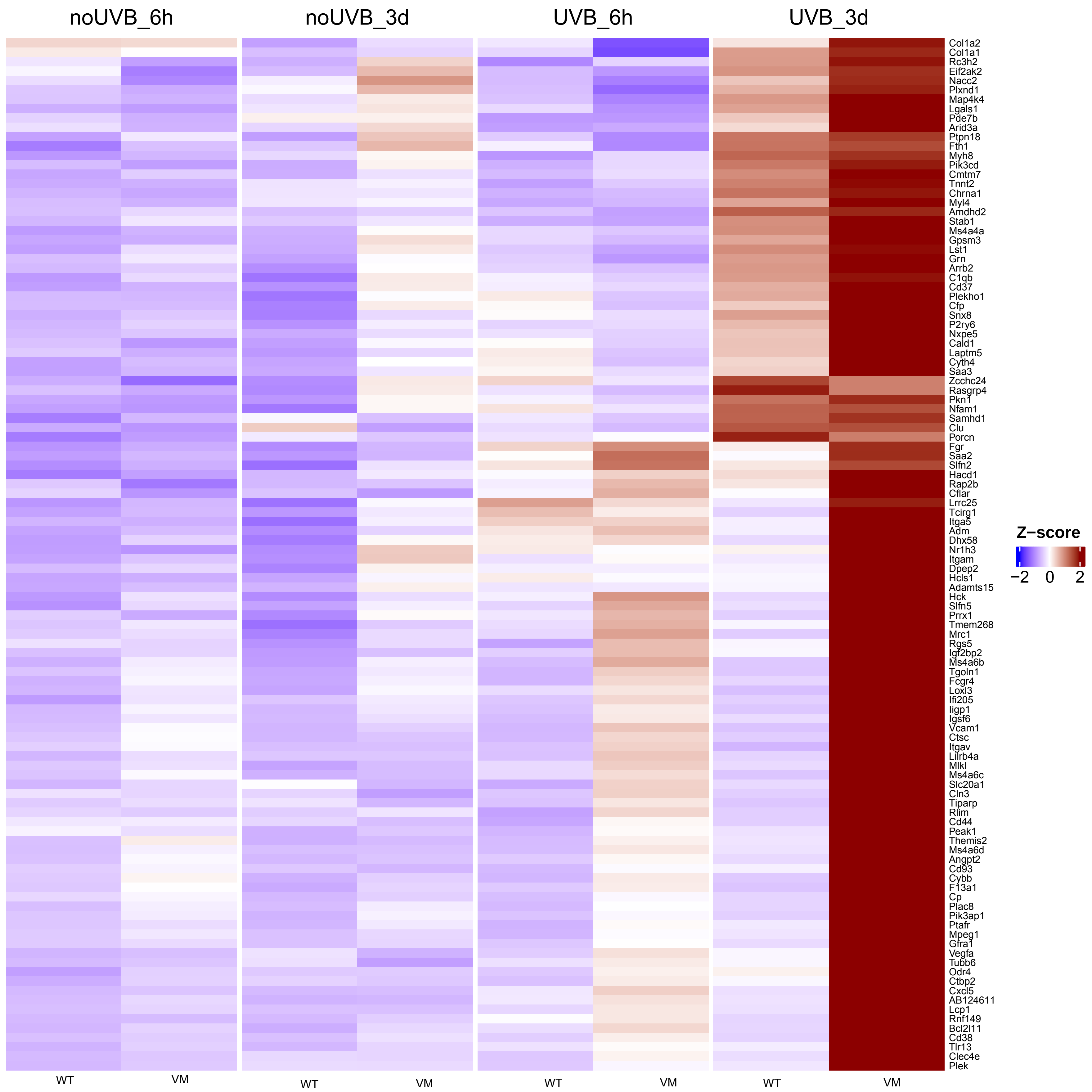

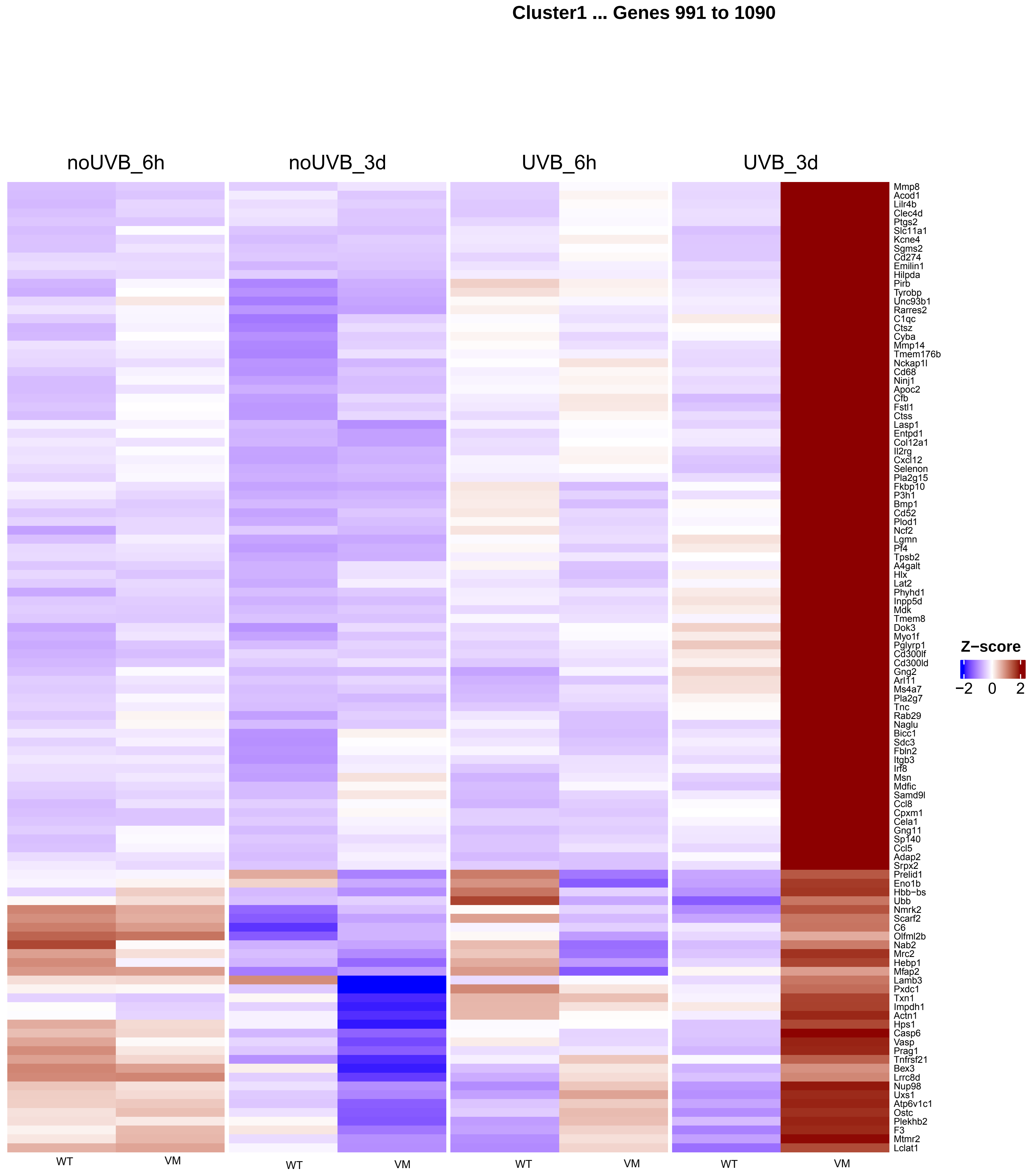
Cluster1 ... Genes 991 to 1090
noUVB_6h
noUVB_3d
UVB_6h
UVB_3d
Mmp8
Acod1
Lilr4b
Clec4d
Ptgs2
Slc11a1
Kcne4
Sgms2
Cd274
Emilin1
Hilpda
Pirb
Tyrobp
Unc93b1
Rarres2
C1qc
Ctsz
Cyba
Mmp14
Tmem176b
Nckap1l
Cd68
Ninj1
Apoc2
Cfb
Fstl1
Ctss
Lasp1
Entpd1
Col12a1
Il2rg
Cxcl12
Selenon
Pla2g15
Fkbp10
P3h1
Bmp1
Cd52
Plod1
Ncf2
Lgmn
Pf4
Tpsb2
A4galt
Hlx
Lat2
Phyhd1
Inpp5d
Mdk
Tmem8
Dok3
Myo1f
Pglyrp1
Cd300lf
Cd300ld
Gng2
Arl11
Ms4a7
Pla2g7
Tnc
Rab29
Naglu
Bicc1
Sdc3
Fbln2
Itgb3
Irf8
Msn
Mdfic
Samd9l
Ccl8
Cpxm1
Cela1
Gng11
Sp140
Ccl5
Adap2
Srpx2
Prelid1
Eno1b
Hbb−bs
Ubb
Nmrk2
Scarf2
C6
Olfml2b
Nab2
Mrc2
Hebp1
Mfap2
Lamb3
Pxdc1
Txn1
Impdh1
Actn1
Hps1
Casp6
Vasp
Prag1
Tnfrsf21
Bex3
Lrrc8d
Nup98
Uxs1
Atp6v1c1
Ostc
Plekhb2
F3
Mtmr2
Lclat1
Z−score
−2
0
2
WT
VM
WT
VM
WT
VM
WT
VM

**Cluster1 ... Genes 1101 to 1200**

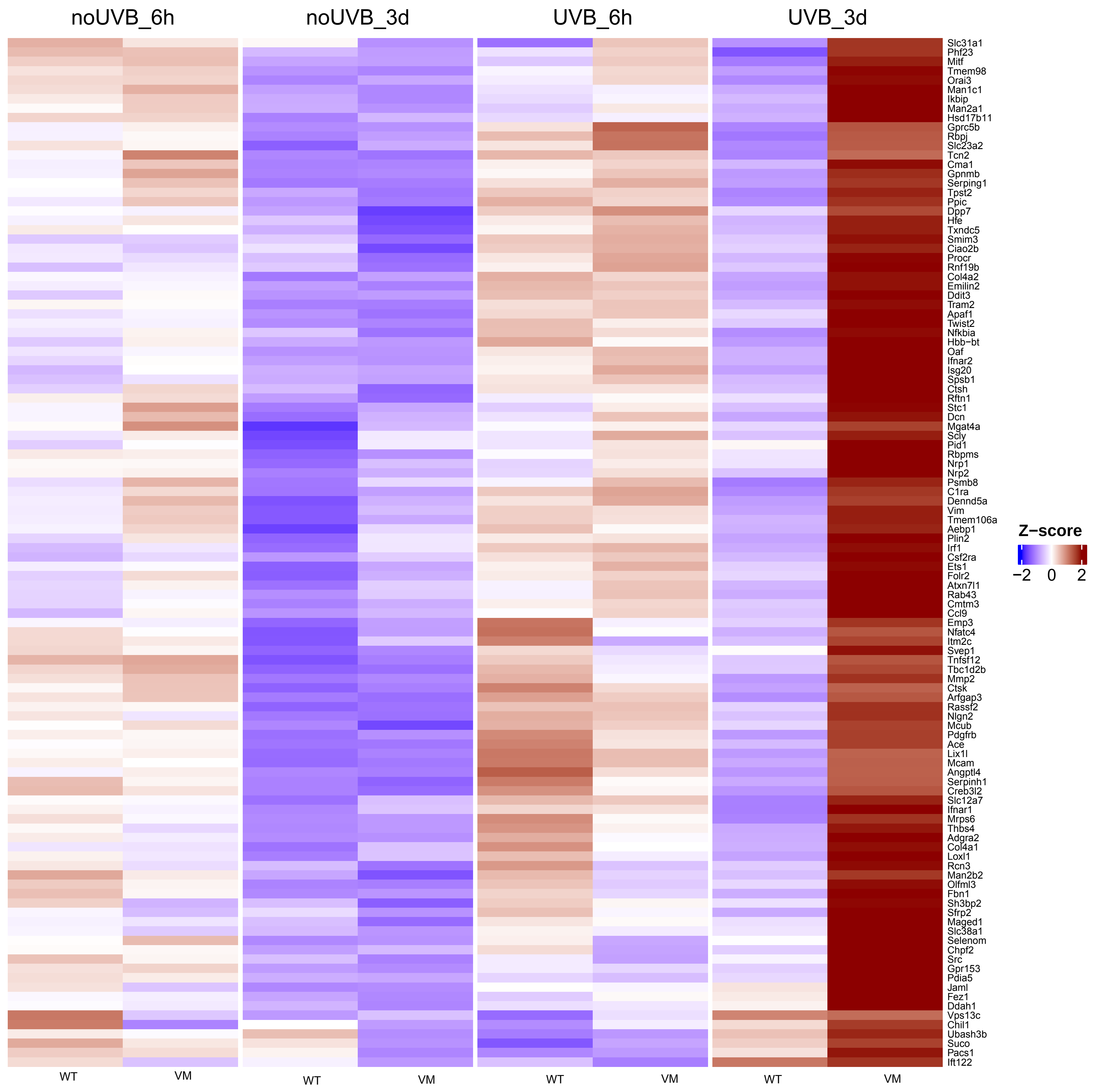

#### Cluster1 ... Genes 1211 to 1310

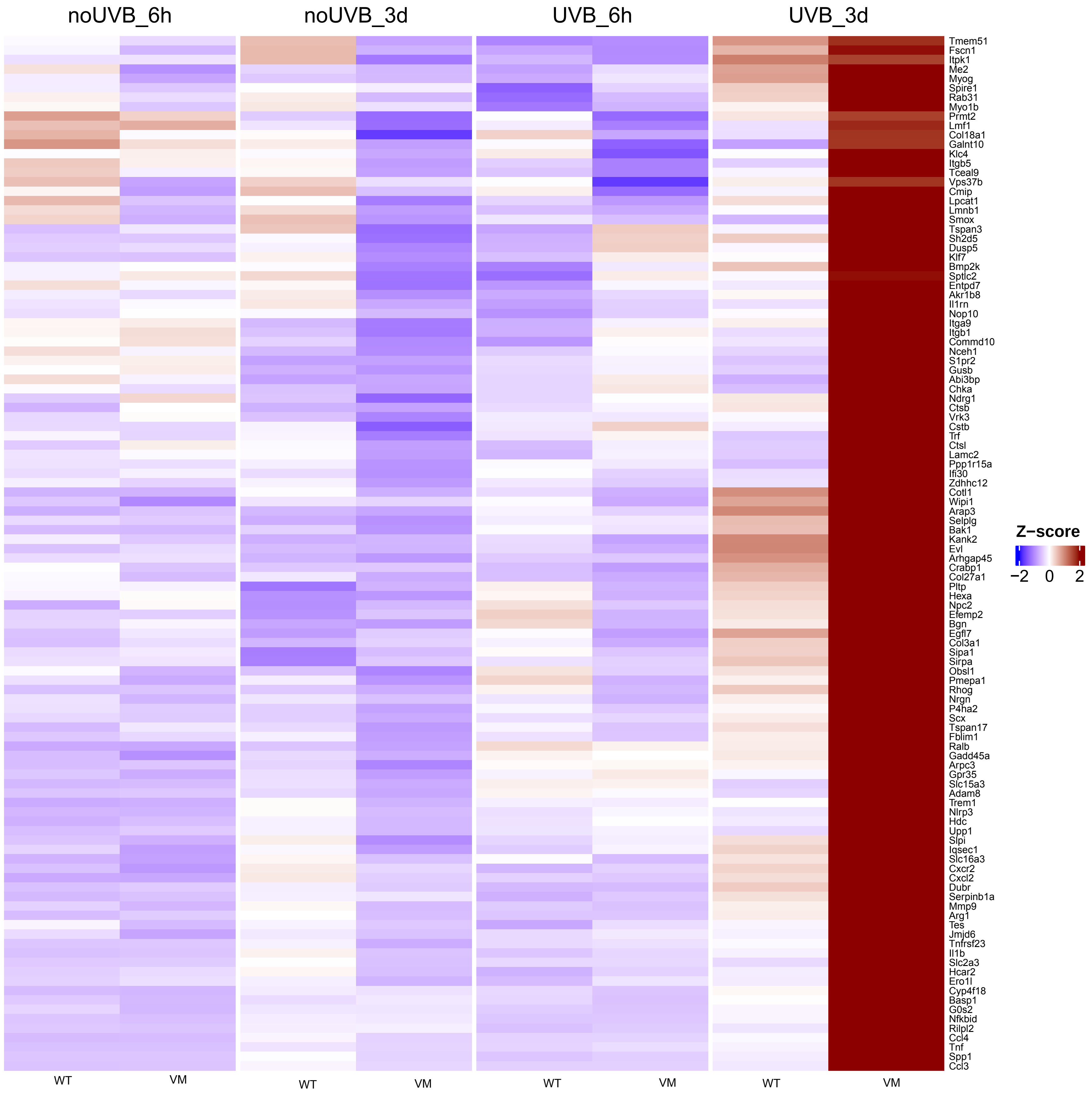

**Cluster1 ... Genes 1321 to 1420**

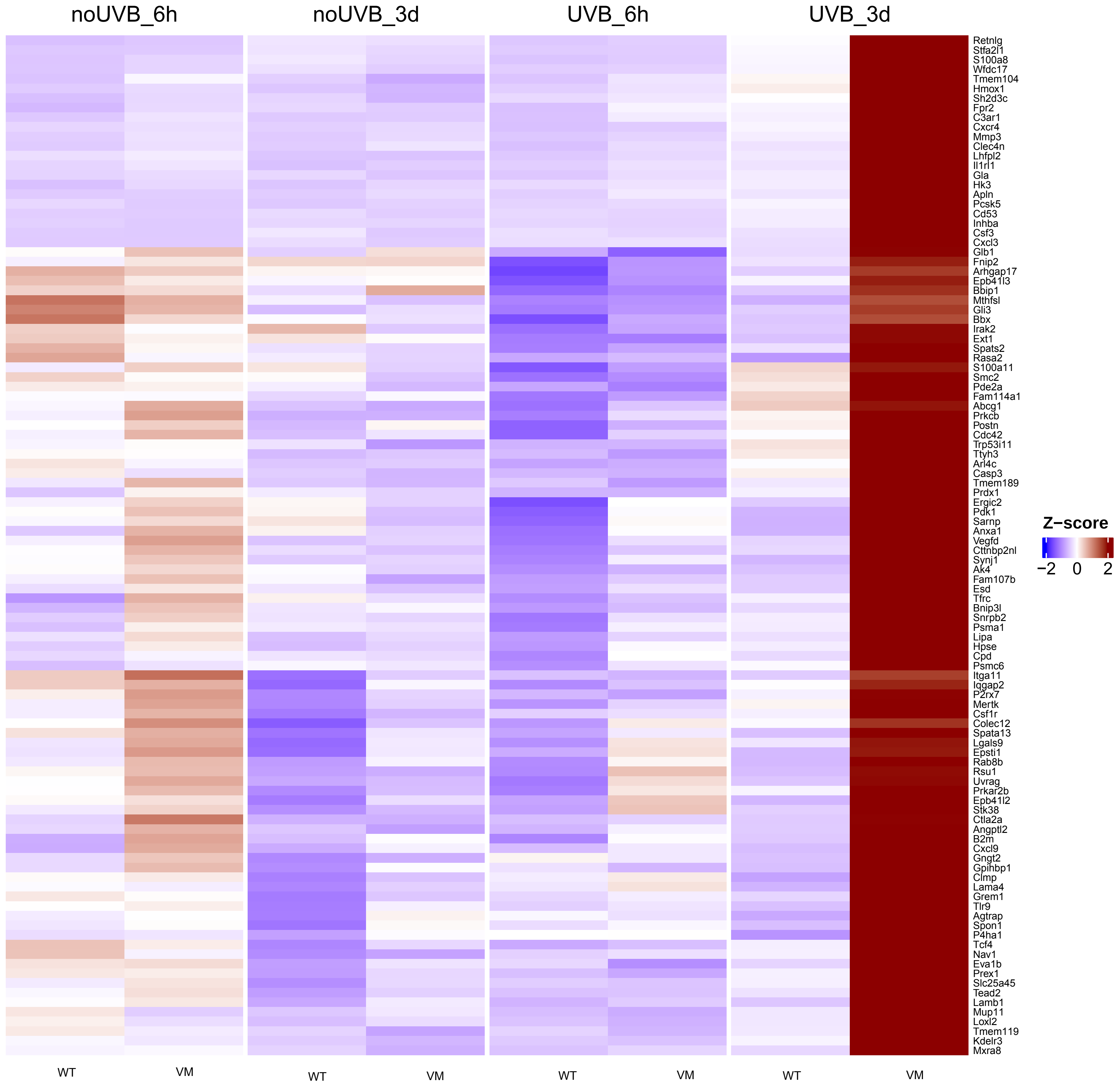

**Cluster2 ... Genes 1 to 100**

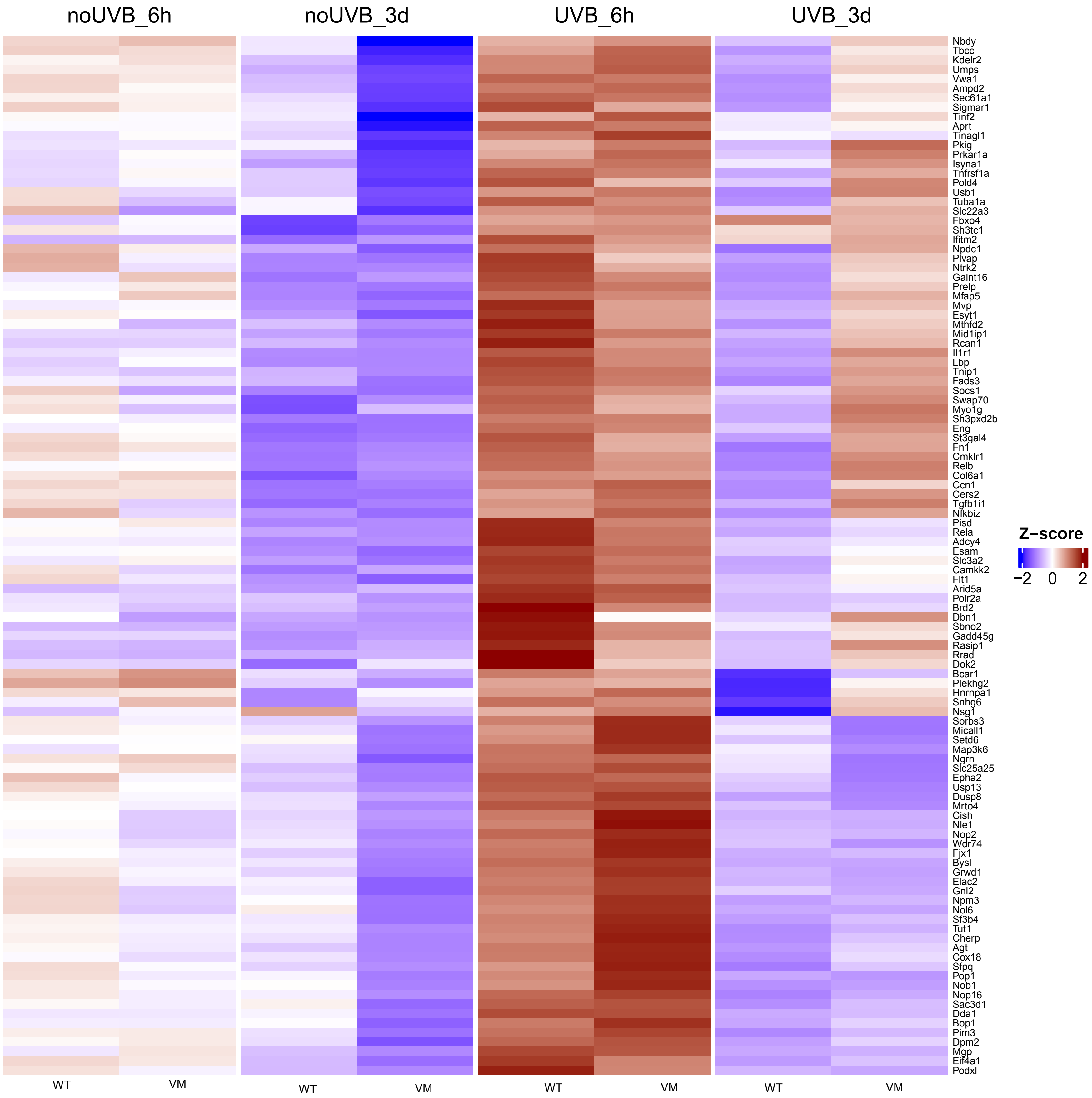

**Cluster2 ... Genes 111 to 210**

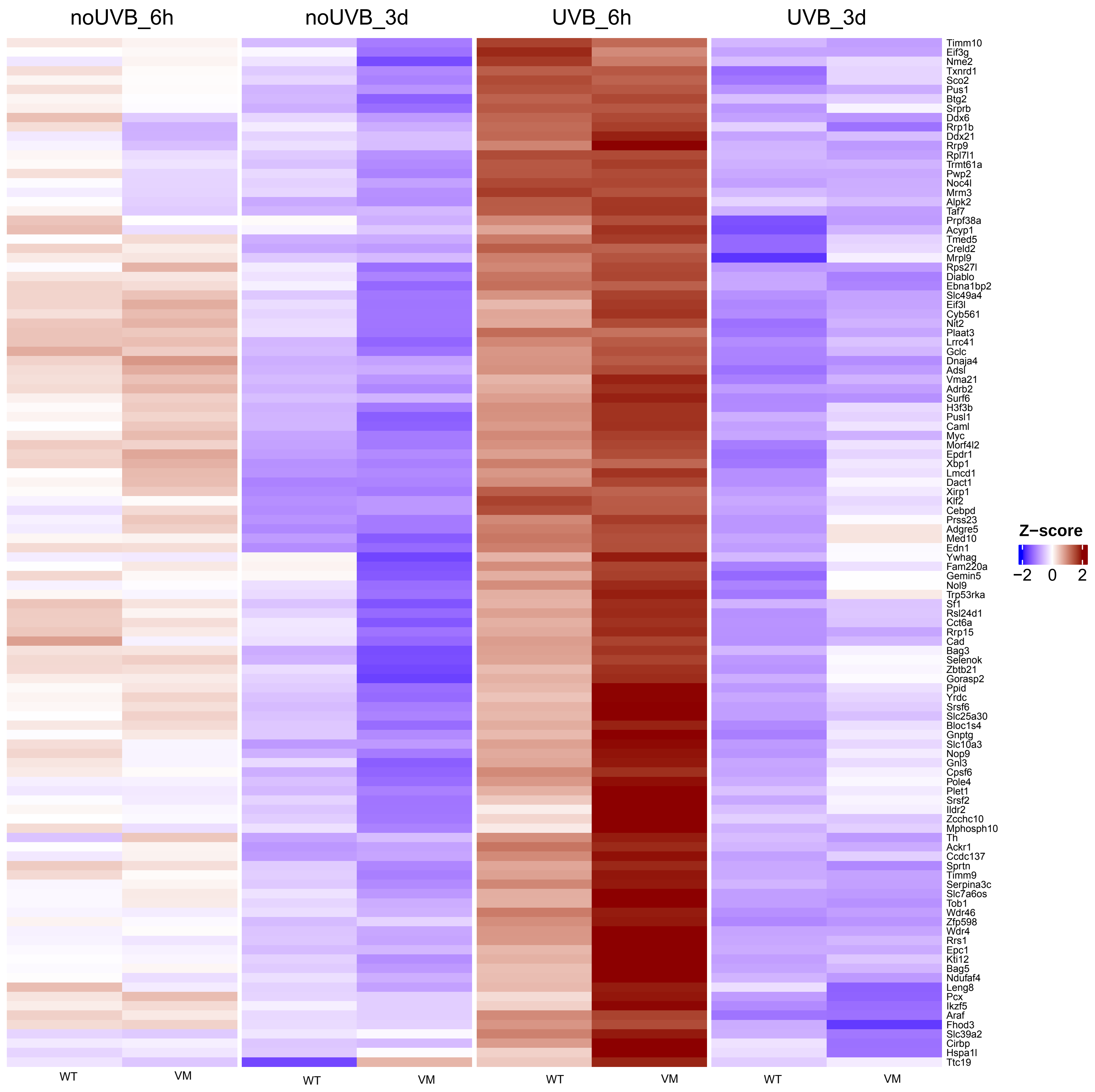

Cluster2 ... Genes 221 to 320

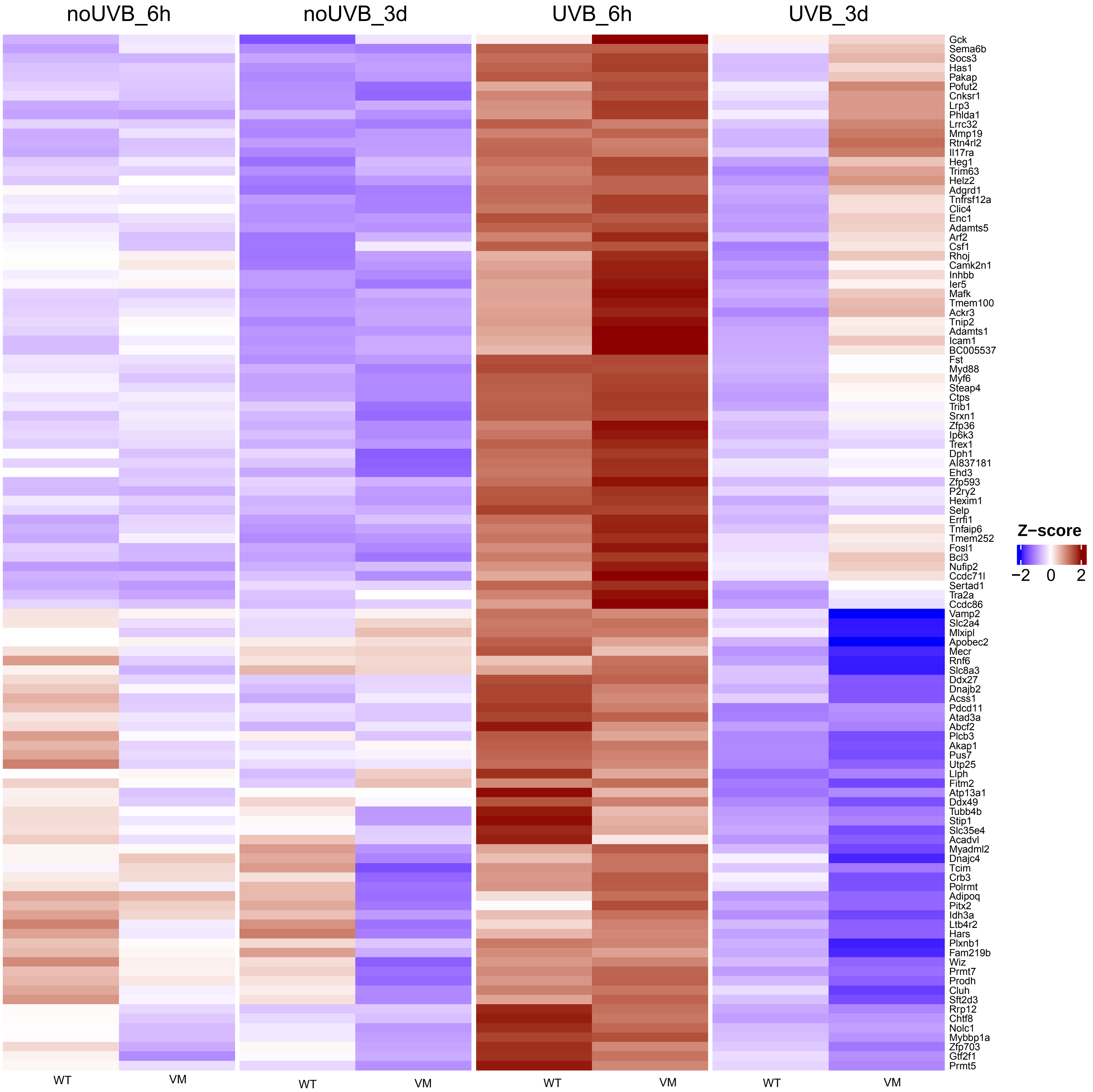

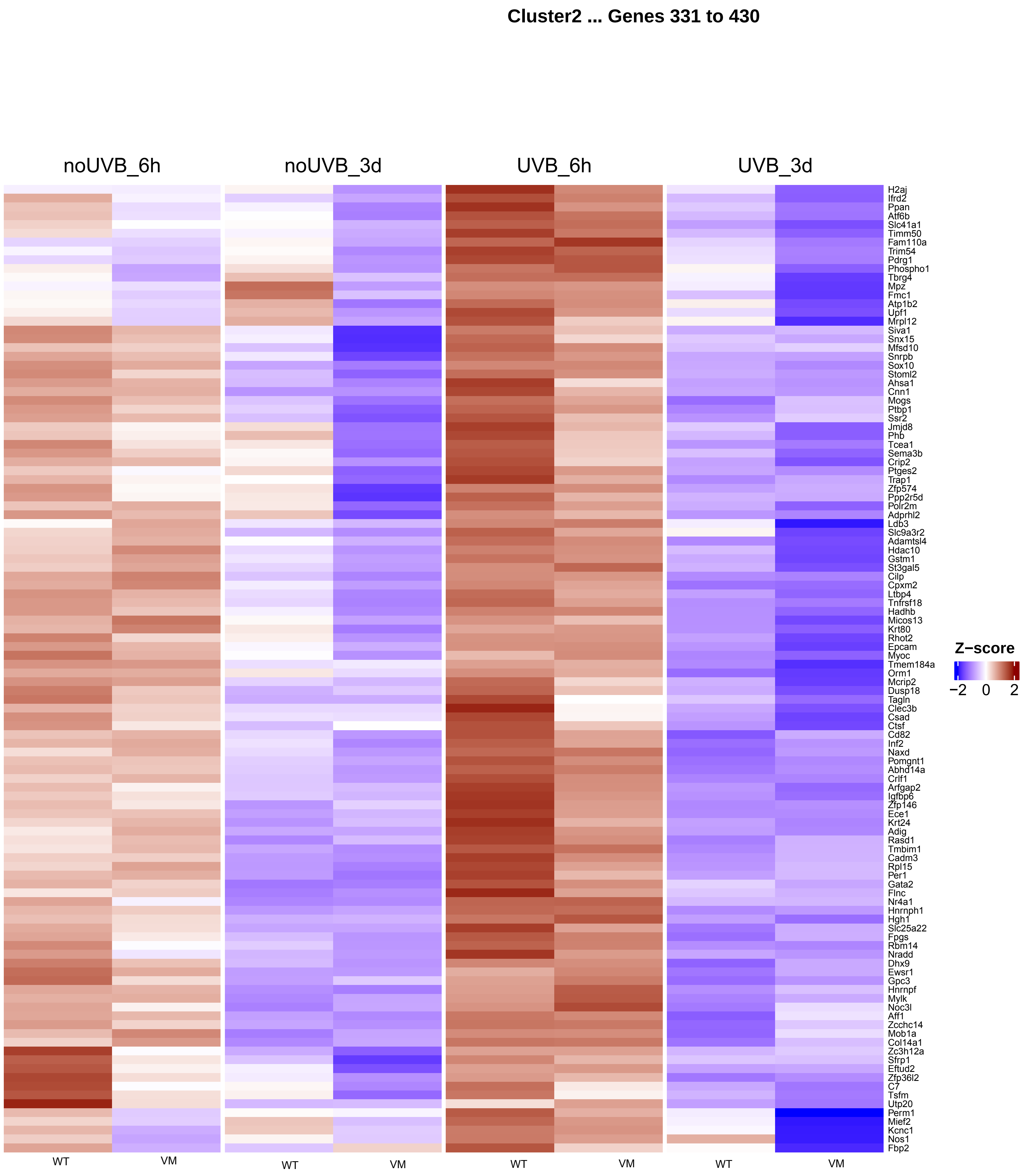
Cluster2 ... Genes 331 to 430
noUVB_6h
noUVB_3d
UVB_6h
UVB_3d
H2aj
Ifrd2
Ppan
Atf6b
Slc41a1
Timm50
Fam110a
Trim54
Pdrg1
Phospho1
Tbrg4
Mpz
Fmc1
Atp1b2
Upf1
Mrpl12
Siva1
Snx15
Mfsd10
Snrpb
Sox10
Stoml2
Ahsa1
Cnn1
Mogs
Ptbp1
Ssr2
Jmjd8
Phb
Tcea1
Sema3b
Crip2
Ptges2
Trap1
Zfp574
Ppp2r5d
Polr2m
Adprhl2
Ldb3
Slc9a3r2
Adamtsl4
Hdac10
Gstm1
St3gal5
Cilp
Cpxm2
Ltbp4
Tnfrsf18
Hadhb
Micos13
Krt80
Rhot2
Epcam
Myoc
Tmem184a
Orm1
Mcrip2
Dusp18
Tagln
Clec3b
Csad
Ctsf
Cd82
Inf2
Naxd
Pomgnt1
Abhd14a
Crlf1
Arfgap2
Igfbp6
Zfp146
Ece1
Krt24
Adig
Rasd1
Tmbim1
Cadm3
Rpl15
Per1
Gata2
Flnc
Nr4a1
Hnrnph1
Hgh1
Slc25a22
Fpgs
Rbm14
Nradd
Dhx9
Ewsr1
Gpc3
Hnrnpf
Mylk
Noc3l
Aff1
Zcchc14
Mob1a
Col14a1
Zc3h12a
Sfrp1
Eftud2
Zfp36l2
C7
Tsfm
Utp20
Perm1
Mief2
Kcnc1
Nos1
Fbp2
WT
VM
WT
VM
WT
VM
WT
VM
Z−score
−2
0
2

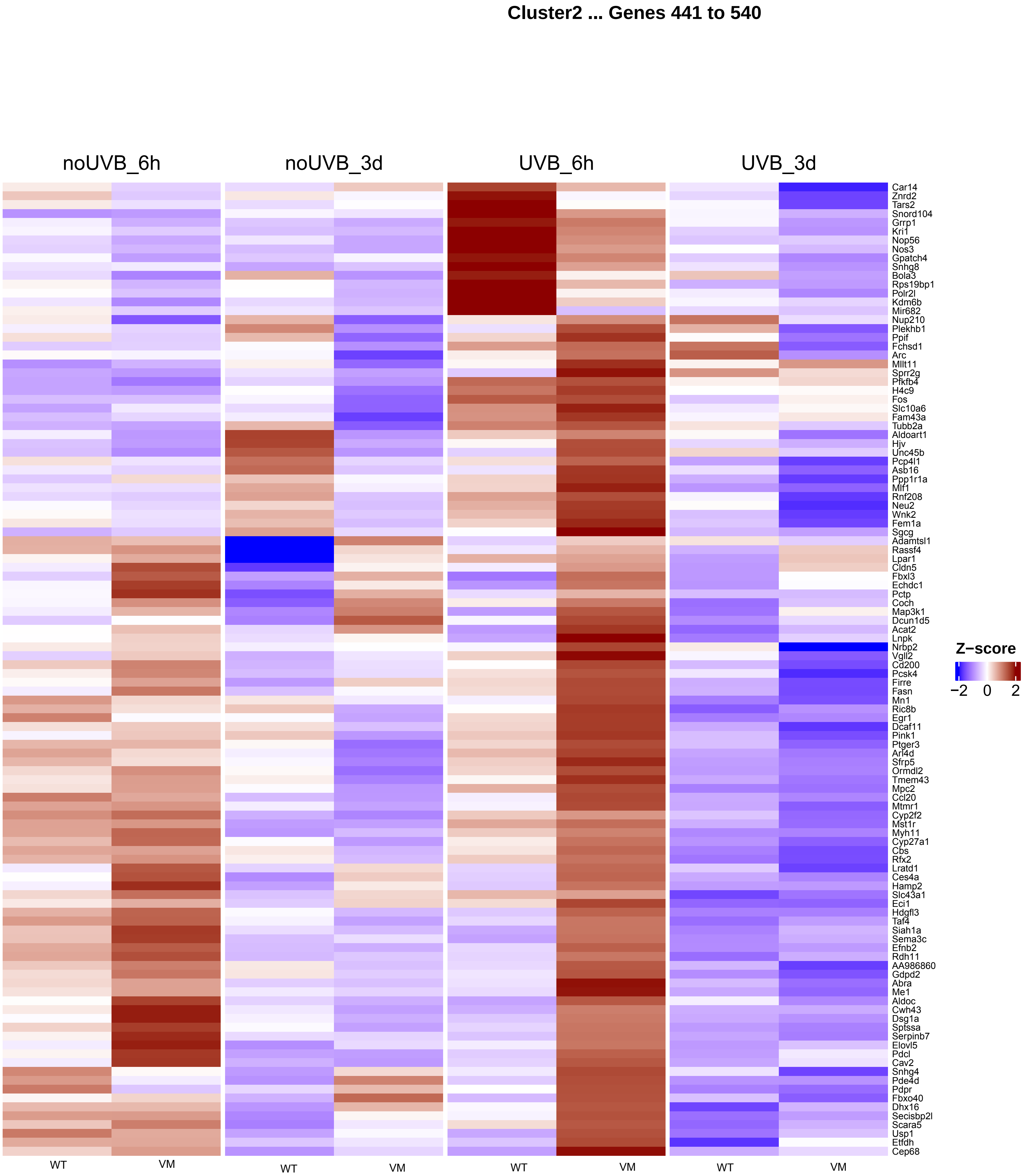
Cluster2 ... Genes 441 to 540
noUVB_6h
noUVB_3d
UVB_6h
UVB_3d
Car14
Znrd2
Tars2
Snord104
Grrp1
Kri1
Nop56
Nos3
Gpatch4
Snhg8
Bola3
Rps19bp1
Polr2l
Kdm6b
Mir682
Nup210
Plekhb1
Ppif
Fchsd1
Arc
Mllt11
Sprr2g
Pfkfb4
H4c9
Fos
Slc10a6
Fam43a
Tubb2a
Aldoart1
Hjv
Unc45b
Pcp4l1
Asb16
Ppp1r1a
Mlf1
Rnf208
Neu2
Wnk2
Fem1a
Sgcg
Adamtsl1
Rassf4
Lpar1
Cldn5
Fbxl3
Echdc1
Pctp
Coch
Map3k1
Dcun1d5
Acat2
Lnpk
Nrbp2
Vgll2
Cd200
Pcsk4
Firre
Fasn
Mn1
Ric8b
Egr1
Dcaf11
Pink1
Ptger3
Arl4d
Sfrp5
Ormdl2
Tmem43
Mpc2
Ccl20
Mtmr1
Cyp2f2
Mst1r
Myh11
Cyp27a1
Cbs
Rfx2
Lratd1
Ces4a
Hamp2
Slc43a1
Eci1
Hdgfl3
Taf4
Siah1a
Sema3c
Efnb2
Rdh11
AA986860
Gdpd2
Abra
Me1
Aldoc
Cwh43
Dsg1a
Sptssa
Serpinb7
Elovl5
Pdcl
Cav2
Snhg4
Pde4d
Pdpr
Fbxo40
Dhx16
Secisbp2l
Scara5
Usp1
Etfdh
Cep68
Z−score
−2
0
2
WT
VM
WT
VM
WT
VM
WT
VM

Cluster2 ... Genes 551 to 650

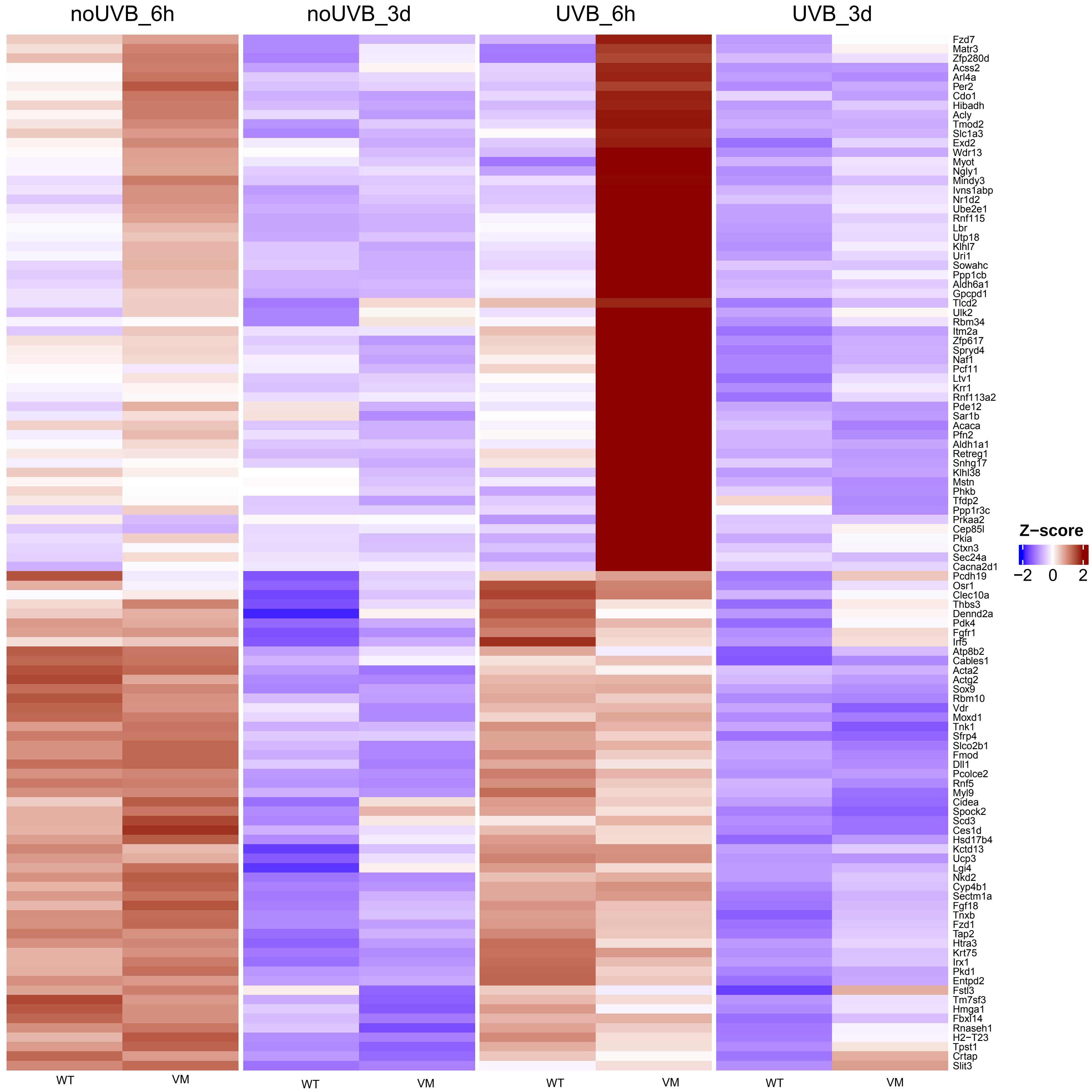

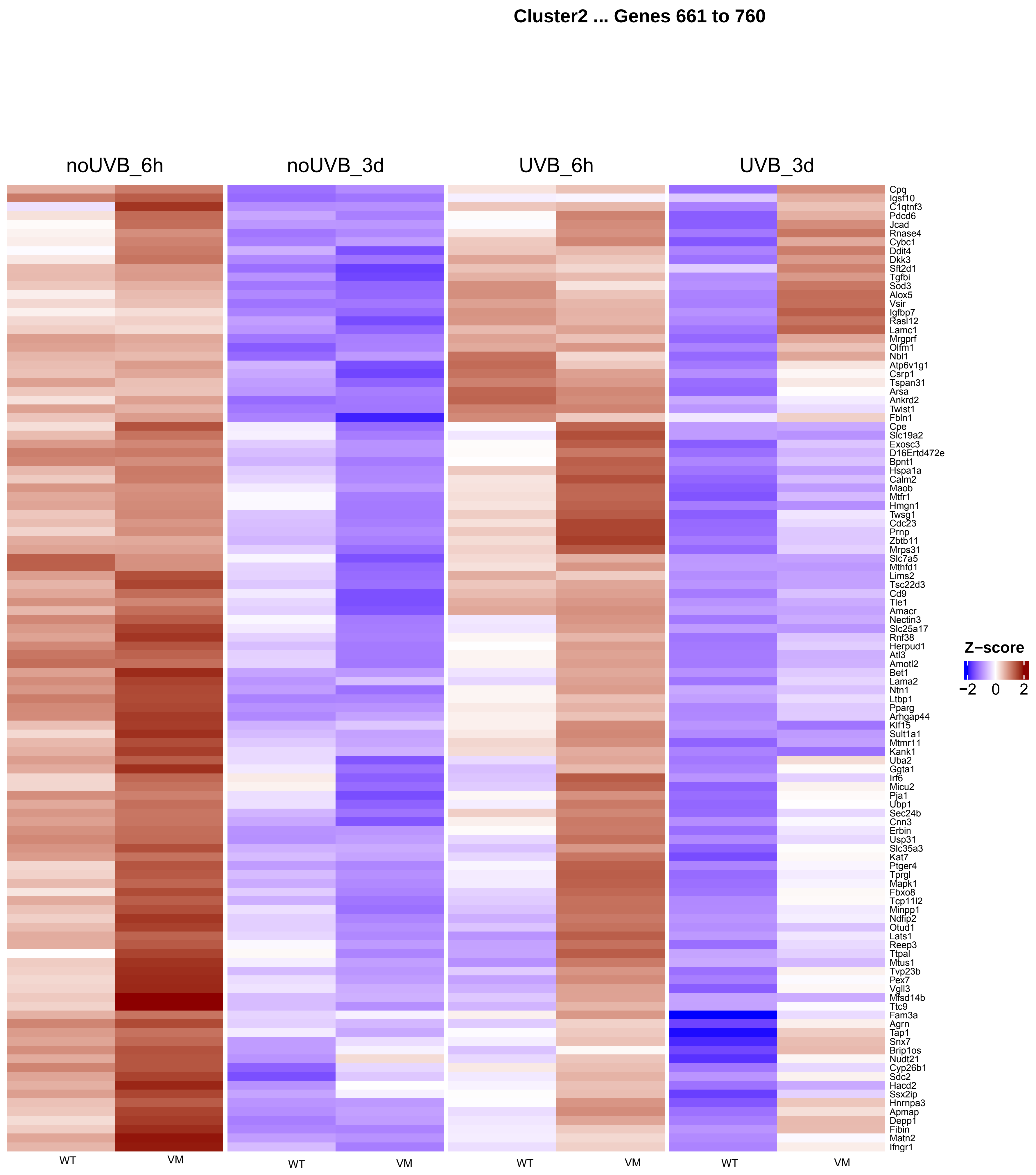
Cluster2 ... Genes 661 to 760
noUVB_6h
noUVB_3d
UVB_6h
UVB_3d
Cpq
Igsf10
C1qtnf3
Pdcd6
Jcad
Rnase4
Cybc1
Ddit4
Dkk3
Sft2d1
Tgfbi
Sod3
Alox5
Vsir
Igfbp7
Rasl12
Lamc1
Mrgprf
Olfm1
Nbl1
Atp6v1g1
Csrp1
Tspan31
Arsa
Ankrd2
Twist1
Fbln1
Cpe
Slc19a2
Exosc3
D16Ertd472e
Bpnt1
Hspa1a
Calm2
Maob
Mtfr1
Hmgn1
Twsg1
Cdc23
Prnp
Zbtb11
Mrps31
Slc7a5
Mthfd1
Lims2
Tsc22d3
Cd9
Tle1
Amacr
Nectin3
Slc25a17
Rnf38
Herpud1
Atl3
Amotl2
Bet1
Lama2
Ntn1
Ltbp1
Pparg
Arhgap44
Klf15
Sult1a1
Mtmr11
Kank1
Uba2
Ggta1
Irf6
Micu2
Pja1
Ubp1
Sec24b
Cnn3
Erbin
Usp31
Slc35a3
Kat7
Ptger4
Tprgl
Mapk1
Fbxo8
Tcp11l2
Minpp1
Ndfip2
Otud1
Lats1
Reep3
Ttpal
Mtus1
Tvp23b
Pex7
Vgll3
Mfsd14b
Ttc9
Fam3a
Agrn
Tap1
Snx7
Brip1os
Nudt21
Cyp26b1
Sdc2
Hacd2
Ssx2ip
Hnrnpa3
Apmap
Depp1
Fibin
Matn2
Ifngr1
Z−score
−2 0 2
WT
VM
WT
VM
WT
VM
WT
VM

**Cluster2 ... Genes 771 to 870**

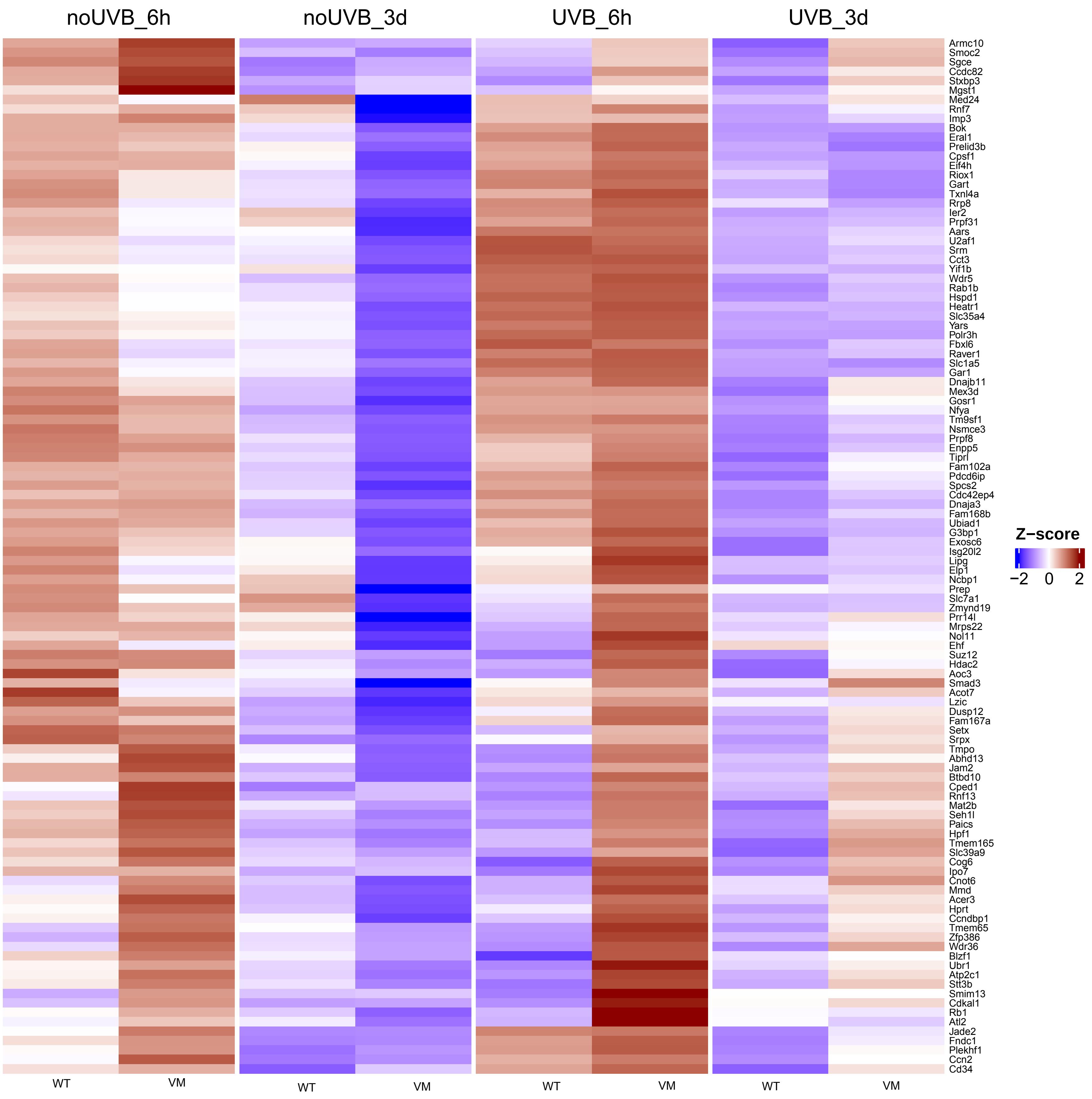

#### Cluster2 ... Genes 881 to 980

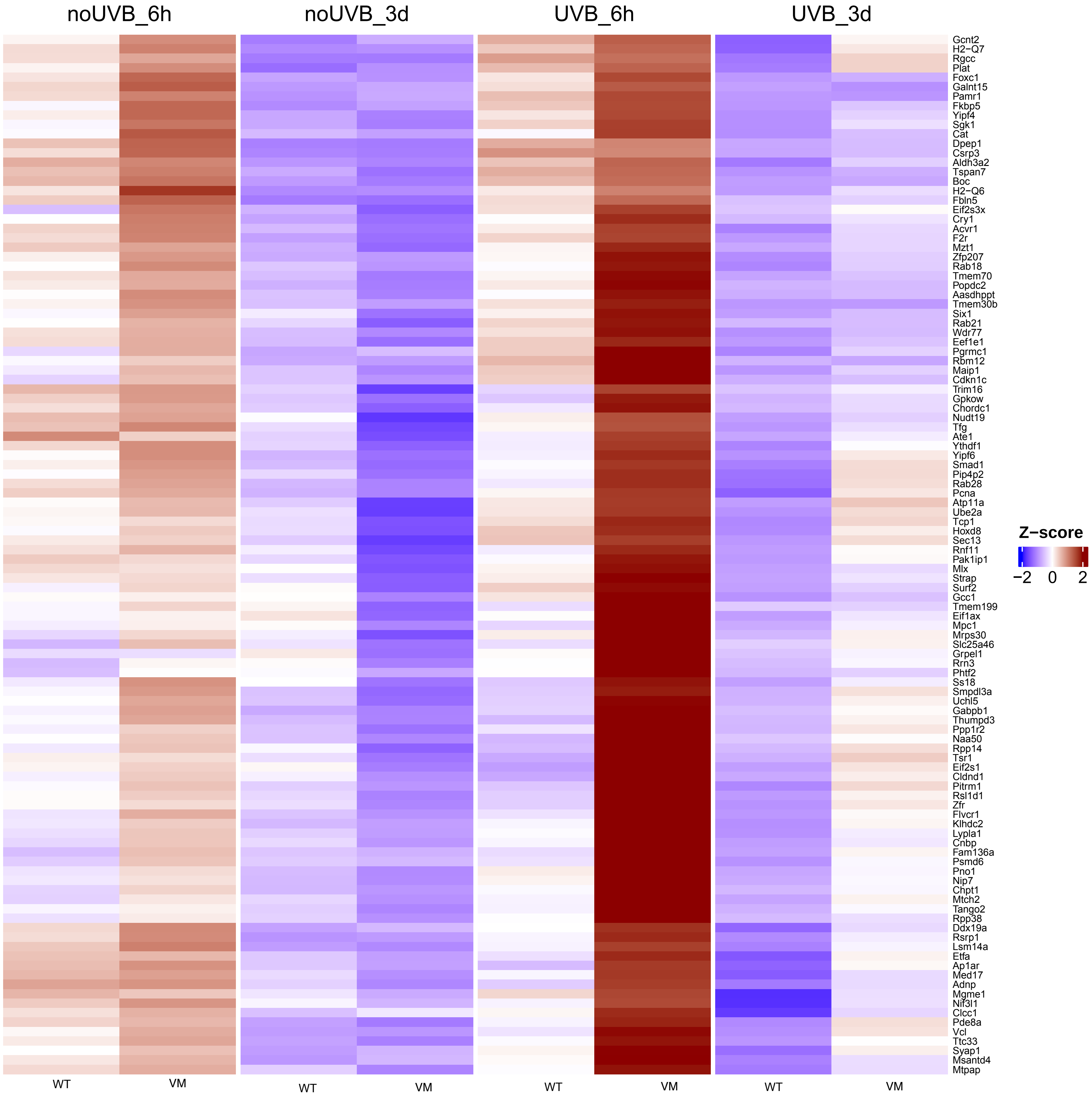

**Cluster2 ... Genes 991 to 1090**

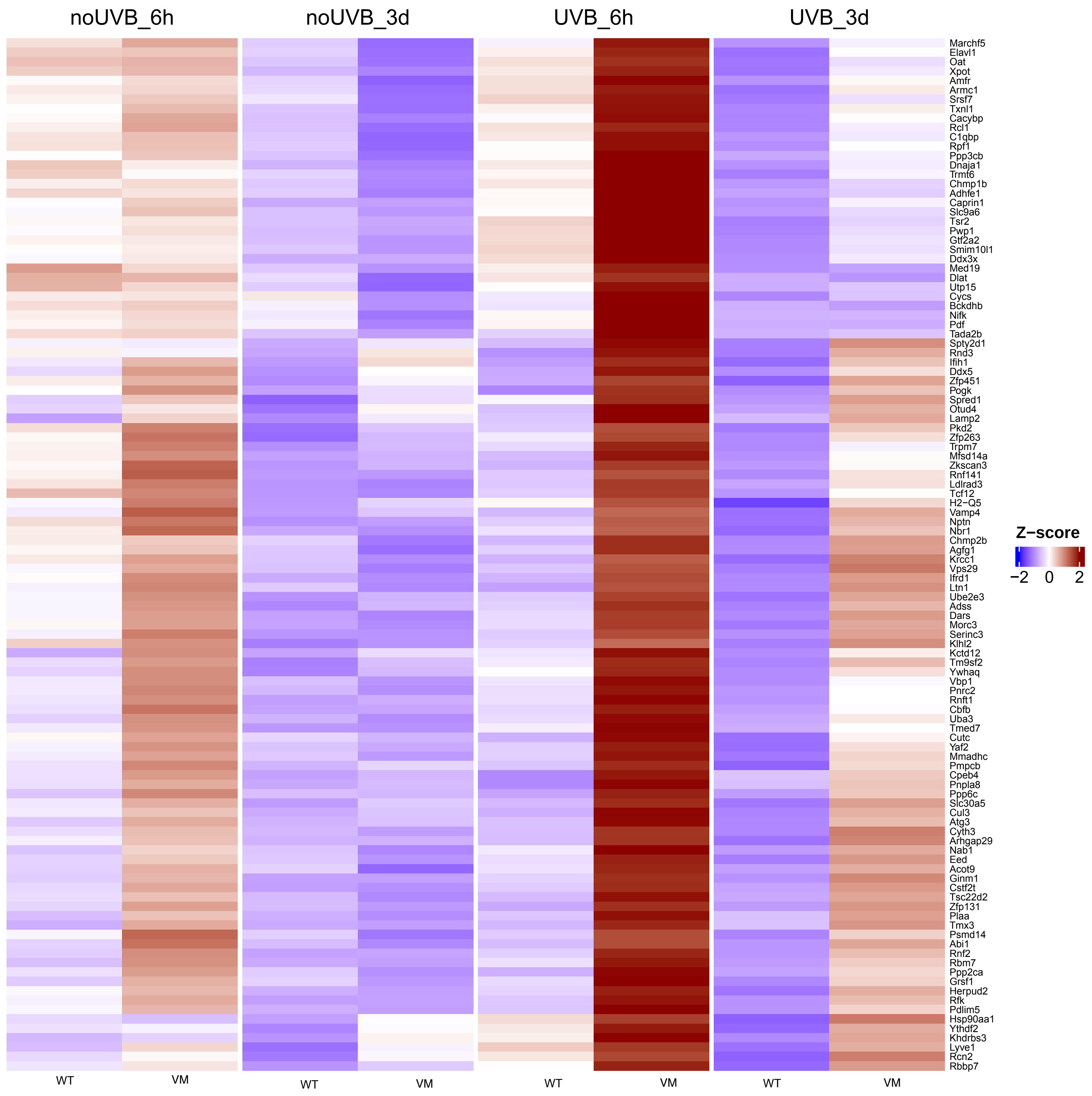

**Cluster2 ... Genes 1101 to 1200**

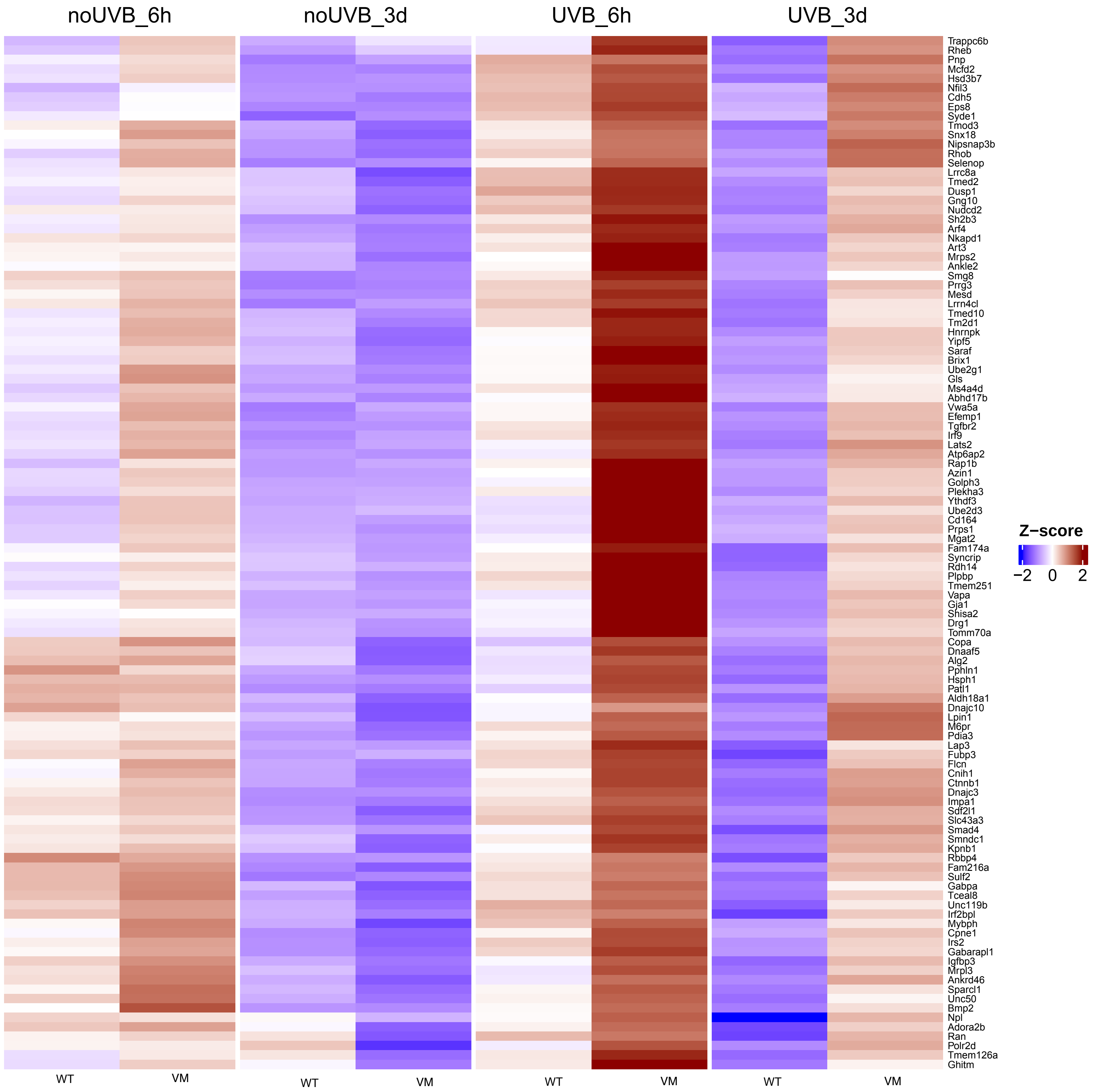

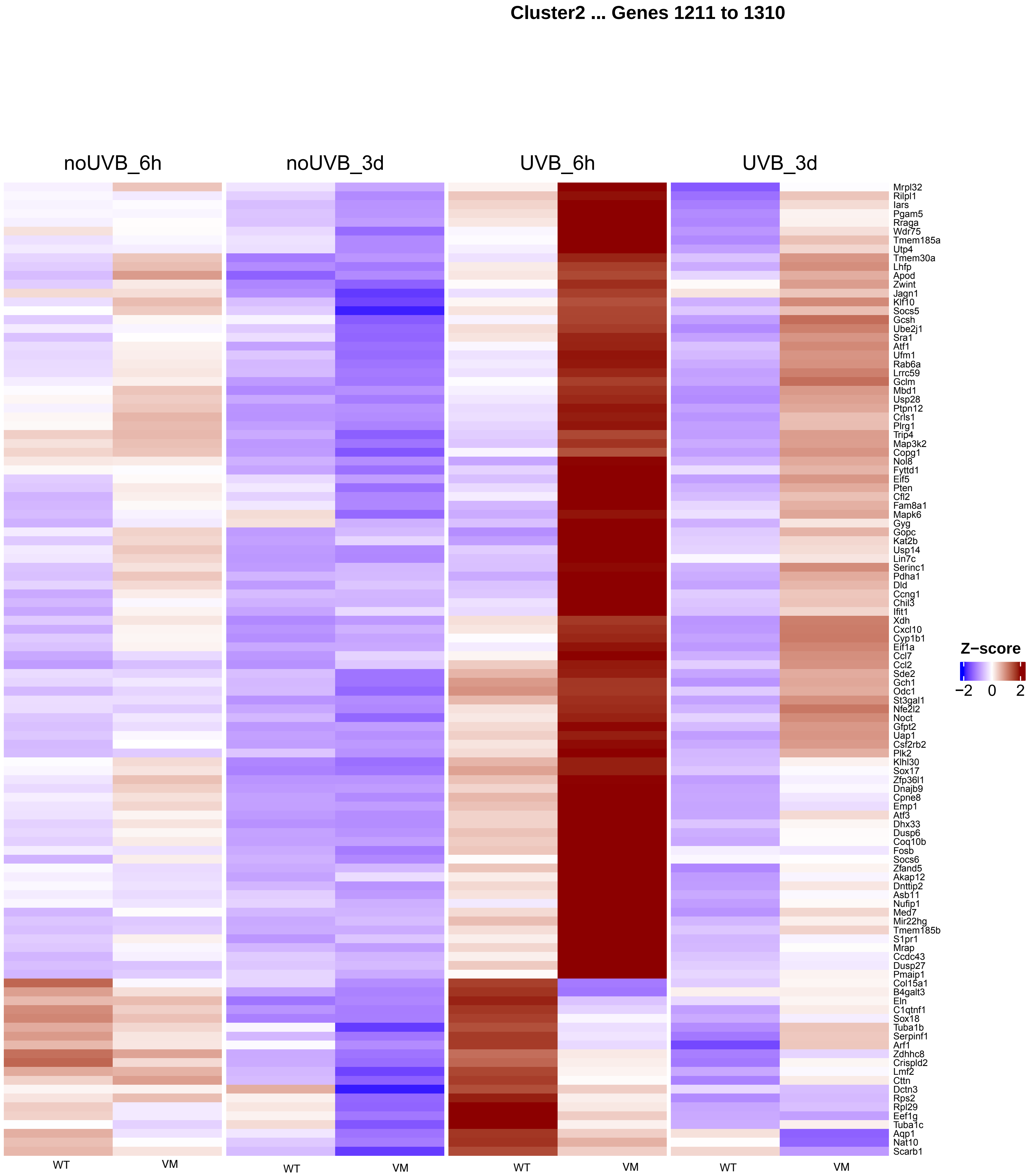
Cluster2 ... Genes 1211 to 1310
noUVB_6h
noUVB_3d
UVB_6h
UVB_3d
Mrpl32
Rilpl1
Iars
Pgam5
Rraga
Wdr75
Tmem185a
Utp4
Tmem30a
Lhfp
Apod
Zwint
Jagn1
Klf10
Socs5
Gcsh
Ube2j1
Sra1
Atf1
Ufm1
Rab6a
Lrrc59
Gclm
Mbd1
Usp28
Ptpn12
Crls1
Plrg1
Trip4
Map3k2
Copg1
Nol8
Fyttd1
Eif5
Pten
Cfl2
Fam8a1
Mapk6
Gyg
Gopc
Kat2b
Usp14
Lin7c
Serinc1
Pdha1
Dld
Ccng1
Chil3
Ifit1
Xdh
Cxcl10
Cyp1b1
Eif1a
Ccl7
Ccl2
Sde2
Gch1
Odc1
St3gal1
Nfe2l2
Noct
Gfpt2
Uap1
Csf2rb2
Plk2
Klhl30
Sox17
Zfp36l1
Dnajb9
Cpne8
Emp1
Atf3
Dhx33
Dusp6
Coq10b
Fosb
Socs6
Zfand5
Akap12
Dnttip2
Asb11
Nufip1
Med7
Mir22hg
Tmem185b
S1pr1
Mrap
Ccdc43
Dusp27
Pmaip1
Col15a1
B4galt3
Eln
C1qtnf1
Sox18
Tuba1b
Serpinf1
Arf1
Zdhhc8
Crispld2
Lmf2
Cttn
Dctn3
Rps2
Rpl29
Eef1g
Tuba1c
Aqp1
Nat10
Scarb1
WT
VM
WT
VM
WT
VM
WT
VM
Z−score
−2
0
2

### Cluster2 ... Genes 1321 to 1369

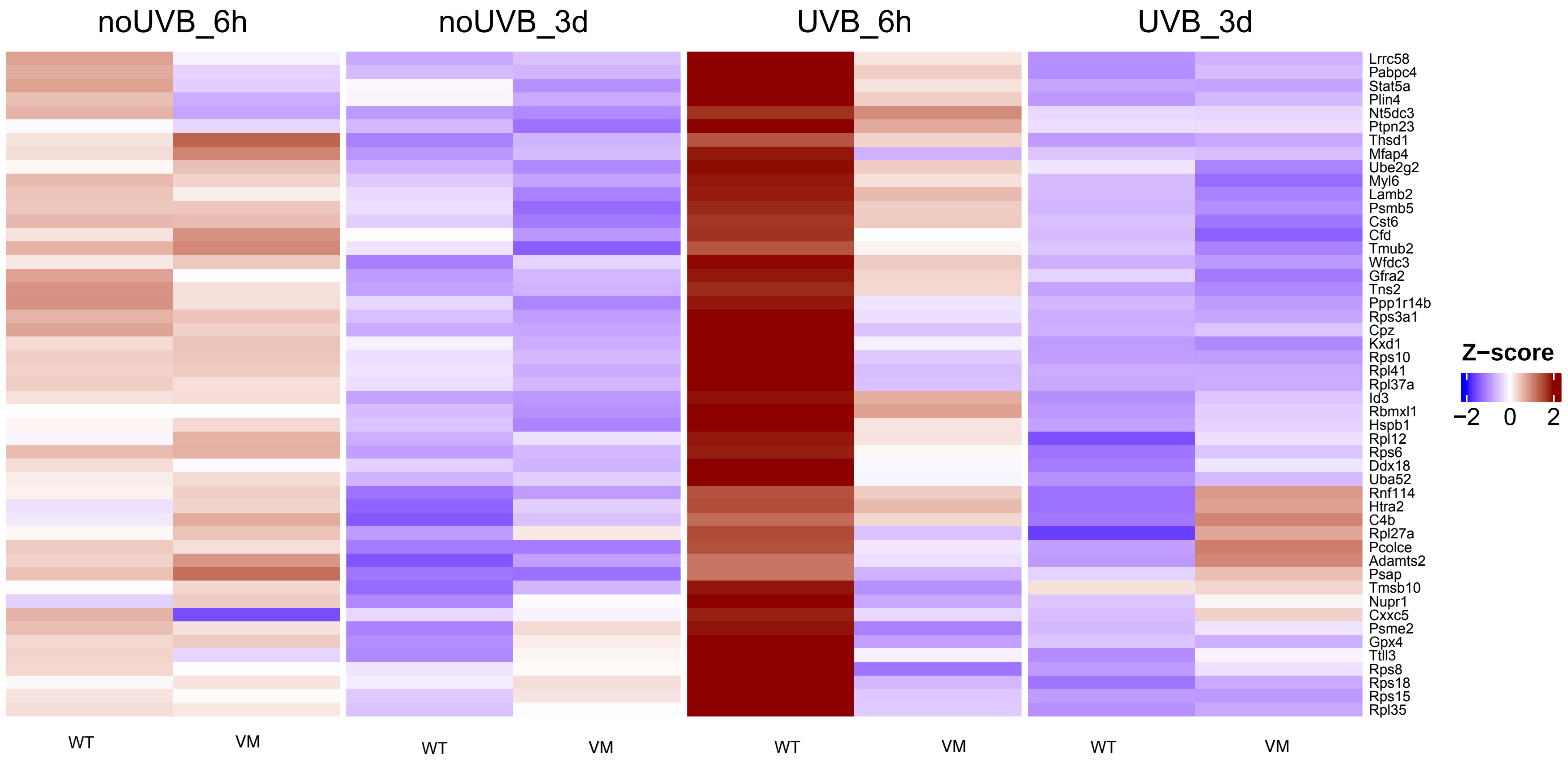
