## Supplemental Table 1 for "Endothelial Cell Expression of STING^V154M^ Gain-of-Function Mutation Delays the Resolution of UVB-induced Skin Injury"

| Color | Marker | Dilution | Clone | Brand |
|---|---|---|---|---|
| Spark UV 387 | CD11b | 1:200 | M1-70 | Biolegend |
| BV 805 | CD45 | 1:200 | 30-F11 | BD Bioscience |
| Pacific Blue | TCR beta chain | 1:200 | H57-597 | Biolegend |
| BV480 | CD11c | 1:200 | HL3 | Biolegend |
| BV510 | CD8b | 1:200 | YTS156.7.7 | Biolegend |
| BV650 | CD19 | 1:200 | 6D5 | Biolegend |
| BV711 | CD103 | 1:200 | 2E7 | Biolegend |
| BV785 | F4/80 | 1:100 | BM8 | Biolegend |
| FITC | CD69 | 1:200 | H1.2F3 | Biolegend |
| SparkBlue550 | MHC II | 1:800 | M5/114.15.2 | Biolegend |
| PerCP | Ly6C | 1:100 | HK1.4 | Biolegend |
| PE | TCR gamma delta | 1:200 | GL3 | Biolegend |
| PE Fire 700 | NK1.1 | 1:800 | S17016D | Biolegend |
| AF700 | CD4 | 1:200 | GK1.4 | Biolegend |
| Zombie NIR | live/dead | 1:1000 | -- | Biolegend |
| APC Fire 810 | Ly6G | 1:200 | 1A8 | Biolegend |

**Supplementary Table 1. Antibodies used for flow cytometry analysis.** List of fluorochrome-conjugated antibodies used for flow cytometric analysis of mouse skin-infiltrating immune cells. The table includes fluorophore, target antigen, dilution, clone, and vendor.
